# Identification of Paired-related Homeobox Protein 1 as a key mesenchymal transcription factor in Idiopathic Pulmonary Fibrosis

**DOI:** 10.1101/2021.01.15.425870

**Authors:** E. Marchal-Duval, M. Homps-Legrand, A. Froidure, M. Jaillet, M. Ghanem, L. Deneuville, A. Justet, A. Maurac, A. Vadel, E. Fortas, A. Cazes, A. Joannes, L. Giersch, H. Mal, P. Mordant, C.M. Mounier, K. Schirduan, M. Korfei, A. Gunther, B. Mari, F. Jaschinski, B. Crestani, A.A. Mailleux

**Affiliations:** Institut National de la Santé et de la Recherche Médical, UMR1152, Labex Inflamex, DHU FIRE, Université de Paris, Faculté de médecine Xavier Bichat, 75018 Paris, France; Institut de Recherche Expérimentale et Clinique, Pôle de Pneumologie, Université catholique de Louvain, Belgium Service de pneumologie, Cliniques Universitaires Saint-Luc, Brussels, Belgium; Assistance Publique des Hôpitaux de Paris, Hôpital Bichat, Service de Pneumologie A, DHU FIRE, Paris, France; Assistance Publique des Hôpitaux de Paris, Hôpital Bichat, Département d’Anatomopathologie, DHU FIRE, Paris, France; Univ Rennes, Inserm, EHESP, Irset (Institut de recherche en santé, environnement et travail) - UMR_S 1085, F-35000 Rennes, France; Assistance Publique des Hôpitaux de Paris, Hôpital Bichat, Service de Pneumologie et Transplantation, DHU FIRE, Paris, France; Assistance Publique des Hôpitaux de Paris, Hôpital Bichat, Service de Chirurgie Thoracique et Vasculaire, DHU FIRE, Paris, France; Université Côte d’Azur, CNRS, IPMC, FHU-OncoAge, Valbonne, France; CYU Université, ERRMECe(EA1391), NEUVILLE SUR OISE, France; Secarna Pharmaceuticals GmbH & Co. KG – Planegg/Martinsried (Germany); Department of Internal Medicine II, University of Giessen-Marburg Lung Center, Justus-Liebig University Giessen, Giessen, Germany

**Keywords:** lung fibrosis, IPF, transcription factor, mesenchyme, fibroblast, bleomycin

## Abstract

Matrix remodeling is a salient feature of idiopathic pulmonary fibrosis (IPF). Targeting cells driving matrix remodeling could be a promising avenue for IPF treatment. Analysis of transcriptomic database identified the mesenchymal transcription factor PRRX1 as upregulated in IPF.

PRRX1, strongly expressed by lung fibroblasts, was regulated by a TGF-β/PGE2 balance in vitro in control and IPF fibroblasts, while IPF fibroblast-derived matrix increased *PRRX1* expression in a PDGFR dependent manner in control ones.

PRRX1 inhibition decreased fibroblast proliferation by downregulating the expression of S phase cyclins. PRRX1 inhibition also impacted TGF-β driven myofibroblastic differentiation by inhibiting SMAD2/3 phosphorylation through phosphatase PPM1A upregulation and TGFBR2 downregulation, leading to TGF-β response global decrease.

Finally, targeted inhibition of *Prrx1* attenuated fibrotic remodeling in vivo with intra-tracheal antisense oligonucleotides in bleomycin mouse model of lung fibrosis and ex vivo using precision-cut lung slices.

Our results identified PRRX1 as a mesenchymal transcription factor driving lung fibrogenesis.

**Brief Summary:** Inhibition of a single fibroblast-associated transcription factor, namely paired-related homeobox protein 1, is sufficient to dampen lung fibrogenesis.

## INTRODUCTION

Chronic remodeling is a key feature of many Human diseases associated with aging. In particular, chronic respiratory diseases, including lung fibrosis, are a major and increasing burden in terms of morbidity and mortality ^1^. For instance, idiopathic pulmonary fibrosis (IPF) is the most common form of pulmonary fibrosis. IPF is defined as a specific form of chronic, progressive fibrosing interstitial pneumonia of unknown cause. IPF patients have an overall median survival of 3 to 5 years ^1^.

According to the current paradigm, IPF results from progressive alterations of alveolar epithelial cells leading to the recruitment of mesenchymal cells to the alveolar regions of the lung with secondary deposition of extracellular matrix, and destruction of the normal lung structure and physiology. IPF develops in a susceptible individual and is promoted by interaction with environmental agents such as inhaled particles, tobacco smoke, inhaled pollutants, viral and bacterial agents. Aging is probably both a susceptibility marker and a major driver of the disease, through mechanisms that are not yet fully elucidated. Two drugs (Pirfenidone and Nintedanib) appear to slow disease progression and may improve long term survival ^1^.

In any given cell, a set of transcription factors is expressed and works in concert to govern cellular homeostasis and function. Dysregulation of transcriptional networks may therefore account to aberrant phenotypic changes observed in lung fibrosis. Among the multifaceted tissue cellular “ecosystem”, cells of mesenchymal origin such as fibroblasts are the main cellular components responsible for tissue remodeling during normal and pathological lung tissue repair ^1,2^. Thus, targeting master transcription factors preferentially expressed in fibroblast could be a promising avenue for IPF treatment.

Using an in silico approach, we screened publicly available transcript microarray databases for expression of mesenchyme-associated transcription factors in control and IPF lung samples. We identified the “Paired Related Homeobox Protein-1” (*PRRX1)* gene as a potential candidate for transcriptional regulation differently modulated in IPF compared to control lungs. The *PRRX1* mRNA generates by alternative splicing two proteins, PRRX1a (216 aa) and PRRX1b (245 aa) that differ at their C-terminal parts. Functional in vitro studies suggested that PRRX1a promoted transcriptional activation whereas PRRX1b may act rather as a transcriptional repressor ^3^.

*Prrx1* is implicated in the regulation of mesenchymal cell fate during embryonic development. PRRX1 is essential for fetal development as *Prrx1*^*-/-*^ mice present severe malformation of craniofacial, limb, and vertebral skeletal structures ^4^. *Prrx1*^*-/-*^ mice also display hypoplastic lungs with severe vascularization defects and die soon after birth ^5^. PRRX1 function is not restricted to embryogenesis. It has been also shown that PRRX1 was a stemness regulator ^6^, involved in adipocyte differentiation ^7^, epithelial tumor metastasis and pancreatic regeneration ^8–10^ as well as liver fibrosis ^11^. PRRX1 transcription factors are also at the center of the network coordinating dermal fibroblast differentiation ^12^.

However, whether and how PRRX1 plays a role in lung fibrogenesis still remains elusive. Given its central position in fibroblast transcriptional network, we hypothesized that PRRX1 transcription factors are important drivers of the fibroblast phenotype in IPF, promoting the development/progression of fibrosis.

## RESULTS

### Identification of *PRRX1* isoforms as mesenchymal transcription factors associated with IPF

Since mesenchymal cells are thought to be one of the major effector cells during fibrosis ^1,2^, we sought to identify mesenchymal transcription factors associated with IPF in patients. We screened three curated publicly available transcript microarray databases from NCBI GEO (GDS1252, GDS4279, GDS3951) for transcription factor expression in IPF and control whole lung samples. Among the 210 common genes upregulated at the mRNA level in all three IPF lung datasets compared to their respective control ones (Figure 1a and supplemental Table S1), 12 genes were annotated as transcription factors (Figure 1a) after gene ontology analysis. One of these transcription factors, *PRRX1* appeared as an appealing candidate since this gene was previously associated with mesenchymal cell fate during embryogenesis ^4^ and is required for proper lung development ^5^. In addition, *PRRX1* mRNA was upregulated in a fourth transcriptome dataset comparing “rapid” and “slow” progressor subgroups of IPF patients ^13^. None of those transcriptome datasets discriminated *PRRX1* isoforms, namely *PRRX1a* and *PRRX1b*.

**Figure 1:**
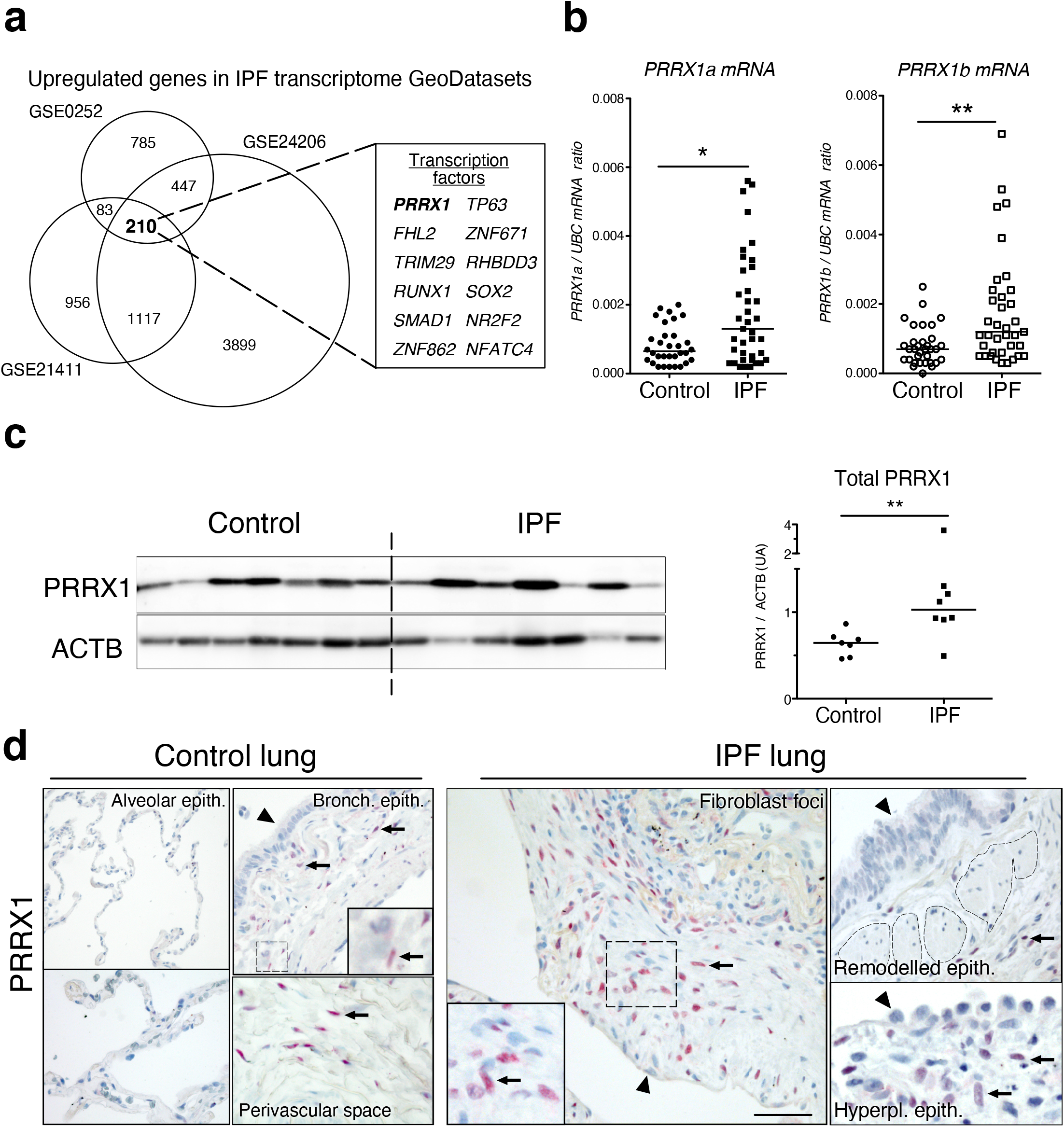
Identification of PRRX1 as a transcription factor reactivated in IPF lung. (**a**) Venn diagram showing the number of genes up-regulated in three IPF lung Transcript microarray databases compared to controls (NCBI GEO GDS1252, GDS4279, GDS3951). Among the 210 common upregulated genes in all three datasets, 12 genes were annotated as transcription factors (table, PRRX1 is in bold). (**b**) Dot plots with median showing the mRNA expression of *PRRX1a* and *PRRX1b* isoforms in control (circle, n=35) and IPF (square, n=38) whole lung homogenates. (**c**) Immunoblot showing PRRX1 expression in control and IPF whole lung homogenates. ACTB was used as loading control. The quantification of PRRX1 relative expression to ACTB in control (circle, n=7) and IPF (square, n=8) is displayed as dot plot with median on the right. (**d**) Representative immunohistochemistry images (n=5 per group) showing PRRX1 staining (red) in control (left panels) and IPF (right panels). Nuclei were counterstained with hematoxylin. Note the absence of PRRX1 staining in the alveolar and bronchiolar epithelium (arrow head). PRRX1 positive cells were only detected in the peri-bronchiolar and peri-vascular spaces (arrows) in control lungs (left panels). In IPF, PRRX1 positive cells (arrow) were detected in the remodeled/fibrotic area (right panels). Note that epithelial cells (arrow head) and bronchiolar smooth muscle cells (dashed areas) are PRRX1 negative. The high magnification pictures match the dashed boxes displayed in the main panels. Scale bar: 80µm in low magnification images and 25µm in high magnification ones. *Abbre*via*tions: epithelium. (epith); bronchiolar. (bronch); Hyperpl. (hyperplastic)*. Mann Whitney U test, *p≤0.05, **p≤0.01.

First, we confirmed that both *PRRX1* isoforms were upregulated in Human IPF lungs at the mRNA and protein levels (Figure 1a-c). Immunoblot revealed that PRRX1a protein (210 aa) was the main PRRX1 isoform expressed in control and IPF lungs. We also investigated PRRX1 expression pattern by immunohistochemistry in control and IPF Human lung tissue sections (the antibody recognized both PRRX1 isoforms, see Figure 1d). Additional lineage markers were also investigated such as Vimentin (mesenchyme marker), ACTA2 (myofibroblast / smooth muscle marker) and CD45 (hematopoietic lineage) as shown in supplemental Figure S1. PRRX1 positive cells were not detected in the distal alveolar space and in the bronchiolar epithelium of control lung (Figure 1d). Nevertheless, PRRX1 nuclear staining was observed in mesenchymal cells (Vimentin positive but ACTA2 and CD45 negative cells) within peri-vascular and peri-bronchiolar spaces (Figure 1d and supplemental Figure S1). In IPF patients, nuclear PRRX1 positive cells were mainly detected in the fibroblast foci (Figure 1d), which are the active sites of fibrogenesis ^1,2^. Those PRRX1 positive cells were all Vimentine positive and CD45 negative but only some were expressing ACTA2 (supplemental Figure S1).

To confirm the identity of PRRX1 expressing cells in the lung as fibroblasts, we took advantage of recently published single-cell transcriptomic analysis performed using lung samples ^14,15^. *PRRX1* mRNA expression was restricted to the fibroblast / mesenchymal cell lineages in either lung transplant donors or recipients with pulmonary fibrosis (Figure 2a and supplemental Figure S2).

**Figure 2:**
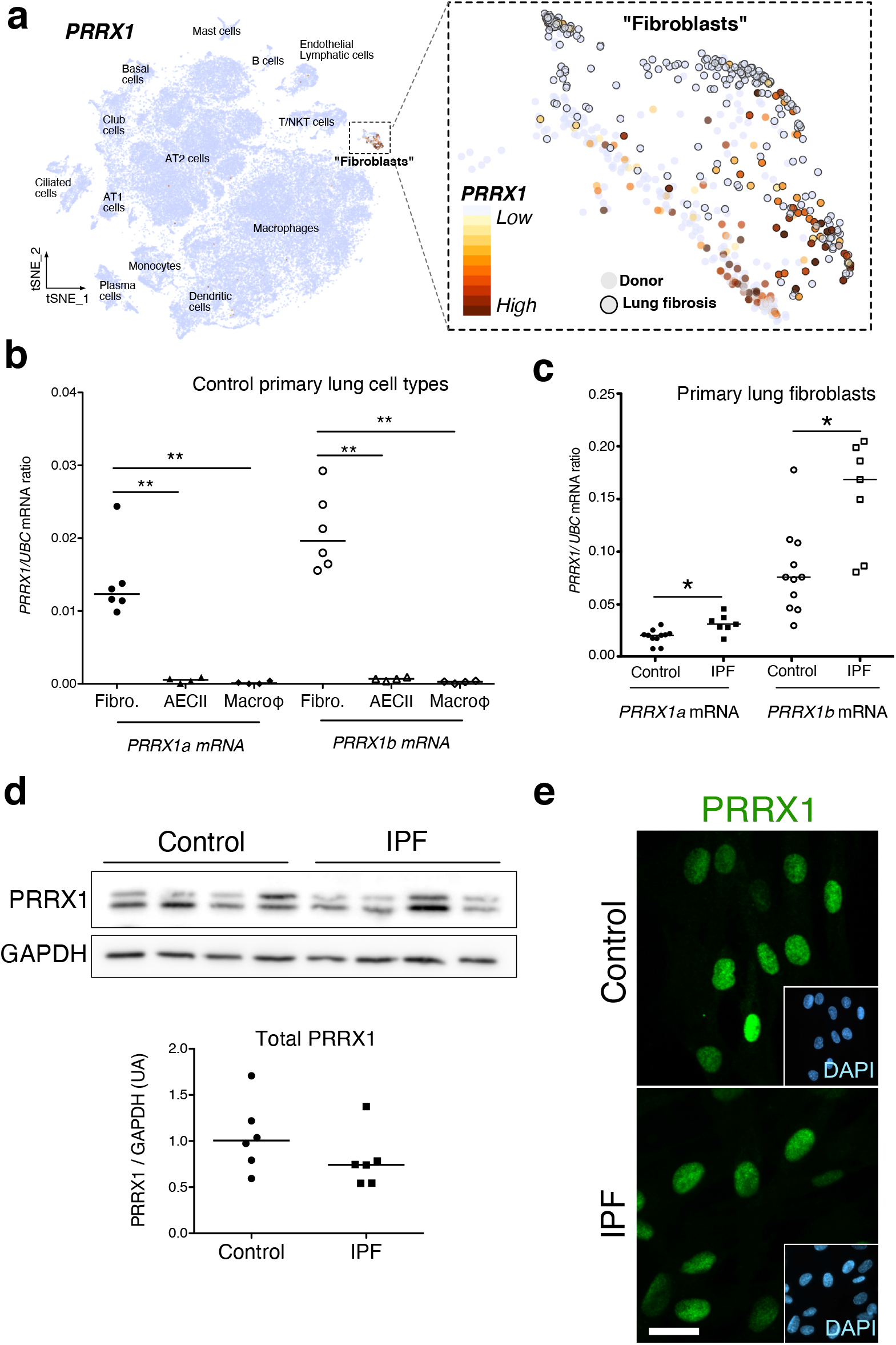
PRRX1 is a mesenchymal transcription factor upregulated in primary Human lung IPF fibroblasts. (**a**) Integrated single-cell RNA-Seq analysis of donors as well as patients with pulmonary fibrosis showing diverse lung cell populations using previously published data from ^14^ (source: www.nupulmonary.org/resources/). *PRRX1* mRNA expression was used to label clusters by cell identity as represented in the tSNE plot. Note that *PRRX1* mRNA expression is restricted to cell types classified as “Fibroblasts”. (**b**) Dot plots with median showing the mRNA expression of *PRRX1a* (black) and *PRRX1b* (white) isoforms in primary Human lung fibroblasts (circle, n=6), alveolar epithelial cells (triangle, n=4) and alveolar macrophages (diamond, n=4). (**c**) Dot plots with median showing the mRNA expression of *PRRX1a* (black) and *PRRX1b* (white) isoforms in control (circle, n=11) and IPF (square, n=7) primary Human lung fibroblasts. (**d**) Immunoblot showing PRRX1 expression in control and IPF primary Human lung fibroblasts. GAPDH was used as loading control. The quantification of PRRX1 relative expression to GAPDH in control (circle, n=6) and IPF (square, n=6) lung fibroblasts is displayed as dot plot with median below. (**e**) Representative Immunofluorescence images (n= 8 per group) showing PRRX1 staining (green) in control (top panel) and IPF (bottom panel) fibroblasts. Nuclei were counterstained with DAPI (inserts in main panels). Scale bar 20µm in main panels and 40µm in inserts. *Abbreviations: fibroblasts (fibro); alveolar epithelial cells (AECII); alveolar macrophages (macroϕ)*. Mann Whitney U test, *p≤0.05.

In vitro, both isoforms, *PRRX1a* and *PRRX1b* mRNA were also found to be strongly expressed by primary lung fibroblasts compared to primary alveolar epithelial type 2 cells (AECII) and alveolar macrophages (Figure 2c) by quantitative PCR (qPCR). In addition, *PRRX1a* and *-1b* levels were increased in IPF primary lung fibroblasts compared to control ones only at the mRNA level as assayed by qPCR and western blot (Figure 2c-d). By immunofluorescence, PRRX1 was detected at the protein level in the nuclei of both control and IPF fibroblasts cultured in vitro (Figure 2e).

### *PRRX1* isoforms expression in primary lung fibroblasts is tightly regulated by growth factors and extracellular matrix stiffness *in vitro*

To better understand the regulation of PRRX1 isoforms in lung fibroblasts, we first assayed the effects of factors known to regulate lung fibroblast to myofibroblasts differentiation on the expression of both PRRX1 isoforms in control and IPF primary lung fibroblasts. “Transforming growth factor beta 1” (TGF-β1, 1ng/ml) treatment which triggers myofibroblastic differentiation^2^ was associated with a decrease in the expression level of both *PRRX1* isoforms at the mRNA level (Figure 3a). This effect was confirmed at the protein level by western blot and immunofluorescence (supplemental Figure S3). Conversely, prostaglandin E_2_ (PGE_2_, 100nM) treatment which decreases myofibroblastic differentiation^2^ was associated with an increase of *PRRX1* isoforms mRNA and protein in both control and IPF fibroblasts (Figure 3a and supplemental Figure S3). These data indicate that PRRX1 isoform expression is controlled by a TGF-β/PGE2 balance in lung fibroblasts.

**Figure 3:**
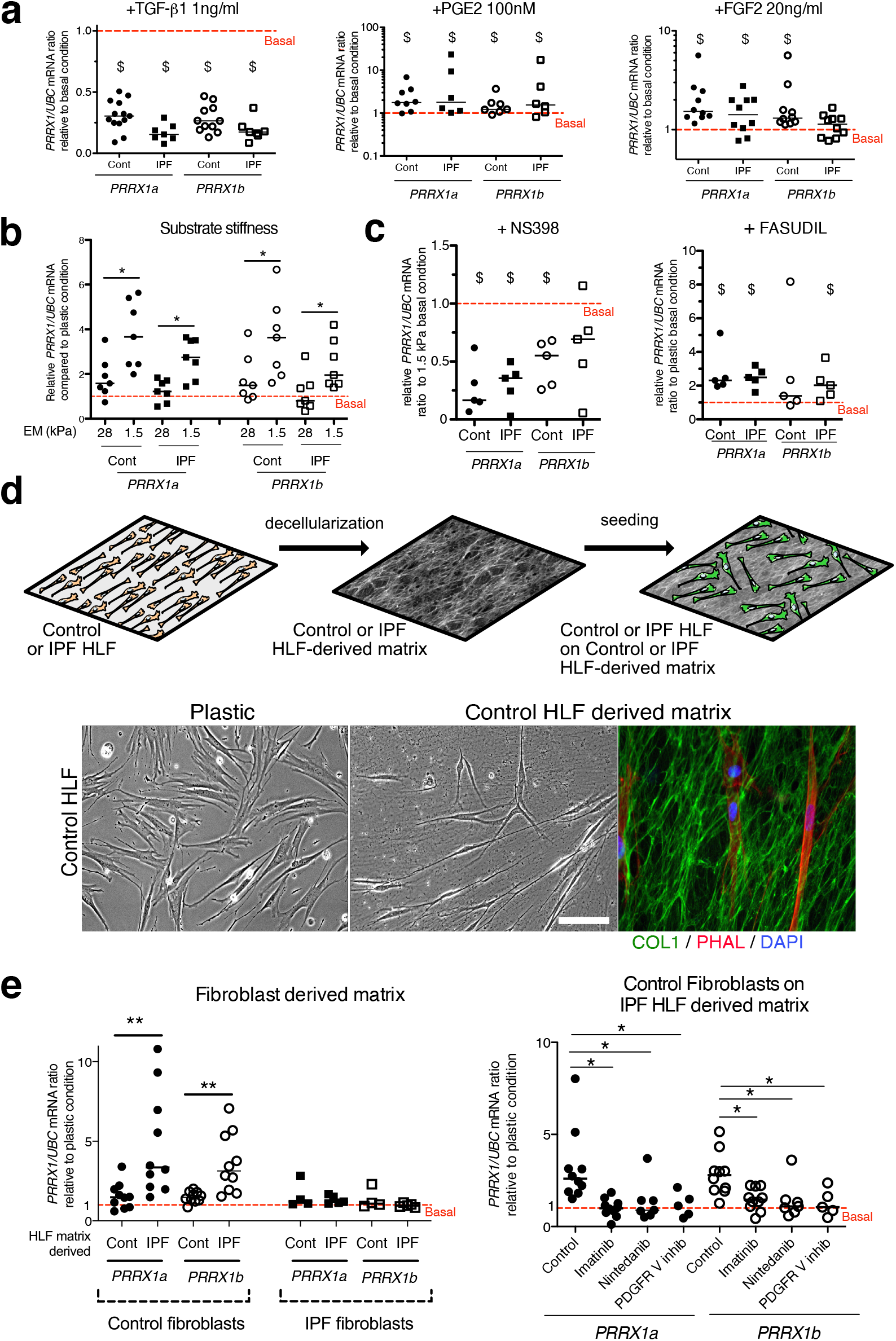
PRRX1 is modulated by growth factors and matrix environment. (**a)** Dot plots with median showing the mRNA expression of *PRRX1a* (black) and *PRRX1b* (white) isoforms in control (circle) and IPF (square) primary Human lung fibroblasts stimulated for 48h with TGF-β1 (left, n=6-7), PGE2 (middle, n=6-7) and FGF2 (right, n=10) compared to basal condition (red dashed line). (**b**) dot plots with median showing the mRNA expression of *PRRX1a* and *PRRX1b* isoforms in control (n=7) and IPF (n=7) lung fibroblasts cultured on stiff (28kPa) and soft (1.5kPa) substrate compared to basal condition. (**c**) Dot plots with median showing the mRNA expression of *PRRX1a* and *PRRX1b* isoforms in control and IPF lung fibroblasts (n=5) stimulated 48h with NS398 (left), or Fasudil (right) compared to basal condition. (**d**) Summary sketch of Fibroblast-derived matrix experiments (upper part). Lower part: representative phase contrast pictures of control primary lung fibroblasts on plastic (left) or seeded in a control fibroblast-derived matrix (middle) and immunofluorescence pictures (right) of Collagen 1 (green) revealing the HLF-derived matrix and actin fibers stained with Phalloidin (red). Nuclei were counterstained with DAPI (blue). (**e**) Left panel: dot plots with median showing the mRNA expression of *PRRX1a* and *PRRX1b* isoforms in control (n=10) or IPF fibroblasts (n=5) seeded on control or IPF derived matrix compared to basal condition. Right panel: dot plots with median showing the mRNA expression of *PRRX1a* and *PRRX1b* isoforms in control fibroblasts cultured on IPF derived matrix and stimulated with Imatinib (n=10), Nintedanib (n=7) or PDGFR V inhibitor (n=5) compared to basal condition. (Scale bar: 30µm in phase contrast pictures and 15µm in the immunofluorescence one) *Abbreviations: Control (Cont), Human lung fibroblast (HLF), Elastic/Young modulus (EM), PHAL (Phalloidin), COL1 (Collagen 1)*. Mann Whitney U test, *p≤0.05 **p≤0.01; Wilcoxon signed-rank test $ p≤0.05.

Concomitantly with an aberrant growth factors/chemokine secretory profile, lung fibrosis is also characterized by local matrix stiffening, which plays a key role in IPF physiopathology ^16^. Previous studies showed that increasing matrix stiffness strongly suppressed fibroblast expression of *PTGS2* ^17^, a key enzyme in PGE_2_ synthesis, and increased Rho kinase (ROCK) activity ^18^, contributing to myofibroblastic differentiation. Control and IPF primary lung fibroblasts were cultured on fibronectin-coated glass (elastic/Young’s modulo in the GPa range) or hydrogel substrates of discrete stiffness, spanning the range of normal (1.5kPa) and fibrotic (28kPa) lung tissue ^16^. We confirmed that soft (1.5kPa) substrate culture condition did increase *PTGS2* mRNA level compared to stiff/glass control condition in both control and IPF fibroblasts (data not shown). The expression levels of both *PRRX1* TFs isoforms mRNA were also increased on soft/normal 1.5kPa stiffness substrate (Figure 3b) compared to stiff substrates (Glass and 28kPa culture conditions). Treatment with NS398 (10µg/ml), a specific PTGS2 inhibitor abrogated the *PRRX1* TFs increase on soft substrate (Figure 3c). Conversely, inhibition of mechanosensitive signalling with Fasudil (35µM), an inhibitor of ROCK1 and ROCK2 ^18^, induced *PRRX1* TFs mRNA expression in both control and IPF fibroblasts grown on glass/stiff substrate (Figure 3c). Collectively, these data indicate that PRRX1 expression is tightly controlled by extra-cellular matrix stiffness through a PTGS2/ROCK activity balance.

### IPF fibroblast-derived matrix increased PRRX1 expression in control fibroblasts in a PDGFR dependent manner

In order to better appreciate the regulation of PRRX1 expression in a complex environment, we cultured lung fibroblasts in a fibroblast-derived 3D ECM. Control and IPF fibroblasts were maintained in high-density culture to generate thick matrices that were extracted with detergent at alkaline pH to remove cellular contents (Figure 3d). This treatment leaves behind a 3D ECM that is intact and cell-free ^19^.

We observed that *PRRX1a* and -*1b* TF mRNA expression was upregulated in control fibroblasts cultured on IPF fibroblast derived 3D ECM compared to plastic culture (Figure 3e). Meanwhile, *PRRX1a* and *-1b* mRNA expression levels were stable in IPF fibroblasts seeded either on control or IPF fibroblast derived 3D ECM compared to plastic culture (Figure 3e).

To better understand the cellular processes and signalling pathways involved, control fibroblasts seeded on IPF fibroblast-derived matrix were treated with two tyrosine kinase protein inhibitors, namely Imatinib (10µg/ml) and Nintedanib (10nM). Those tyrosine kinase inhibitors have anti-fibrotic properties on lung fibroblasts ^20^ and Nintedanib is one of the two drugs currently approved for IPF treatment^1^. Both inhibitors reverted the effect of IPF fibroblast-derived matrix upon *PRRX1a* and *PRRX1b* mRNA levels (Figure 3e). Interestingly, amongst their multiple targets ^1,2^, Imatinib and Nintedanib are known to both inhibit PDGFR. Treatment with a specific PDGFR inhibitor (PDGFR V ^21^) at the nanomolar range (10nM) did confirm the PDGFR-dependency of this effect of IPF fibroblast-derived ECM on *PRRX1a* and -*1b* mRNA levels in control fibroblasts (Figure 3e).

### PRRX1 TF isoforms promote fibroblast proliferation

Since PRRX1 expression is strongly associated with fibroblasts in IPF, we next investigated whether PRRX1 TFs may drive the phenotype of primary lung fibroblasts. The involvement of PRRX1 was studied in vitro by using siRNA targeting both *PRRX1a* and *PRRX1b* isoforms (loss of function).

First, we investigated the effects of *PRRX1* TFs knock down using two different siRNA sequences (see Figure 4a-b). Knockdown of *PRRX1* TFs significantly decreased primary lung fibroblast proliferation in complete growth medium after 72h (Figure 4c) as compared to the control siRNA. Cell cycle analysis revealed a significant decrease in S phase concomitantly with an increase in G1 phase, suggestive of a G1/S arrest in control and IPF lung fibroblasts treated with *PRRX1* siRNA (Figure 4d and supplemental Figure S4). This potential G1/S arrest was also associated with a strong decrease in *CCNA2* and *CCNE2* mRNA expression after 72 hours (Figure 4e). These two cyclins play a key role in the replicative S phase during the cell cycle ^22^. To further characterize the impact of *PRRX1* TF inhibition on cell cycle progression, we performed a FACS analysis of KI67 expression in primary control and IPF lung fibroblasts transfected with *PRRX1* siRNA sequences for 72h compared to control siRNA. KI67 protein is usually present during all active phases of the cell cycle, but is absent from resting cells in G0 ^23^. *PRRX1* inhibition strongly decreased the number of KI67 (MKI67; official name) positive cells in control and IPF lung fibroblasts (Figure 4f and supplemental Figure S4). Of note, *MKI67* expression was also decreased at the mRNA level in control and IPF lung fibroblasts treated with *PRRX1* siRNA (supplemental Figure S4). Next, we used a chromatin immunoprecipitation approach (ChIP) to assay a possible direct regulatory effect of PRRX1 upon *CCNA2, CCNE2* and *MKI67* gene loci in primary normal Human lung fibroblasts (NHLF). We observed an enrichment of PRRX1 binding at the vicinity of PRRX1 response element *(PRE)* identified in the *CCNA2, CCNE2* and *MKI67* promoter regions, suggesting that those genes could be direct PRRX1 TFs target genes. Meanwhile, no PRRX1 binding was detected at the *GAPDH* transcription starting site (TSS); devoid of PRE (Figure 4g).

**Figure 4:**
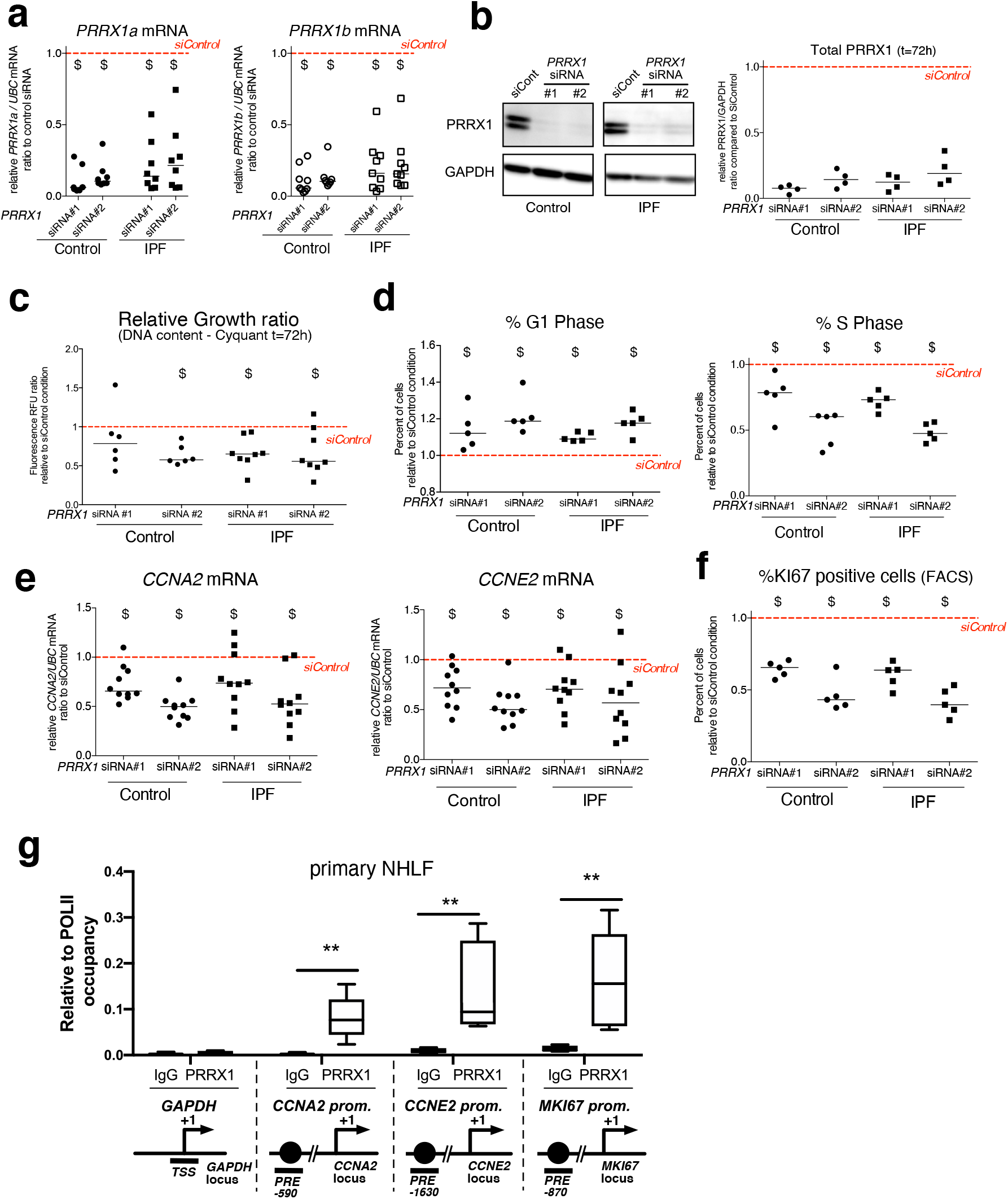
*PRRX1* knock down decreased cell proliferation. (**a**) Dot plots with median showing *PRRX1a* (black) and *PRRX1b* (white) mRNA expression relative to the siControl condition (red dashed line) in control (circle) and IPF (square) fibroblasts (n=8) treated for 48h with *PRRX1* siRNA (#1 or #2). (**b**) Immunoblot showing PRRX1 expression (n=4) in control and IPF fibroblasts treated 48h with *PRRX1* siRNA (#1 or #2) or siControl. The quantification of PRRX1 expression relative to GAPDH (loading control) is displayed as dot plot with median. (**c**) Dot plots with median showing the relative growth ratio of control (n=6) and IPF (n=8) fibroblasts stimulated 72h with FCS 10% and treated with *PRRX1* siRNA compared to siControl. (**d**) Dot plots with median showing the percent of cells in G1 (right) or S (left) phase in control and IPF fibroblasts (n=5) stimulated 72h with FCS and *PRRX1* siRNA relative to siControl. (**e**) Dot plots with median showing mRNA expression of *CCNA2* and *CCNE2* relative to siControl in control and IPF fibroblasts stimulated 72h with FCS and treated with *PRRX1* siRNA (n=10). (**f**) Dot plots with median showing the percent of cells positive for Ki67 marker in control and IPF fibroblasts stimulated 72h with FCS 10% and treated with *PRRX1* siRNA relative to siControl (n=5). (**g**) ChIP analysis for PRRX1 recruitment at the promoter of *GAPDH, CCNA2, CCNE2* and *MKI67* in NHLF (n=5) relative to RNA POL-II occupancy, displayed as boxes with median and min to max. The diagrams of the different loci are showing the PRRX1 response element position relative to the TSS. The PCR amplified regions are underscored. *Abbreviations: FCS (fetal calf serum); TSS (transcription starting site); IgG (Immunoglobulin); PRE (PRRX1 responses element); SRE (SRF response element), control siRNA sequence (siControl)*. Wilcoxon signed-rank test, $ p≤0.05, Wilcoxon matched-paired signed rank test ** p<0.01.

We also assayed the effect of *PRRX1* knock down on the mRNA expression of *CDKN2A* (p16), *CDKN1A* (p21) and *TP53*, major negative regulators of cell cycle also associated with cellular senescence ^22^. The expression of all three cell cycle inhibitors was increased only at the mRNA level in both control and IPF lung fibroblasts treated with *PRRX1* siRNA compared to control siRNA as assayed by qPCR and western blot (see supplemental Figure S4 and data not shown). This cell cycle arrest in control and IPF lung fibroblasts treated with *PRRX1* siRNA was not associated with an increase in β-Galactosidase activity, a senescence marker, compared to cells transfected with control siRNA (data not shown).

In conclusion, our results showed that PRRX1 controlled fibroblast proliferation in vitro.

### PRRX1 TFs are required for the induction of alpha smooth muscle actin during TGF-β1-driven myofibroblastic differentiation

Next, we investigated the effects of PRRX1 TFs partial loss of function on myofibroblastic differentiation in primary control and IPF lung fibroblasts. In an appropriate microenvironment, fibroblasts can differentiate by acquiring contractile properties (such as expression of alpha smooth muscle actin (ACTA2); gamma smooth muscle actin (*ACTG2*) and Transgelin (*TAGLN* / *SM22)* and becoming active producers of extracellular matrix (ECM) proteins (such as Collagen 1, COL1; Fibronectin, FN1; Tenascin C, TNC and Elastin, ELN). Aberrant activation of fibroblasts into myofibroblasts is thought to be a major driver of lung fibrogenesis ^1,2^.

We evaluated the effects of PRRX1 modulation on the expression of myofibroblast markers such as ACTA2, COL1 and FN1 at basal condition. PRRX1 TF loss of functions did not robustly modify the basal expression of these markers at the mRNA and proteins levels after 48h of treatment as assayed respectively by qPCR and western blot (supplemental Figure S5). Recently, PRRX1 has been implicated in a positive feed-back loop in which TWIST1 directly increased PRRX1 which subsequently induced Tenascin-C that itself stimulated TWIST1 activity in Cancer associated fibroblast (CAF), and in dermal and fetal Human lung fibroblast lines ^24^. However, we observed that *PRRX1* inhibition with siRNA failed to modulate *TNC* and *TWIST1* mRNA levels in adult primary control and IPF lung fibroblasts at basal condition (supplemental Figure S5).

As mentioned before, TGF-β1 is a major regulator of myofibroblastic differentiation. We determined whether PRRX1 TFs may regulate myofibroblastic differentiation upon TGF-β1 stimulation. Control and IPF primary lung fibroblasts were first treated with *PRRX1* siRNA for 48h and then stimulated with 1ng/ml TGF-β1 for 48h. The inhibition of *PRRX1* TFs impacted the upregulation of contractile-associated actin isoforms at the mRNA levels such as *ACTA2* (*α-SMA*) and *ACTG2* (*γ-SMA*) in response to TGF-β1 stimulation (Figure 5a and supplemental Figure S6) while the expression of the actin binding protein *TAGLN* (*SM22*) was not perturbed (supplemental Figure S6). The effect of PRRX1 inhibition upon ACTA2 upregulation was confirmed at the protein level in both control and IPF fibroblasts (Figure 5b).

**Figure 5:**
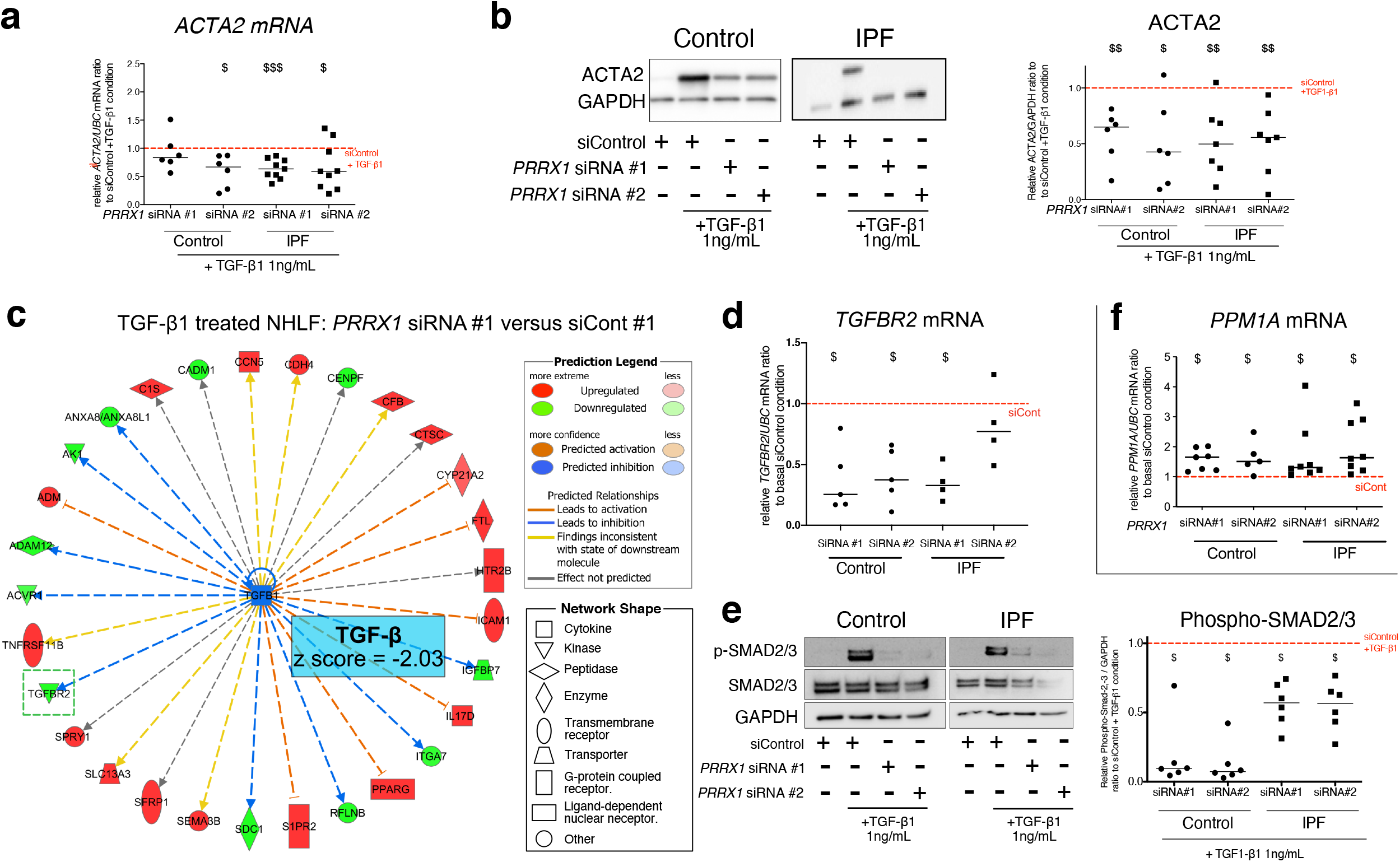
*PRRX1* inhibition decreased myofibroblast differentiation upon TGF1-β1 stimulation. (**a**) Dot plots with median showing the mRNA expression of *ACTA2* relative to the siControl +TGF-β1 condition (red dashed line), in control (circle, n=6) and IPF (square, n=9) lung fibroblasts treated with TGF-β1 and PRRX1 siRNA (#1 or #2). (**b**) Immunoblot showing ACTA2 expression in control (n=6) and IPF (n=7) fibroblasts treated with siControl in absence or presence of TGF-β1 or with *PRRX1* siRNA and TGF-β1. The quantification of ACTA2 expression relative to GAPDH (loading control) in control and IPF fibroblasts treated with control or *PRRX1* siRNA in presence of TGF-β1 relative to siControl + TGF-β1 condition is displayed as dot plot with median on the right. (**c**) Ingenuity Pathway Analysis of whole transcriptome in NHLF treated for 48h with *PRRX1* siRNA in the presence of TGF-β1 indicated that the best predicted upstream regulator was TGFB1 (z score=-2.03, *PRRX1* siRNA#1 versus siControl#1, n=2). Inhibition of *TGFBR2* is framed with a green dashed border. Figure legend displays molecules and function symbol types and colors. (**d**) Dot plots with median showing the mRNA expression of *TGFBR2* (n=4 to 5) r*e*lative to siControl in control and IPF fibroblasts treated for 48h with *PRRX1* siRNA. **(e**) Immunoblot showing phospho-SMAD2/3 and SMAD2/3 expression in control and IPF fibroblasts treated for 30 minutes with TGF-β1 after 48h transfection with *PRRX1* siRNA. The quantification of phospho-SMAD2/3 and SMAD2/3 expression relative to GAPDH (loading control) in control (n=6) and IPF (n=6) lung fibroblasts treated for 30 minutes with TGF-β1 after 48h transfection with *PRRX1* siRNA relative to siControl + TGF-β1 condition (red dashed line), is displayed as dot plot with median on the right. (**f**) Dot plots with median showing the mRNA expression of *PPM1A* (n=7 to 8) r*e*lative to siControl in control and IPF fibroblasts treated for 48h with *PRRX1* siRNA. *Abbre*via*tions: control siRNA sequence (siControl)*, β-Tubulin (TUB). Wilcoxon signed-rank test, $ p≤0.05, $$ p≤0.01, $$$ p≤0.001.

With respect to ECM synthesis, *PRRX1* knock down did not influence *FN1* or *COL1A1* upregulation after TGF-β1 stimulation, both at mRNA and protein levels (Supplemental Figure S6). Nevertheless, other ECM proteins associated with IPF were modulated after PRRX1 down regulation in presence of TGF-β1. For instance, the expression of *TNC* mRNA was increased in *PRRX1* siRNA treated control and IPF lung fibroblasts compared to control siRNA treated in presence of TGF-β1 (supplemental Figure S6). Meanwhile, the expression of *ELN* mRNA was downregulated in IPF lung fibroblasts (supplemental Figure S6) after TGF-β1 stimulation.

### PRRX1 TFs modulate SMAD2 and SMAD3 phosphorylation in response to TGF-β1 by regulating the expression of TGFβ Receptor 2 (TGFBR2) and the serine/ threonine phosphatase PPM1A

To better appreciate the *PRRX1* siRNA effects upon TGF-β1 pathway, whole transcriptome profiling was performed on NHLF treated with *PRRX1* or control siRNAs for 48h and then in presence or absence of TGF-β1 (for an additional 48h). Ingenuity Pathway Analysis at 96h indicated that the most significantly modulated pathway by *PRRX1* inhibition was the TGF-β1 pathway, which was significantly inhibited in TGF-β1-stimulated NHLF treated with *PRRX1* siRNA compared to control siRNA (Figure 5c, supplemental Figure S7 and supplemental Table S2).

Interestingly, *PRRX1* knockdown significantly affected the expression of the transmembrane Serine/Threonine kinase receptor *TGFBR2*, a key component of the TGF-β pathway (Figure 5c and supplemental Figure S7). We confirmed this observation in control and IPF fibroblasts at mRNA and protein levels (Figure 5d and supplemental Figure S8). We performed a ChIP assay to investigate a possible interaction of PRRX1 TFs with *TGFBR2* gene promoter regions. However, we detected no enrichment in PRRX1 TF binding in *TGFBR2* promoter regions by ChIP in primary NHLF (data not shown).

TGFBR2 is part of the receptor complex with TGFBR1 controlling TGF-β/SMAD signaling cascade by promoting SMAD2 and SMAD3 phosphorylation upon TGF-β1 stimulation. We assayed SMAD2 and SMAD3 phosphorylation in control and IPF fibroblasts treated with *PRRX1* siRNA, compared to cells transfected with control siRNA, in presence or absence of TGF-β1 (30min stimulation) (Figure 5e). In control and IPF fibroblasts, PRRX1 knock down strongly inhibited TGF-β1-induced SMAD2 and SMAD3 phosphorylation (Figure 5e). Most importantly, *PRRX1* knock down did not inhibit the activation of non-canonical/SMAD-independent TGF-β receptor-mediated signalling pathway such as AKT and JNK, in both control and IPF fibroblast (data not shown).

To understand this discrepancy between inhibition of SMAD2/3 phosphorylation and persistent activation of non-canonical pathways, we investigated whether PRRX1 TFs may regulate the expression of intracellular phosphatases known to control SMAD2 and SMAD3 phosphorylation downstream of TGF-β receptor activation ^25^. We observed that *PRRX1* siRNA-mediated inhibition was associated with an increase of PPM1A, a phosphatase member of the PP2C protein family, at both mRNA and protein levels compared to control and IPF fibroblasts treated with control siRNA (Figure 5f and supplemental Figure S8). In addition, siRNA-mediated inhibition of *PPM1A* partially rescued SMAD3 phosphorylation levels after TGF-β1 stimulation in *PRRX1* siRNA treated control and IPF fibroblasts (Figure 5F and supplemental Figure S8). Next, we performed a ChIP assay in primary NHLF to investigate a possible interaction of PRRX1 TFs with *PPM1A* gene *loci*. Indeed, we detected an enrichment in PRRX1 TF binding in *PPM1A* promoter regions (supplemental Figure S8).

Altogether, our results suggested that PRRX1 TFs are required to achieve proper myofibroblastic differentiation upon TGF-β1 stimulation. The effect is at least partially mediated through the regulation of TGFβ receptor 2 and PPM1A phosphatase expression.

### PRRX1 TFs expression levels are upregulated in the bleomycin-induced model of lung fibrosis

In the light of our in vitro results regarding PRRX1 fundamental role in the control of fibroblast proliferation and TGF-β1 responsiveness, we investigated whether alteration in PRRX1 expression may also contribute to fibrogenesis in the bleomycin-induced model of lung fibrosis (single intratracheal instillation ^26^). In this model, the expression levels of both *Prrx1* isoforms mRNA were mainly increased during the fibrotic phase from day 7 compared to the control PBS treated animals (Figure 6a). The upregulation of PRRX1 expression level was confirmed at the protein level only at day 14 during fibrosis phase peak (Figure 6b).

**Figure 6:**
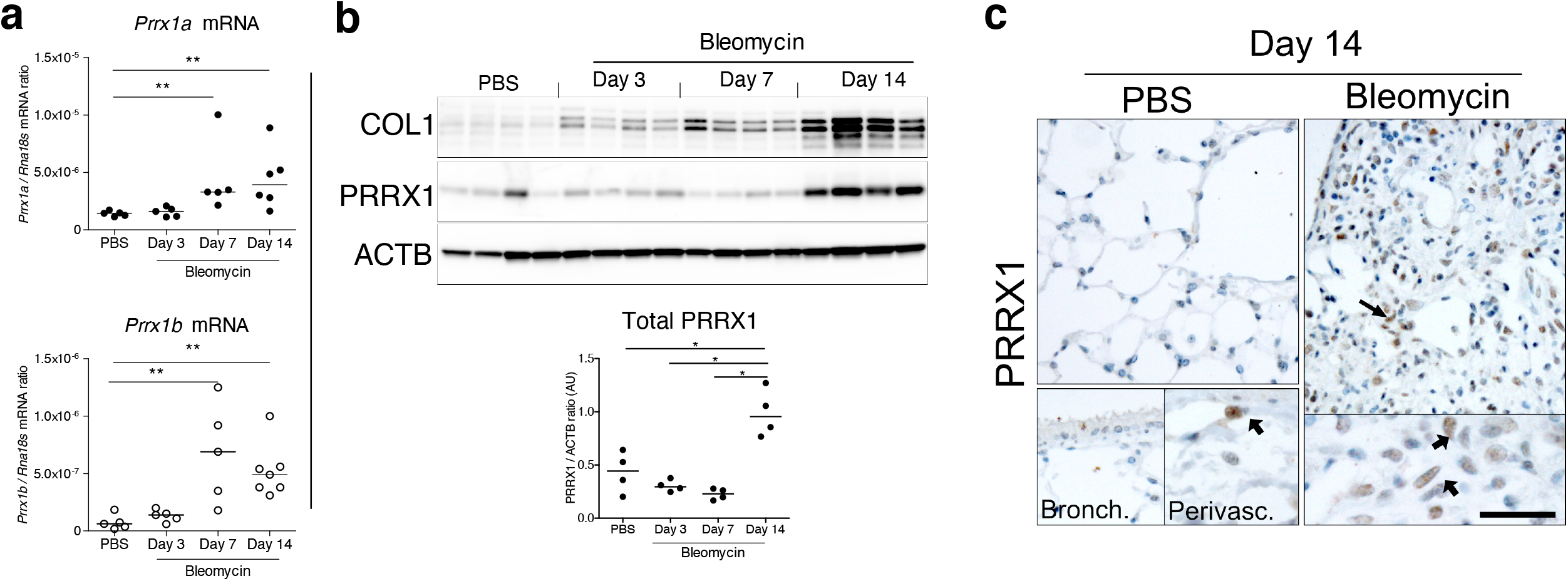
PRRX1 is increased during fibrotic phase in mice bleomycin-induced fibrosis. (**a**) Dot plots with median showing the mRNA expression of *Prrx1a* (black circle) and *Prrx1b* (white circle) isoforms in PBS mice (n=5) and bleomycin-treated mice at day 3 (n=5), 7 (n=5) and 14 (n=6). (**b**) Immunoblot showing COL1 and PRRX1 expression in PBS and bleomycin-treated mice at day 3, 7 and 14 (n=4 per group). ACTB was used as loading control. The quantification of PRRX1 expression relative to ACTB in PBS and bleomycin mice is displayed as dot plot with median in the lower part of the panel. (**c**) Representative immunohistochemistry pictures (n= 3 per group) showing PRRX1 staining (brown) in PBS and bleomycin mice at day 14 after saline or bleomycin administration. Nuclei were counterstained with hematoxylin. Note the absence of PRRX1 staining in the bronchiolar epithelium. PRRX1 positive cells were only detected in the peri-vascular spaces (arrows) in naive mice lungs (lower left panels). In bleomycin-treated mice, PRRX1 positive cells (arrow) were detected in the remodeled/fibrotic area (right panels). Scale bar: 30 µm in low magnification images and 15µm in high magnification ones. *Abbre*via*tions: bronchiolar (bronch); perivascular (perivasc)*. Kruskal-Wallis test with Dunns post-test, *p≤0.05, **p≤0.01.

Similarly to control Human lungs, PRRX1 positive cells were detected only within the peri-vascular and peri-bronchiolar spaces in PBS control mice, while the distal alveolar space and the bronchiolar epithelium were devoid of PRRX1 staining as assayed by immunohistochemistry. Meanwhile, PRRX1 positive cells were detected in the remodeled fibrotic area of bleomycin treated animals at day 14 (Figure 6c). In summary, our results indicated that PRRX1 TFs upregulation was associated with fibrosis development in the bleomycin–induced model of lung fibrosis.

### In vivo inhibition of PRRX1 dampens experimental lung fibrosis

Since *Prrx1* loss of function is associated with perinatal lethality in *Prrx1*^*-/-*^ pups ^4,5,27^, we sought first to evaluate *Prrx1* function during lung fibrosis using *Prrx1*^*+/-*^ heterozygous mice. We observed that the loss of one *Prrx1* allele was not associated with any haploinsufficiency (supplemental Figure S9) and those *Prrx1*^*+/-*^ heterozygous mice were not protected from lung fibrosis at day 14 after intratracheal instillation of bleomycin (supplemental Figure S9).

In order to evaluate the involvement of PRRX1 TFs in pulmonary fibrosis in vivo, we then chose to treat wild type mice with a third generation antisense LNA-modified oligonucleotide (ASO) targeting both *Prrx1* isoforms in the bleomycin-induced model of lung fibrosis. The control or *Prrx1* ASO were administrated during the fibrotic phase (from day 7 to 13) by an endotracheal route to target specifically the lung. As compared to control ASO, *Prrx1* ASO strongly reduced the expression of both *Prrx1* isoforms at the mRNA and protein levels (Figure 7a-b) and reduced the extent of lung lesions on day 14 (Figure 7c). Lung collagen content was decreased as assessed with picrosirius staining, immunohistochemistry and hydroxyproline assay (Figure 7d-f). A similar decrease in ACTA2 staining was observed in *Prrx1* ASO treated animals at day 14 by immunohistochemistry (Figure 7e). In addition, *Prrx1* ASO decreased *Col1a1, Fn1* and *Acta2* mRNA content (Figure 8a) in *PRRX1* ASO treated bleomycin mice compared to control ASO treated ones. Finally, the expression levels of COL1, FN1 and ACTA2 were also decreased at the protein level as assayed by Western Blot (Figure 8b-c). The dampened fibrosis development observed in *PRRX1* ASO treated bleomycin mice was also associated with a decrease in key fibrosis mediators such as *Tgfb1* and *Ctgf* as well as inflammatory markers as *Tnf* and *Serpin-1* at the mRNA level (supplemental Figure S10). Furthermore, *Prrx1* was recently identified as the master transcription factor in the *Col14a1* subtype mesenchymal cell during fibrogenesis in this experimental model ^28^. Interestingly, the expression of *Col14a1* mRNA was also strongly decreased in the *Prrx1* ASO treated animals compared to control ASO at day 14 as assayed by qPCR (supplemental Figure S10). With respect to another fibrosis-associated key ECM protein, *Tnc* mRNA level was also decreased in the *Prrx1* ASO treated bleomycin group (supplemental Figure S10).

**Figure 7:**
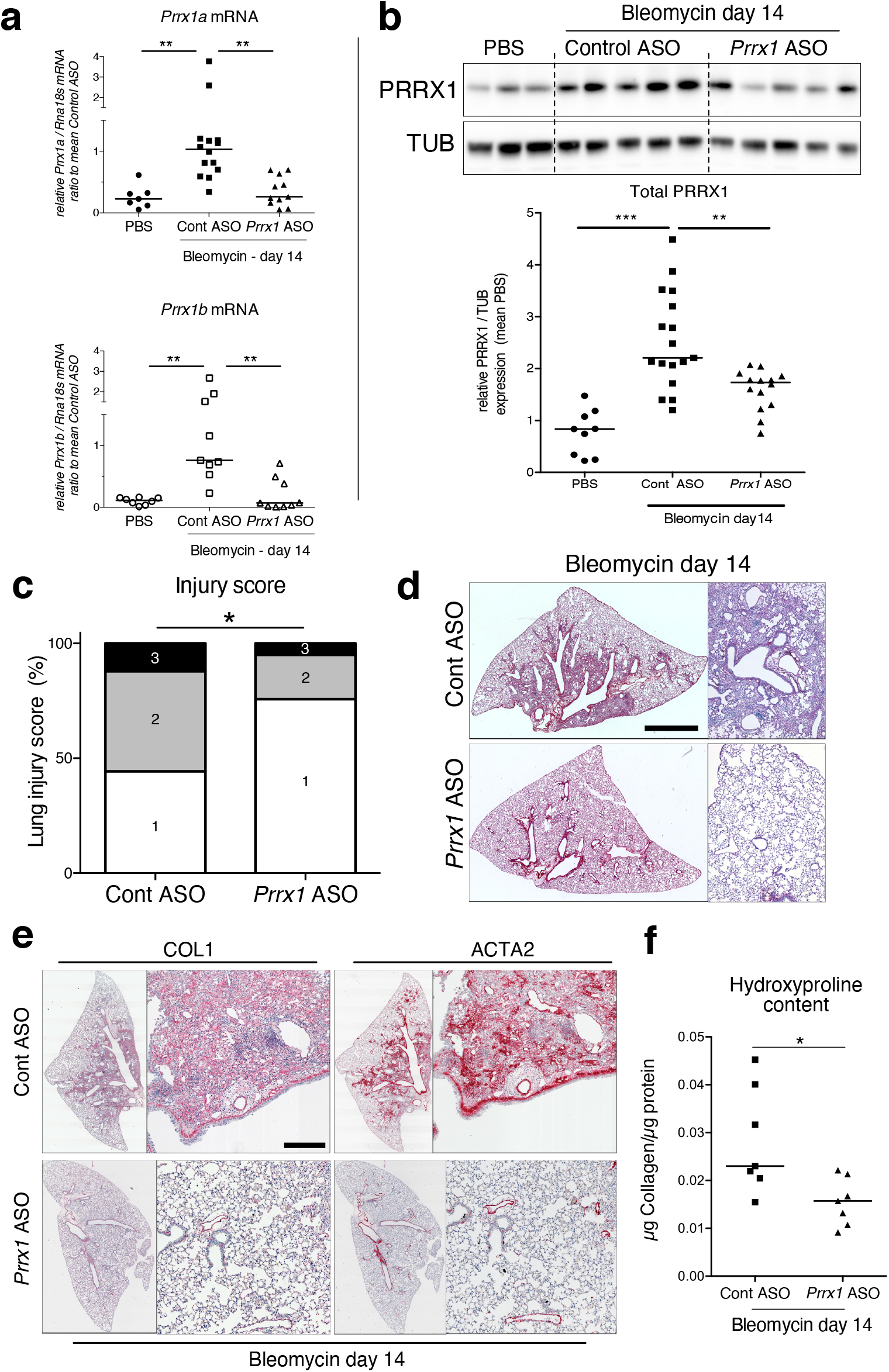
PRRX1 inhibition attenuates lung fibrosis in bleomycin murine model. (**a**) Dot plots with median showing the mRNA expression of *Prrx1a* (black) and *Prrx1b* (white) isoforms at day 14 (n=8) in PBS (circle) mice and bleomycin mice treated with control ASO (square, n=14) or *PRRX1* ASO (triangle, n=11). (**b**) Immunoblot showing PRRX1 expression at day 14 in PBS mice and bleomycin mice treated with Control ASO or *Prrx1* ASO. TUB was used as loading control. The quantification of PRRX1 expression relative to TUB at day 14 in PBS mice (circle, n=9) and bleomycin mice treated with Control ASO (square, n=16) or *Prrx1* ASO (triangle, n=14) is displayed as dot plot with median on the lower panel. (**c**) Injury score at day 14 of bleomycin mice treated with *Prrx1* ASO or Control ASO. (**d**) Representative immunohistochemistry images (n= 7 per group) showing picrosirius staining (red) at day 14 in bleomycin mice treated with Control ASO or *Prrx1* ASO. (**e**) Representative immunohistochemistry images (n= 7 per group) showing COL1 (left panel) and ACTA2 (right panel) staining (red) at day 14 in bleomycin mice treated with Control ASO or *Prrx1* ASO. Nuclei were counterstained with hematoxylin. (**f**) Dot plot with median showing the relative Collagen content as measured by hydroxyproline at day 14 in bleomycin mice treated with control ASO (square, n=7) or *PRRX1* ASO (triangle, n=7). Scale bar: 80µm in low magnification images and 40µm in high magnification ones. *Abbreviations: Control (Cont), Antisense oligonucleotide (ASO)*. Kruskal-Wallis test with Dunns post-test (A and B), Fisher’s exact test (C) and Mann Whitney U test (F); *p≤0.05, **p≤0.01, ***p≤0.001

**Figure 8:**
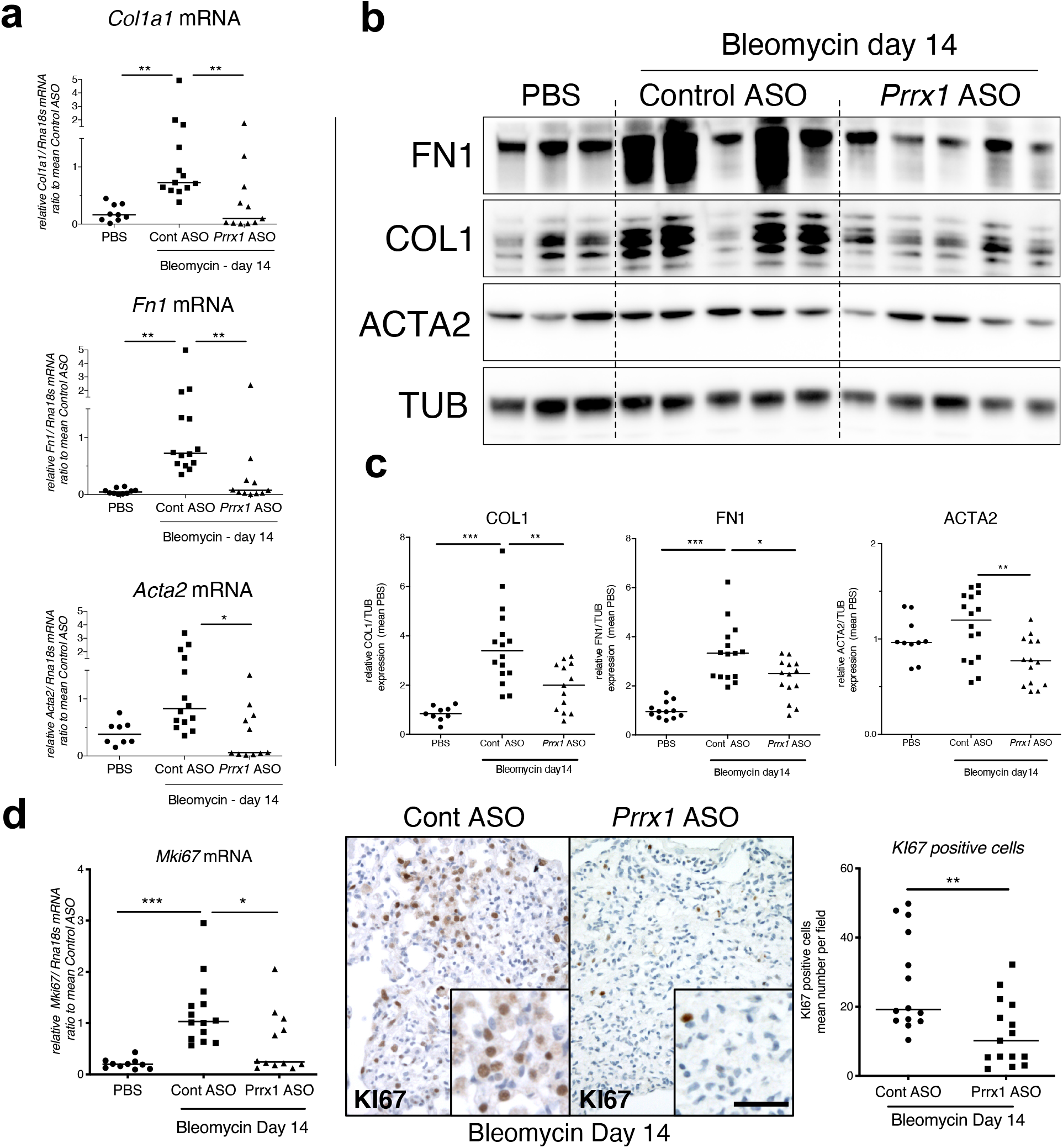
PRRX1 inhibition decreases fibrosis markers in bleomycin mice. (**a**) Dot plots with median showing the mRNA expression of *Col1a1, Fn1* and *Acta2* at day 14 in PBS mice (circle, n=9) and bleomycin mice treated with Control ASO (square, n=14) or *Prrx1* ASO (triangle, n=11). (**b**) Immunoblot showing FN1, COL1 and ACTA2 expression at day 14 in PBS mice and bleomycin mice treated with Control ASO or *Prrx1* ASO. TUB was used as loading control. (**c**) Quantification of FN1, COL1 and ACTA2 relative expression to TUB at day 14 in PBS mice (n=9) and bleomycin mice treated with Control ASO (n=16) or *Prrx1* ASO (n=13) (**d**) Left panel: dot plots with median showing *Mki67* mRNA expression of at day 14 in PBS mice (circle, n=10) and bleomycin mice treated with Control ASO (square, n=14) or *Prrx1* ASO (triangle, n=14). Middle panel: representative immunohistochemistry pictures (n= 14 per group) showing KI67 staining (brown) in bleomycin treated with Control ASO (left) or *Prrx1* ASO (right) mice at day 14. The quantification of the number of KI67 positive cells per high magnification field is shown on the right as dot plots with median. Scale bar: 40µm in low magnification images and 20µm in high magnification ones. *Abbreviations: Control (Cont), Antisense oligonucleotide (ASO)*. Kruskal-Wallis test with Dunns post-test (A, B) and Mann Whitney U test (C); *p≤0.05, **p≤0.01f, ***p≤0.001

The expression levels of the proliferation marker *Mki67* was decreased at the mRNA levels in the *Prrx1* ASO treated animals compared to control ASO at day 14 as assayed by qPCR (Figure 8d). A decrease in KI67 positive cells was also observed by immunochemistry in the *Prrx1* ASO treated animal lungs after bleomycin challenge compared to control ASO ones at day 14 (Figure 8d).

These data demonstrate that PRRX1 targeting in the lung has the potential to inhibit lung fibrosis development.

### PRRX1 inhibition attenuates lung fibrosis in mouse and Human precision-cut lung slices (PCLS)

To confirm the effect of the *Prrx1* ASO in a second model of lung fibrosis, we took advantage of a well-established ex vivo model of lung fibrosis using precision-cut lung slices (PCLS) derived from mouse and Human lung samples ^29^. PCLS have the major advantage to include the lung primary cell populations in a 3-dimensional preserved lung architecture and microenvironment. Our *Prrx1* ASO was designed to target both Human *PRRX1* and mouse *Prrx1* TFs orthologs. In basal condition, *Prrx1* knock down was associated with a decrease in *Acta2* mRNA expression in mouse PCLS (supplemental Figure S11). Next, mouse or Human control lung PCLS were treated with a fibrosis cytokine cocktail (FC) consisting of TGF-β1, PDGF-AB, TNFα and LPA to trigger fibrosis-like changes ^29^. In both mouse and Human PCLS, *ACTA2, COL1A1* and *FN1* mRNA upregulation was lessened in FC-stimulated PCLS with *PRRX1* ASO compared to control (Figure 9a-c). We confirmed those findings at the protein levels for ACTA2 and COL1 by western blot in mouse PCLS (Figure 9b). In Human PCLS stimulated with FC, morphological analysis revealed that *PRRX1* ASO treatment was associated with a decreased Collagen accumulation compared to control ASO (Figure 9d).

**Figure 9.**
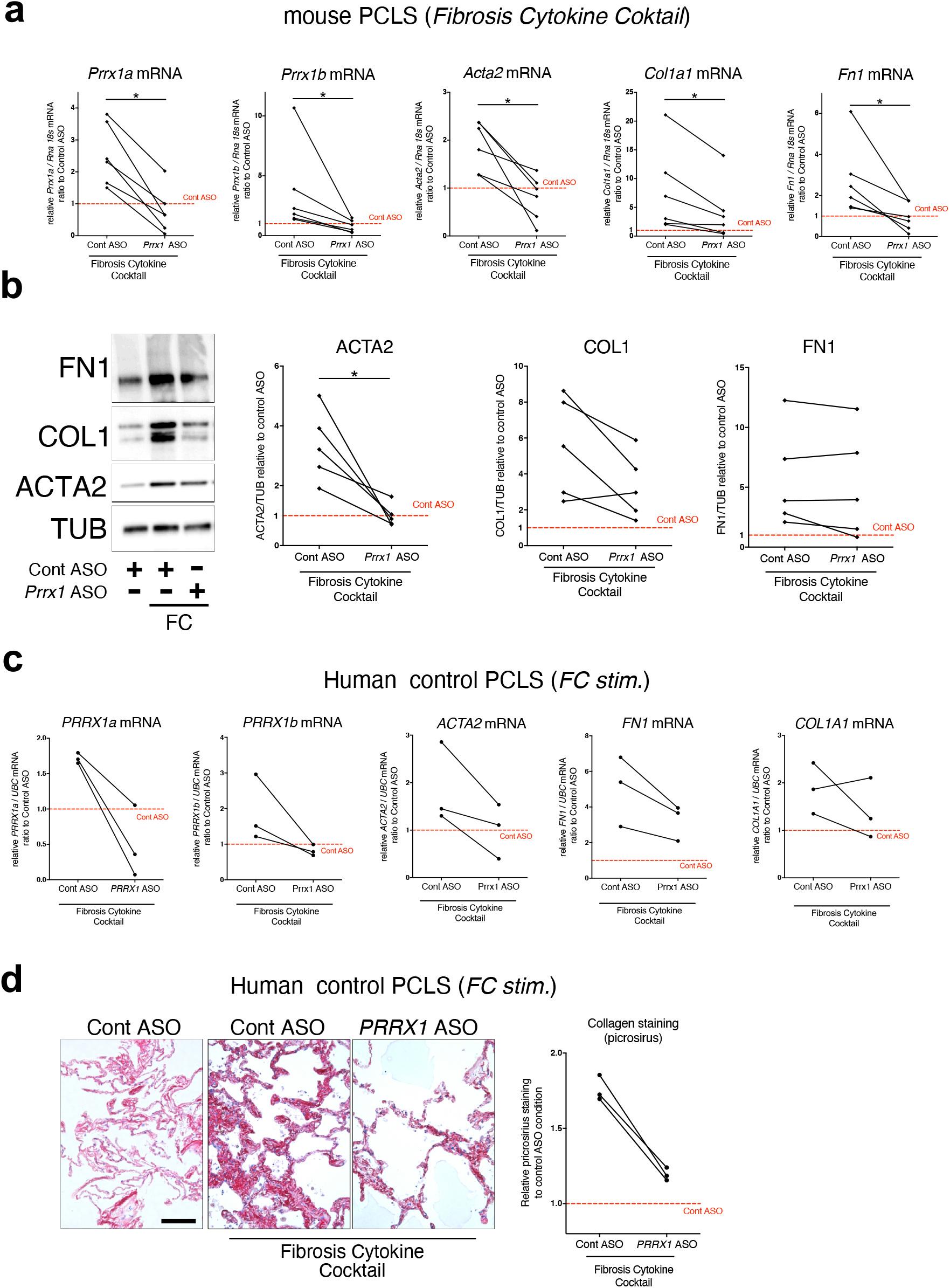
*PRRX1* ASO attenuates lung fibrosis in mouse and Human Precision-cut Lung slices (PCLS) (**a**) Before-after plots showing the mRNA expression of *Prrx1a, Prrx1b, Acta2, Col1a1* and *Fn1* (n=6) relative Control ASO alone condition (red dashed line) in mouse PCLS stimulated with fibrosis cytokine cocktail (FC) and then treated either with control ASO or *PRRX1* ASO. (**b**) Representative immunoblot showing FN1, COL1 and ACTA2 expression relative to control ASO alone condition (red dashed line) in mouse PCLS stimulated with FC and then treated either with control or *Prrx1* ASO. The corresponding quantifications of ACTA2, COL1 and FN1 expression ratio to Tubulin are displayed as before-after plots on the right. Note that COL1 expression was decreased in 4 out 5 experiments. (**c**) Before-after plots showing the mRNA expression of *PRRX1a, PRRX1b, ACTA2, COL1A1* and *FN1* (n=3) relative Control ASO condition (red dashed line) in Human PCLS treated either with control ASO or *PRRX1* ASO in presence or absence of FC. *COL1A1* upregulation was lessened in 2 out 3 experiments while ACTA2 and FN1 levels were decreased in 3 out of 3 experiments. (**d**) Representative picrosirius staining (n=3) in Human PCLS treated with control ASO alone (left panel, basal condition) or after stimulation with Fibrosis Cytokine cocktail and treated with either control (middle panel) or *PRRX1* (right panel) ASO. Nuclei were counterstained with hematoxylin. The quantification of picrosirius staining relative to control ASO alone (red dashed line) is showed on the right (Before-after plot). Scale bar: 50µm. *Abbre*via*tions: Precision-Cut Lung slices (PCLS), Fibrosis Cytokine Cocktail (FC), Control (Cont), Antisense oligonucleotide (ASO), Stimulation (stim*.). Wilcoxon test * p≤0.05.

Altogether, these results demonstrate that inhibition of PRRX1 transcription factors, using an ASO approach, reduced fibrosis development in vivo and ex vivo.

## DISCUSSION

This is the first study to evidence the critical role of the PRRX1 transcription factors in lung fibrosis pathophysiology. Our results demonstrate that 1) PRRX1 TFs are upregulated in mesenchymal cells accumulating in the fibrotic areas of IPF lungs, 2) the expression of PRRX1 TFs is positively regulated by cues associated with an undifferentiated phenotype in control and IPF primary lung fibroblasts, 3) PRRX1 TFs are required for proliferation as well as proper myofibroblastic differentiation in vitro (see Figure 10 for summary). We identified the underlying mechanisms; including PRRX1 TFs effects on cell cycle (modulation of cyclins and MKI67) and on SMAD 2/3 phosphorylation (regulation of TGFBR2 and phosphatase PPM1A) respectively (see Figure 10). Finally, inhibition of *Prrx1* with LNA-modified ASO strongly impacted lung fibrosis development in in vivo and ex vivo preclinical models.

**Figure 10:**
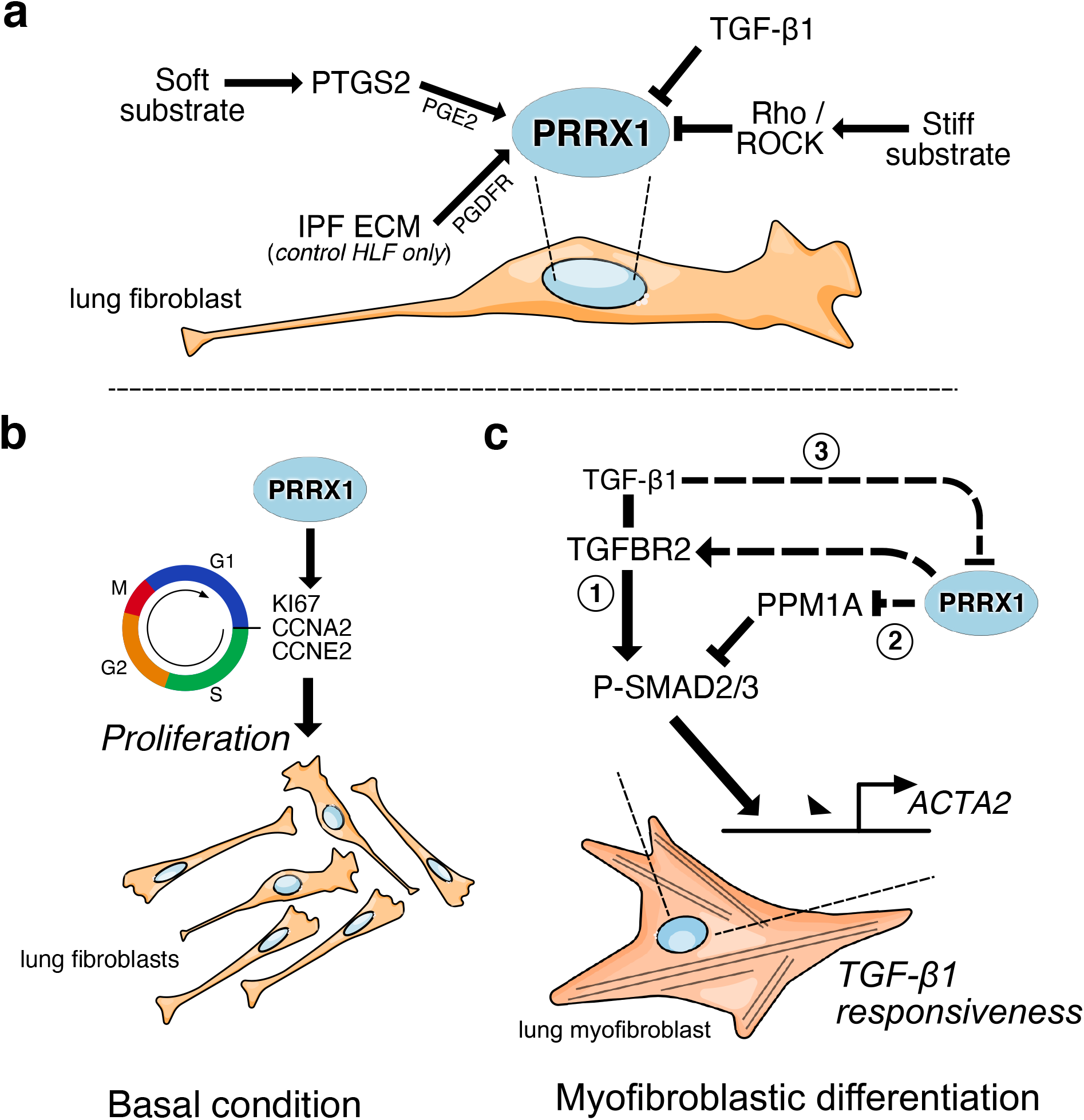
summary sketch of PRRX1 regulation and functions in lung fibroblasts. (**a**) Regulation of *PRRX1* TF expression in lung fibroblasts. On one hand, PRRX1 expression was up-regulated by the anti-fibrotic factor PGE2 and soft culture substrate (in a PTGS2 dependent-manner). IPF fibroblast-derived matrix also increased *PRRX1* TFs expression in a PDGFR dependent manner in control primary lung fibroblasts only. On the other hand, stiff culture substrate (in a Rho/ROCK dependent manner) and TGF-β1 stimulation, which both promote myofibroblastic differentiation, decreased PRRX1 TF expression levels in both control and IPF fibroblasts seeded on plastic. (**b**) Model of PRRX1 function in lung fibroblasts at steady state. In complete growth medium, PRRX1 TFs influence cell cycle progression by regulating key factors associated with cycle progression during the G1 and S phases (KI67, Cyclin A2 and E2). PRRX1 was detected in the promoter regions of those genes by chromatin immunoprecipitation (ChIP). (**c**) Model of PRRX1 function in lung fibroblasts during myofibroblastic differentiation. (1) TGF-β1 stimulation of lung fibroblasts will trigger their differentiation into myofibroblasts by promoting the phosphorylation of SMAD2 and SMAD3. P-SMAD2/3 will then induce the upregulation of ACTA2 expression. (2) In presence of PRRX1, the expression of the serine / threonine phosphatase PPM1A is downregulating (PRRX1 TFs binding to PPM1A promoter region was demonstrated by ChIP) and TGFBR2 expression is also maintained. Thus, the phosphorylation of SMAD2 and SMAD3 is therefore not impacted. (3) During myofibroblastic differentiation, the expression of PRRX1 TFs was then decreased after TGF-β1 treatment for 48h (not at 24h). This negative feedback loop could limit cell-responsiveness to long exposure of TGF-β1 by upregulating the expression of PPM1A and downregulating TGFBR2 levels. *Abbre*via*tions: IPF (Idiopathic Pulmonary Fibrosis), HLF (Human Lung Fibroblasts), ECM (Extracellular matrix), G1 (Gap 1 phase 1), S (Synthesis / Replicative phase), G2 (Gap phase 2), M (Mitosis), CCNA2 (Cyclin A2), CCNE2 (Cyclin E2)*.

### Reactivation of the developmental and mesenchyme-associated PRRX1 transcription factors in IPF

The *Prrx1* gene encodes transcription factor isoforms (*Prrx1a* and *1b*) involved in the maintenance of cell fate within the limb and craniofacial mesenchyme during ontogeny ^4^. *Prrx1* is also required for cell fate decision during lung development. *Prrx1*^*-/-*^ newborn display hypoplastic lungs and die at birth from respiratory distress ^5^. PRRX1 TFs were also identified as key drivers of mesenchymal phenotype acquisition during epithelial-mesenchyme transition (EMT) in cancer ^8^. A *Prrx1* positive fibroblast subpopulation was also recently characterized as pro-fibrotic in the mouse ventral dermis ^30,31^. Interestingly, *Prrx1* mRNA expression was associated with a sub-population of matrix fibroblast in the bleomycin experimental mouse model of lung fibrosis using single-cell RNA sequencing ^28^.

However, the role of PRRX1 TFs in IPF, a disease associated with major perturbations in mesenchymal compartment was not known. We first identified *PRRX1* as potentially upregulated TF in IPF lung after analysis of public transcriptomic database comparing control and IPF lungs ^32–34^. *PRRX1* also appeared in the upregulated gene hit list of a subgroup of IPF patients, displaying an accelerated clinical course in another transcriptomic study ^13^. We confirmed that the mRNA levels of *PRRX1* TFs isoforms were actually increased in IPF lung patients as well as in primary fibroblast isolated from IPF lung compared to control ones. This increase was confirmed at the protein level in IPF whole lung extracts compared to control ones. The upregulation of PRRX1 protein levels in fibrotic lungs may therefore reflect the accumulation of PRRX1 positive cells in IPF. Indeed, PRRX1 expression was restricted in control lungs to interstitial fibroblasts within peri-vascular and peri-bronchiolar space. In IPF lungs, PRRX1 was strongly detected in the nucleus of fibroblasts organized in foci as also observed by others ^24^ and in scattered mesenchymal cells within the remodeled / fibrotic lung areas. Our findings are also supported by recent single cell transcriptomic studies performed in donor and fibrotic lungs ^14,15^. Datamining from these studies confirmed that *PRRX1* TF expression was restricted to lung mesenchymal lineage.

### *PRRX1* TFs expression is increased in lung fibroblasts by cues promoting an undifferentiated state

Our data suggest that PRRX1 expression is controlled by a PGE2/TGF-β balance in lung fibroblasts in vitro.

On one hand, PGE2 up-regulated *PRRX1a* and *1b* expression in both control and IPF fibroblasts. Substrate stiffness in physiological range also increased *PRRX1* isoforms expression in a PTGS2 dependent manner. The PTGS2/PGE2 axis is known to promote an undifferentiated state in fibroblasts ^17^. On the other hand, signals triggering myofibroblastic differentiation (TGF-β1 stimulation and stiff substrate) decreased *PRRX1* TFs levels in primary lung fibroblasts (Figure 10). Interestingly, several studies reported that PRRX1 expression level was rather increased upon the activation of the TGF-β pathway in other cell types such as mouse embryonic lung mesenchymal cells ^35^, embryonic mouse 3T3-L1 adipocyte precursor ^7^ and transformed epithelial cells undergoing EMT ^8^. PRRX1 upregulation in response to TGF-β1 in the two later cell types promoted their dedifferentiation toward a more plastic phenotype ^7,8^. Conversely, *PRRX1* downregulation in primary lung fibroblasts grown in presence of TGF-β1 was associated with a differentiation process toward myofibroblastic phenotype.

Overall, these different studies and our results strongly suggested that PRRX1 expression is associated with an undifferentiated phenotype. In addition, *PRRX1a* and -*1b* TF mRNA expression levels were upregulated only in control fibroblasts seeded on IPF fibroblast-derived 3D ECM in a PDGFR dependent manner. *PRRX1* TF mRNA levels seemed to be regulated by both ECM origin and stiffness in control fibroblasts, while it was only modulated by the latter in IPF fibroblasts.

### PRRX1 TFs drive key basic fibroblast functions involved in fibrogenesis

In adult lung fibroblasts, PRRX1 TFs appeared to strongly influence cell cycle progression and myofibroblastic differentiation, two entangled cellular processes (Figure 10). There was generally no difference between control and IPF fibroblasts regarding *PRRX1* functions in those cells (with respect to proliferation and myofibroblastic differentiation at least). Overall, this may suggest that *PRRX1* TFs function might be central to fibroblast biology independently of their origin (control versus IPF lungs). However, differential *PRRX1* regulation between control and IPF fibroblasts by the micro-environment or soluble factors could therefore have a higher impact on PRRX1 overall function in lung fibroblasts.

While PRRX1 TFs are required for fibroblast proliferation in complete growth medium, PRRX1 partial loss of function perturbed only some key features of myofibroblastic differentiation in response to TGF-β1 stimulation (see Figure 10). Only the expression of markers involved in the acquisition of contractile properties (ACTA2 and *ACTG2)* was decreased in a SMAD2/3 dependent way. The effect of PRRX1 inhibition upon P-SMAD3 was partially mediated through TGFBR2 downregulation and the upregulation of the PPM1A phosphatase ^25^, which are critical components of the canonical TGF-β/SMAD signalling cascade. Whole transcriptome profiling data performed in NHLF were also consistent with a global impact of *PRRX1* downregulation on TGF-β response in lung fibroblasts.

Meanwhile, the expression levels of key ECM proteins such as Collagen 1 and Fibronectin were still upregulated in TGF-β1 stimulated lung fibroblasts, transfected with *PRRX1* siRNA. Even thought, SMAD3 phosphorylation was impacted, the non-canonical ERK, AKT and JNK pathways were still fully activated in those stimulated cells. The activation of those pathways has been previously showed to be sufficient to upregulate the expression of FN1 and Collagen 1 in fibroblasts ^2,25^. However, the expression of other IPF-associated ECM proteins such as TNC and ELN was perturbed in *PRRX1* siRNA treated control and IPF lung fibroblasts stimulated with TGF-β1. Our results suggest that PRRX1 inhibition in presence of TGF-β1 might promote a different myofibroblastic phenotype with potentially less contractile capability and with a different ECM secretome.

Albeit we showed that PRRX1 is required for proper myofibroblastic differentiation, the expression of both PRRX1 isoforms was decreased after TGF-β1 treatment for 48h. This paradoxical downregulation of PRRX1 in response to TGF-β1, could be the signature of a negative feedback loop to limit cell-responsiveness to TGF-β1 long exposure (Figure 10). TGF-β1 induced PRRX1 inhibition in lung fibroblasts could also correlate with progressive proliferation loss during the differentiation process.

Overall, we propose that PRRX1 TFs would maintain lung mesenchymal cells in an undifferentiated and proliferative state but would also act as enablers to promote full myofibroblastic differentiation in response to pro-fibrotic cues such as TGF-β1.

### Inhibition of the mesenchymal PRRX1 transcription factor is sufficient to dampen lung fibrosis in vivo

PRRX1 TFs expression levels were also upregulated during the fibrosis phase at day 14 in mice treated with bleomycin (intratracheal route). However, the *Prrx1* homozygous and heterozygous mice were not suitable for studying *Prrx1* function during lung fibrosis in adult mice. Indeed, *Prrx1*^*-/-*^ mice present a lethal respiratory failure at birth ^4,5,27^ and the lack of haploinsufficiency in *Prrx1*^*+/-*^ heterozygous mice did not prevent lung fibrosis development after intratracheal instillation of bleomycin. Thus, we chose to inhibit *Prrx1* in adult mice using a LNA-modified ASO ^36^ targeting both *Prrx1* isoforms. This ASO was administered by the endotracheal route in a “curative” protocol from day 7 in this experimental mouse model of pulmonary fibrosis. LNA-modified ASOs are protected from nuclease-mediated degradation, which significantly improves their stability and prolongs their activity in vivo. In comparison to earlier generations of ASO modifications, they have a massively increased affinity to their target RNA and their in vitro and in vivo activity does not depend on delivery reagents ^36^. As a proof of concept, we confirmed that intratracheal administration of *Prrx1*-specific ASO inhibited the upregulation of mouse PRRX1a and -1b expression at both mRNA and protein level at day 14 in bleomycin treated mice. Pulmonary fibrosis development was also reduced in these animals. While our in vitro findings in adult Human lung fibroblasts showed that PRRX1 inhibition mainly impacted ACTA2 expression levels, *Prrx1* ASO treatment in the bleomycin mouse model of lung fibrosis also inhibited the deposition of Collagen and Fibronectin. This difference regarding ECM compound at day 14 may reflect the effect of *Prrx1* ASO on the overall fibrosis development; upon the proliferation / accumulation as well as impaired myofibroblastic differentiation of mesenchymal cells in vivo from the beginning of the ASO treatment at day 7. We confirmed the anti-fibrotic effect of the *Prrx1* ASO in a second model of fibrosis using ex vivo culture of Human or mouse PCLS stimulated with a cocktail of fibrosis-associated cytokines ^29^.

Targeting of others transcription factors such as GLI ^26^, FOXM1 ^37^, FOXF1 ^38^, FOXO3 ^39^ and TBX4 ^40^ was also shown to inhibit fibrosis development in this mouse experimental model of pulmonary fibrosis. However, at the exception of TBX4 and PRRX1, the expression of all these other TFs is not restricted to mesenchymal lineages, which means that targeting those TFs may impact both lung fibrosis and epithelial regeneration/repair. Finally, PRRX1 inhibition as a potential therapeutic approach in fibrosis is not restricted to the lung. Recently, adenoviral shRNA mediated inhibition of *Prrx1* in the thioacetamide model of liver fibrosis in rats also decreased fibrotic lesions, collagen deposition and hepatic stellate cells myofibroblastic differentiation ^11^.

In conclusion, our study unveils the role of the pro-fibrotic and mesenchyme associated PRRX1 TFs in lung fibrosis by controlling fibroblasts proliferation and TGF-β pathway responsiveness during myofibroblastic differentiation. Direct inhibition of PRRX1 transcriptional activity in mesenchymal cells may be a potential therapeutic target in IPF. Furthermore, the effectiveness of the late administration of *Prrx1* ASO in the bleomycin model of pulmonary fibrosis is particularly interesting. The route of administration we used constitutes a first attempt to locally inhibit a pro-fibrotic TF. The possibility of a local administration of an antifibrotic is seductive: current antifibrotics, administered systemically, are burdened with significant adverse events, which significantly attenuates their effect on health-related quality of life ^41^. Inhaled pirfenidone and other inhaled compounds ^42^ are currently investigated in IPF, but none are directly acting on a mesenchymal transcription factor. Although already proved effective in asthma ^43^, local transcription factor inhibition has never been investigated in IPF so far.

## METHODS

### Human lung samples

IPF lung samples were obtained from patients undergoing open lung biopsy or at the time of lung transplantation (*n* =39; median age 61 yr; range 51–70 yr). IPF was diagnosed according to 2011 ATS/ERS/JRS/ALAT criteria, including histopathological features of usual interstitial pneumonia ^44^. Lung samples obtained after cancer surgery, away from the tumor, were used as controls; normalcy of control lungs was verified histologically (*n* =35 patients; median age 64 yr, range 28–83 yr).

### In vivo experiments

All experiments were performed using adult male C57BL/6 mice and intratracheal bleomycin administration, as previously described ^26^. To investigate the involvement of PRRX1 in fibrogenesis, mice were treated with third generation locked nucleic acid (LNA)-modified ASO targeting PRRX1 designed by Secarna Pharmaceuticals GmbH& Co, Planegg/Martinsried, Germany. The following sequence was used (+ indicates an LNA modification, while * indicates a phosphorothioate (PTO) linkage) to target *Prrx1* (*Prrx1* ASO):

+T*C*+A*+G*G*T*T*G*G*C*A*A*T*G*+C*+T*+G

A previously published and validated ^45^ negative control ASO (Cont ASO) was used:

+C*+G*+T*T*T*A*G*G*C*T*A*T*G*T*A*+C*+T*+T

Bleomycin control mice received only PBS. ASO and PBS were given by endotracheal instillation every other day from Day 7 after the bleomycin injection, until Day 14. All the mice were under isoflurane anesthesia during the instillation and received one injection every other day of 25µL with 20nmoles of ASO or PBS 1X. Lungs were harvested on Day 14 for further analysis. Hematoxylin, eosin and picrosirius staining were performed routinely to evaluate the morphology of the lung. Semiquantitative histological assessment of lung injury used the grading system described by Inoshima and colleagues ^46^. Total mRNA was extracted from mouse lung homogenates, and the expression of the genes of interest was quantified by real-time PCR, as previously described. Proteins were extracted from mouse lung homogenates and western blotting was performed by standard techniques as previously described ^26^. The *Prrx1* heterozygous mouse strain (129S-*Prrx1*^*tm1Jfm*^/Mmmh ^27^, RRID:MMRRC_000347-MU) was obtained from the Mutant Mouse Resource and Research Center (MMRRC) at University of Missouri (USA), an NIH-funded strain repository, and was donated to the MMRRC by Pr James Martin (Texas Agricultural and Mechanical University: Health Science Center, USA).

### Statistical Analysis

Most data are represented as dot plots with median, unless specified. All statistical analysis were performed using Prism 5 (GraphPad Software, La Jolla, CA). We used non-parametric Mann-Whitney U test for comparison between two experimental conditions. Paired data were compared with Wilcoxon signed-rank test. We used non-parametric Kruskall Wallis test followed by Dunn’s comparison test for group analysis. Comparison of histological scores on Day 14 was performed with Fisher’s exact test. A p-value *<* 0.05 was considered to be statistically significant. Exact *P* values and definition and number of replicates are given in the respective figure legend.

### Study approval

The study on human material was performed in accordance with the Declaration of Helsinki and approved by the local ethics committee (CPP Ile de France 1, No.0811760). Written informed consent was obtained from all subjects.

All animal experiments were conducted in accordance with the Directive 2010/63/EU of the European Parliament and approved by the local Animal ethics committee (“Comité d’éthique Paris Nord n°121”, APAFiS #4778 Etudedufacteurdetran_2016031617411315).

See supplementary materials for further details.

## Author contributions

EMD, MHL, AF, MJ, EF, MG, AJ, AM, AJ, AAM, LG, AV, MK, CMM carried out the experiments; KS and FJ designed and provided reagents; AC, HM, PM provided the lung samples; AAM, BC supervised the study. AAM, BC, MHL, AF, CMM, AG, BM and EMD designed the work, analyzed the data and wrote the manuscript. All authors reviewed and approved the manuscript.

## Acknowledgements

We thank Olivier Thibaudeau and Laure Wingertsmann (Morphology Platform, Inserm U1152 of X. Bichat Medical School, Paris) for their efficient collaboration and the Flow Cytometry Platform of Inserm U1149 (CRI, X. Bichat Medical School, Paris) as well. We also thank Dr. J.W. Duitman (Academic Medical Center, Amsterdam, The Netherlands) for his technical help. This work was supported by the ANR (JCJC ANR-16-CE14-0018), by grants from the Chancellery of Paris Universities (Poix Legacy), and by the “Association pour la fibrose pulmonaire idiopathique Pierre ENJALRAN”. E. Marchal-Duval was supported by “Ecole Doctorale Bio-SPC” (Grant 2015); A. Froidure an European Respiratory Society (ERS) and the European Molecular Biology Organization (EMBO) - ERS-LTRF 2015 – 4476 and by a fellowship from the Belgian Society for Pulmonology (BVP-SBP); M Homps-Legrand by the Fond de dotation “Recherche en Santé Respiratoire” (Grant 2018). A. Justet by the “Fondation Recherche Médicale” (Grant FRM 2016, FDM41320); M. Ghanem by the Fond de dotation “Recherche en Santé Respiratoire” (Grant 2015) and A Maurac by the ARS Lorraine (« année Recherche » 2016). We thank Alberto Baeri (IPMC), Nicolas Nottet (C3M, Nice) and Kevin Lebrigand (UCA genomics platform, IPMC) for their technical help. The authors declare no conflicts of interest.

