## Supplemental data for "Identification of Paired-related Homeobox Protein 1 as a key mesenchymal transcription factor in Idiopathic Pulmonary Fibrosis"

### SUPPLEMENTARY MATERIALS:

#### Supplemental figures

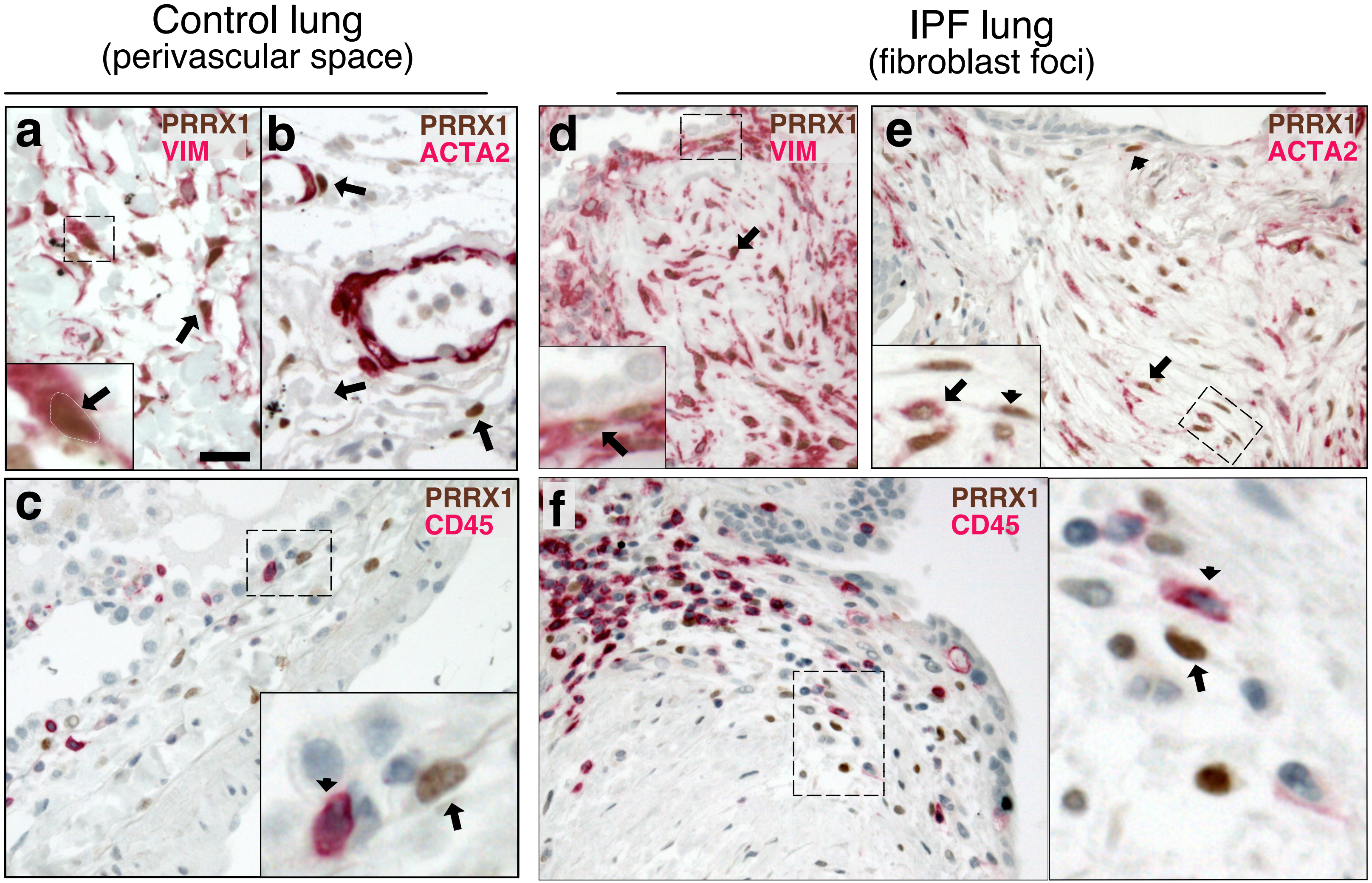


Supplemental Figure 1

**<u>Supplemental Figure S1:</u> Co-expression of PRRX1 with Vimentin and ACTA2 in control and IPF lungs by immunochemistry.**

(**a-c**) Representative immunohistochemistry images (n=3 per group) in control lung showing PRRX1 staining (Brown Chromogen, nuclear staining) in perivascular space with (a) Vimentin (Red chromogen), (b) ACTA2 (Red chromogen) and (c) CD45 (Red chromogen). Note that PRRX1 positive cells (arrow) were Vimentine positive (a) but ACTA2 (b) and CD45 negative (c). Insert in (a): high magnification of the black dashed box in the main left panel showing a double positive PRRX1 (brown nuclear staining, outlined with dashed white line) and Vimentin (red cytoplasmic staining) cell. Insert in (c): high magnification of the black dashed box in the main bottom panel showing a PRRX1 (brown nuclear staining) positive but CD45 negative cell (arrow) as well as a CD45 (red cytoplasmic staining) positive but PRRX1 negative cell (arrowhead). (**d-f**) Representative immunohistochemistry images (n=3 per group) showing PRRX1 staining (Brown Chromogen, nuclear staining) in IPF fibroblast foci with (d) Vimentin (Red chromogen), (e) ACTA2 (Red chromogen) and (f) CD45 (Red chromogen). Note that PRRX1 positive cells were Vimentin positive (see black arrow in (d)) but only some were ACTA2 positive (arrow and dashed box in (e)). PRRX1$^{pos}$ ACTA2$^{neg}$ cells were also present (see arrowhead in (e)). All PRRX1$^{pos}$ cell populations were also CD45 negative (f). Insert in (d): high magnification of the black dashed box in the main left panel showing a double positive PRRX1 (brown nuclear staining, see black arrow) and Vimentin (red cytoplasmic staining) cells. Note that the epithelium is negative for both markers. Panel in (e): high magnification of the black dashed box in the main right panel showing a double positive PRRX1 (brown nuclear staining, see black arrow) and ACTA2 (red cytoplasmic staining) cell. PRRX1$^{pos}$ ACTA2$^{neg}$ cells were also present (see arrowhead). Right panel in (f): high magnification of the black dashed box in the main left bottom panel showing a PRRX1 (brown nuclear staining) positive but CD45 negative cell (arrow) and a CD45$^{pos}$ (red cytoplasmic staining) positive but PRRX1$^{neg}$ cell (arrowhead). Nuclei were counterstained with hematoxylin in all panels. *Abbreviations: Vim (Vimentin), Pos (positive), neg (negative).* (Scale bar: 50µm in (a-b), 80µm in (c) and 25µm in high magnification (a and c); 80µm in (d-f) and 40µm in high magnification (d-f)).

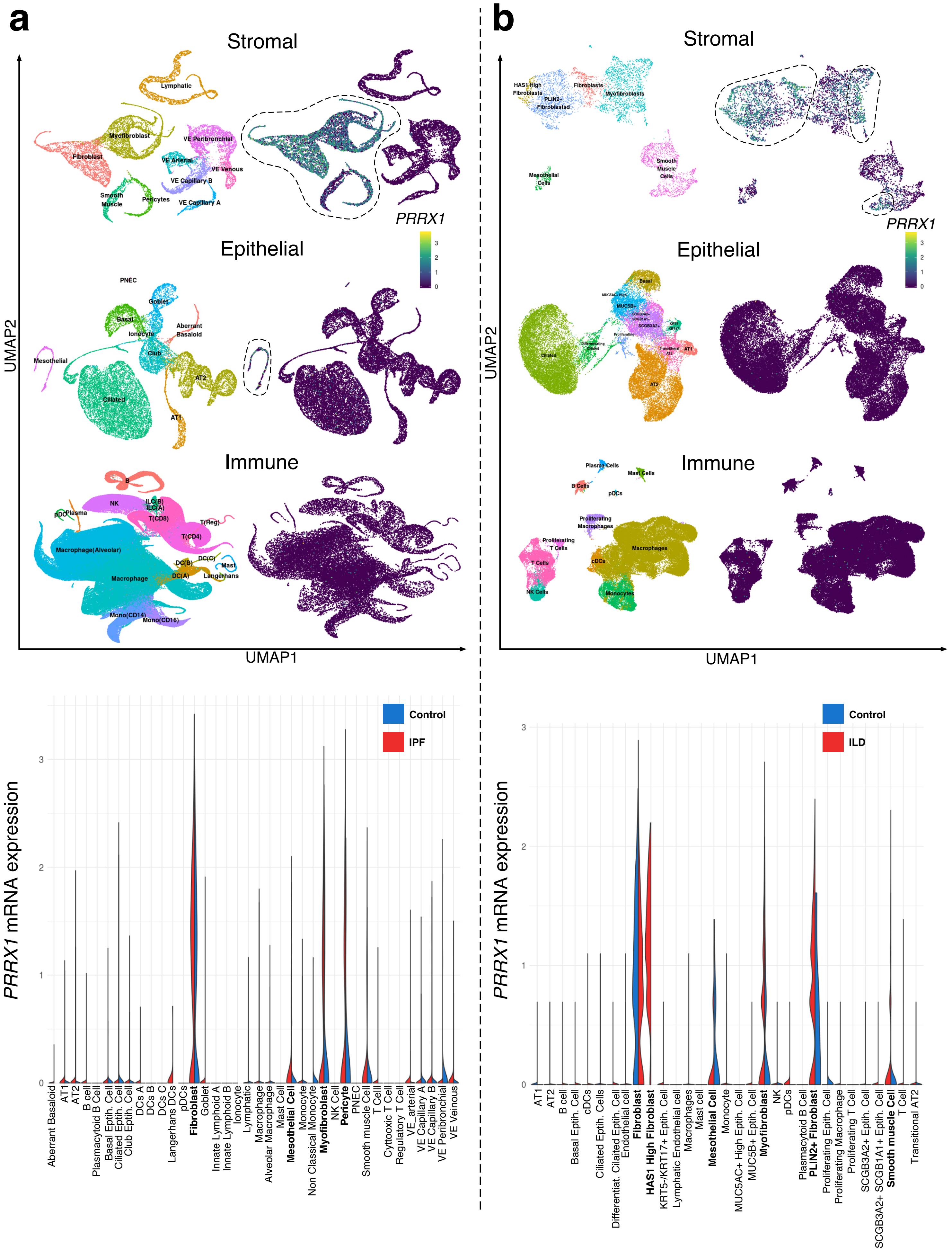


Supplemental Figure 2

**<u>Supplemental Figure S2 :</u> *PRRX1* expression profiles at single cell resolution using the "IPF Cell Atlas" web database (http://ipfcellatlas.com/).**

(**a-b**) UMAP plots describing the distribution of *PRRX1* expressing cells in different lung cell lineage clusters were drawn with UMAP Explorer (upper part) using (a) Kropski's and (b) Misharin's datasets. The labeling of each cell cluster (including all samples) are shown on the left and *PRRX1* relative expression in those clusters is shown on the right. In the lower part, violin plots visualizing *PRRX1* mRNA expression in each cell type stratified by disease states (control lung cell types in blue (a-b) and IPF (a) or ILD (b) ones in red) were drawn with Gene Explorer using (a) Kropski's dataset and in (b) Misharin's dataset. Note that *PRRX1* mRNA expression was associated with stromal clusters in both datasets (dashed lines in the upper part and labels in bold font in the lower part). *Abbreviations: AT1 (alveolar type 1 epithelial cell), AT2 (alveolar type 2 epithelial cell), Epith. (epithelial), B Cell (B lymphocyte), T Cell (T lymphocyte), DCs (dendritic cells), pDCs (plasmacytoid dendritic cells), NK Cell (Natural Killer cell), PNEC (pulmonary neuroendocrine cell), VE (vascular endothelium), ILD (interstitial lung disease).*

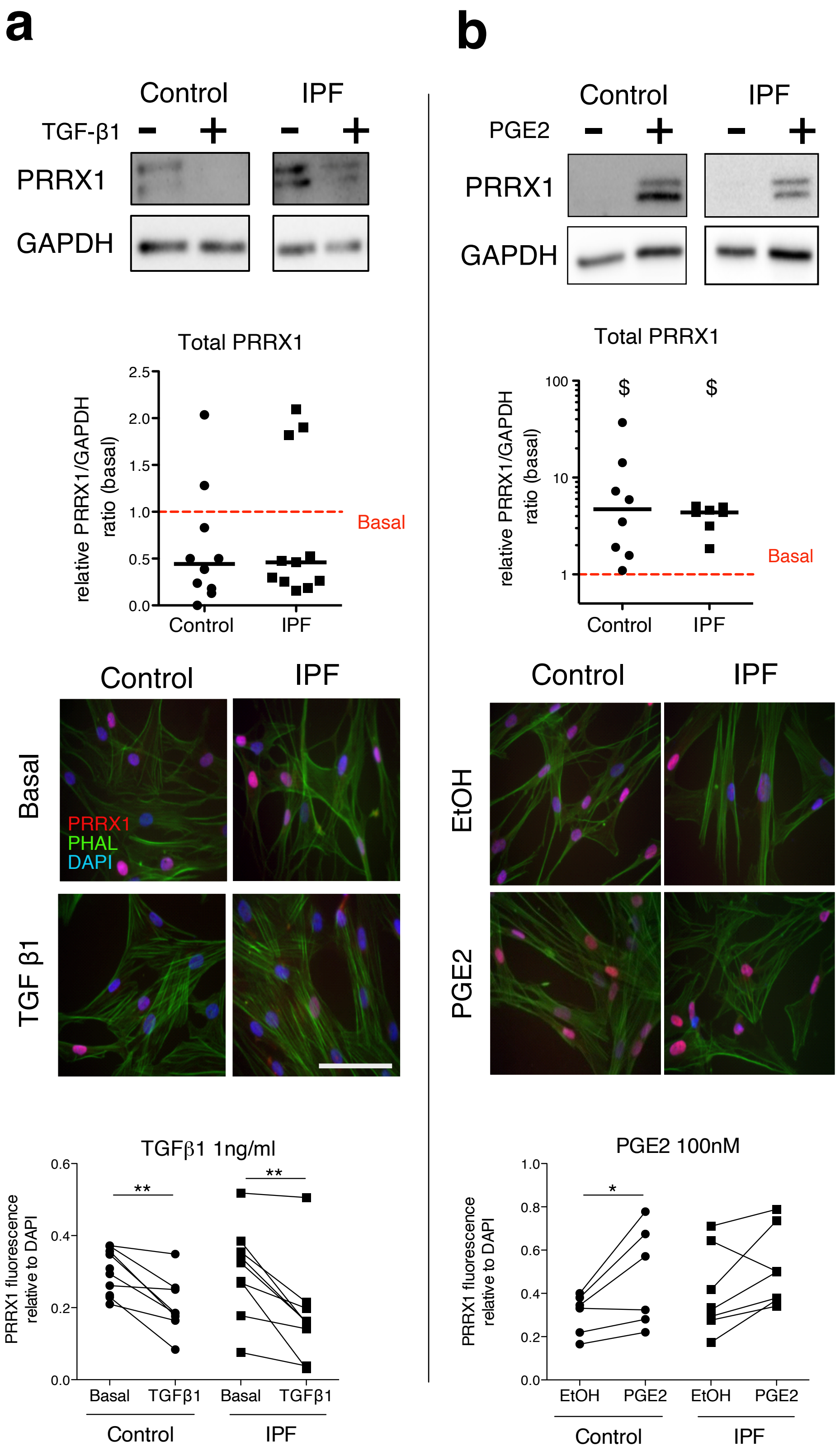


Supplemental Figure 3

**Supplemental Figure S3: Regulation of PRRX1 protein expression by growth factors as assayed by immunoblots and immunofluorescence.**

(**a**) Upper part: immunoblot showing PRRX1 expression in control and IPF primary Human lung fibroblasts stimulated 48h with TGF-β1. GAPDH was used as loading control. Middle panel: quantification of PRRX1 relative expression to GAPDH in control (circle) and IPF (square) lung fibroblasts stimulated for 48h with TGF-β1 compared to basal condition (red dashed line). Lower part: representative immunofluorescence images (n=7 per group) of PRRX1 expression (red) in control and IPF lung fibroblasts at basal level or stimulated for 48h with TGF-β1 (1ng/ml). DAPI was used as loading control. Actin fibers (green) were stained with Phalloidin. The quantification of PRRX1 relative expression to DAPI in control (circle) and IPF (square) lung fibroblasts at basal level or stimulated 48h with TGF-β1 is displayed as dot plot with median on the lower panel. (**b**) Upper part: immunoblot showing PRRX1 expression in control and IPF primary Human lung fibroblasts stimulated 48h with PGE2. GAPDH was used as loading control. Middle panel: quantification of PRRX1 relative expression to GAPDH in control (circle) and IPF (square) lung fibroblasts stimulated for 48h with PGE2 compared to basal condition (red dashed line). Lower part: representative immunofluorescence images (n=7 per group) of PRRX1 expression (red) in control and IPF lung fibroblasts stimulated for 48h with EtOH or PGE2 (100nM). DAPI was used as loading control. Actin fibers (green) were stained with Phalloidin. The quantification of PRRX1 relative expression to DAPI in control (circle) and IPF (square) lung fibroblasts stimulated for 48h with EtOH or PGE2 is displayed as dot plot on the bottom. (**c**) Upper part: immunoblot showing PRRX1 expression in control and IPF primary Human lung fibroblasts stimulated 48h with FGF2. GAPDH was used as loading control. Middle panel: quantification of PRRX1 relative expression to GAPDH in control (circle) and IPF (square) lung fibroblasts stimulated for 48h with FGF2 compared to basal condition (red dashed line). Lower part: representative immunofluorescence images (n=7 per group) of PRRX1 expression (red) in control and IPF lung fibroblasts stimulated for 48h with Heparin or FGF2 (20ng/ml). DAPI was used as loading control. Actin fibers (green) were stained with Phalloidin. The quantification of PRRX1 relative expression to DAPI in control (circle) and IPF (square) lung fibroblasts stimulated 48h with Heparin or FGF2 is displayed as dot plot with median on the bottom (Scale bar: 50μm), (*Abbreviations: Phalloidin (PHAL)).* Mann Whitney U test, *$p \leq 0.05$, **$p \leq 0.01$, Wilcoxon signed-rank test \$ $p \leq 0.05$

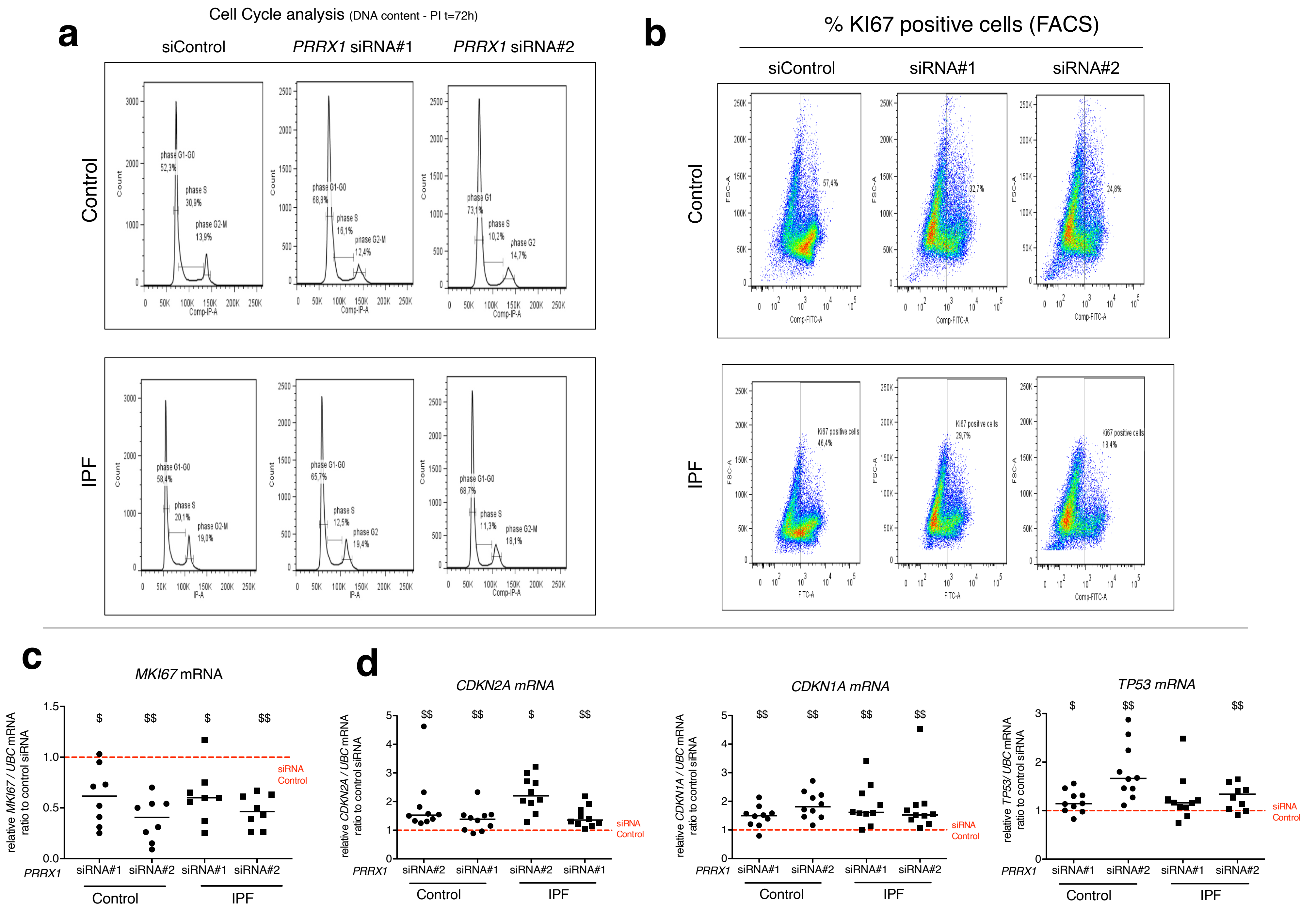


Supplemental Figure 4

**<u>Supplemental Figure S4</u>: PRRX1 inhibition impacted fibroblast proliferation.**

(**a**) Representative flow cytometry analysis of DNA content (n=5 per group) of control (upper panel) and IPF (lower panel) lung fibroblasts stimulated 72h with FCS 10% and treated with *PRRX1* siRNA (#1 or #2) or siControl. (**b**) Representative flow cytometry analysis of Ki67positive cells (n=5 per group) of control (top panel) and IPF (bottom panel) lung fibroblasts stimulated 72h with FCS 10% and treated with *PRRX1* siRNA (#1 or #2) or siControl. (**c**) Dot plots with median showing the mRNA expression of *MKI67* mRNA relative to the siControl condition (red dashed line), in control (black circle) and IPF (black square) lung fibroblasts treated for 48h with *PRRX1* siRNA (#1 or #2). (**d**) Dot plots with median showing the mRNA expression of *CDKN2A, CDKN1A* and *TP53* relative to the siControl condition in control (circle) and IPF (square) lung fibroblasts stimulated 72h with FCS 10% and treated with PRRX1 siRNA (#1 or #2). (*Abbreviations: control siRNA sequence (siControl); Fluorescence-activated cell sorting (FACS)*). Wilcoxon signed-rank test \$ $p \leq 0.05$ \$\$ $p \leq 0.01$

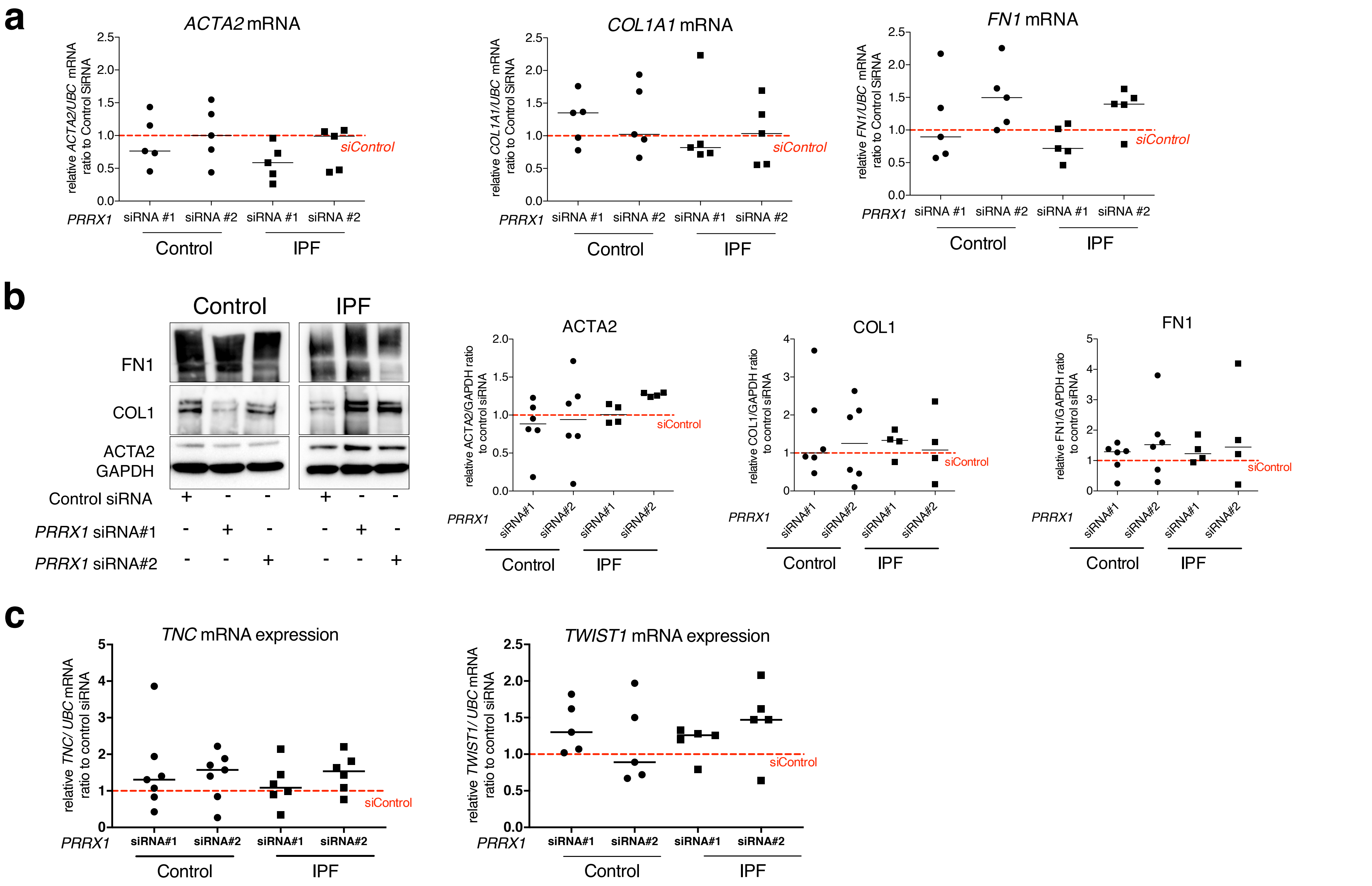


Supplemental Figure 5

**<u>Supplemental Figure S5</u>: PRRX1 inhibition did not modulate the basal expression of myofibroblastic markers in control and IPF fibroblasts.**

(**a**) Dot plots with median showing the mRNA expression of *ACTA2*, *COL1A1* and *FN1* relative to the siControl condition (red dashed line), in control (circle) and IPF (square) lung fibroblasts treated for 48h with *PRRX1* siRNA (#1 or #2). (**b**) Immunoblot showing FN1, COL1 and ACTA2 expression in control and IPF lung fibroblasts treated for 48h with siControl or PRRX1 siRNA (#1 or #2). GAPDH was used as loading control. Right part: quantification of COL1,FN1 and ACTA2 relative expression to GAPDH in control (circle) and IPF (square) lung fibroblasts treated for 48h with PRRX1 siRNA (#1 or #2) relative to the siControl condition (red dashed line). (**c**) Dot plots with median showing the mRNA expression of *TNC* and *TWIST1* relative to the siControl condition (red dashed line), in control (circle) and IPF (square) lung fibroblasts treated for 48h with *PRRX1* siRNA (#1 or #2). (*Abbreviations: control siRNA sequence (siControl)).*

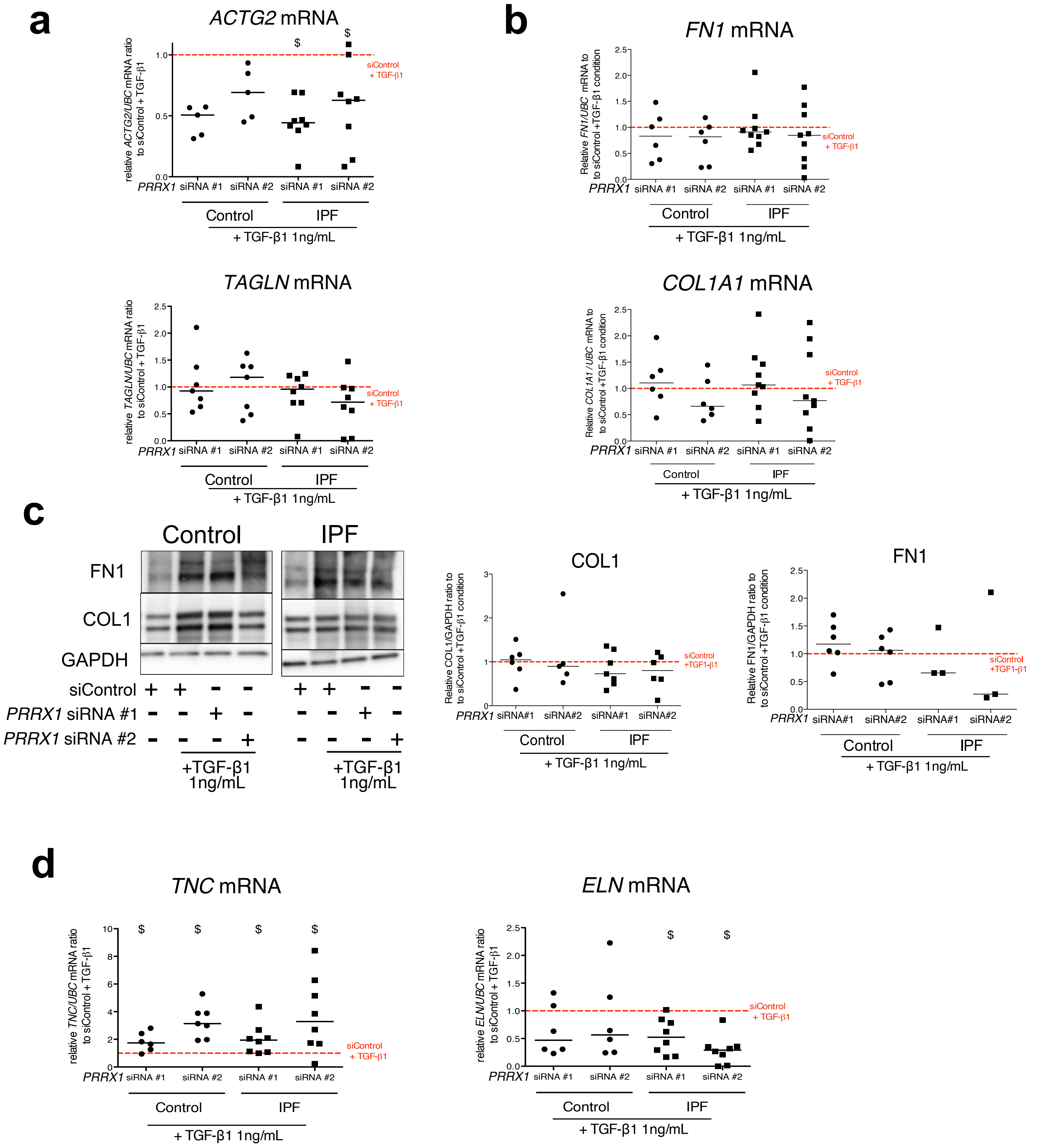


Supplemental Figure 6

**Supplemental Figure S6: Effect of PRRX1 inhibition upon the expression of myofibroblastic markers after TGF-β1 stimulation in control and IPF fibroblasts.**

(**a**) Dot plots with median showing the mRNA expression of *ACTG2* and *TAGLN* relative to the siControl +TGF-β1 condition (red dashed line), in control (circle) and IPF (square) lung fibroblasts treated with TGF-β1 and PRRX1 siRNA (#1 or #2). (**b**) Dot plots with median showing the mRNA expression of *FN1* and *COL1A1* relative to the siControl +TGF-β1 condition (red dashed line), in control (circle) and IPF (square) lung fibroblasts treated with TGF-β1 with *PRRX1* siRNA (#1 or #2). (**c**) Immunoblot showing FN1 and COL1 expression in control and IPF lung fibroblasts treated with control siRNA in absence or presence of TGF-β1 or with *PRRX1* siRNA (#1 or #2) and TGF-β1. GAPDH was used as loading control. The quantification of COL1 and FN1 expressions relative to GAPDH in control (circle) and IPF (square) lung fibroblasts treated with control or *PRRX1* siRNA (#1 and #2) in presence of TGF-β1 relative to the siControl + TGF-β1 condition (red dashed line), are displayed as dot plot with median in the right part of the panel. (**d**) Left part: dot plots with median showing the mRNA expression of *TNC* relative to the siControl + TGF-β1 condition (red dashed line), in control (circle) and IPF (square) lung fibroblasts treated with TGF-β1 and *PRRX1* siRNA (#1 or #2). Right part: dot plots with median showing the mRNA expression of *ELN* relative to the siControl +TGF-β1 condition (red dashed line), in control (circle) and IPF (square) lung fibroblasts treated with TGF-β1 and *PRRX1* siRNA (#1 or #2). (*Abbreviations: TSS (transcription starting site); IgG (Immunoglobulin); PRE (PRRX1 responses element); SRE (SRF response element), control siRNA sequence (siControl)).* Wilcoxon signed-rank test, $ $p \leq 0.05$

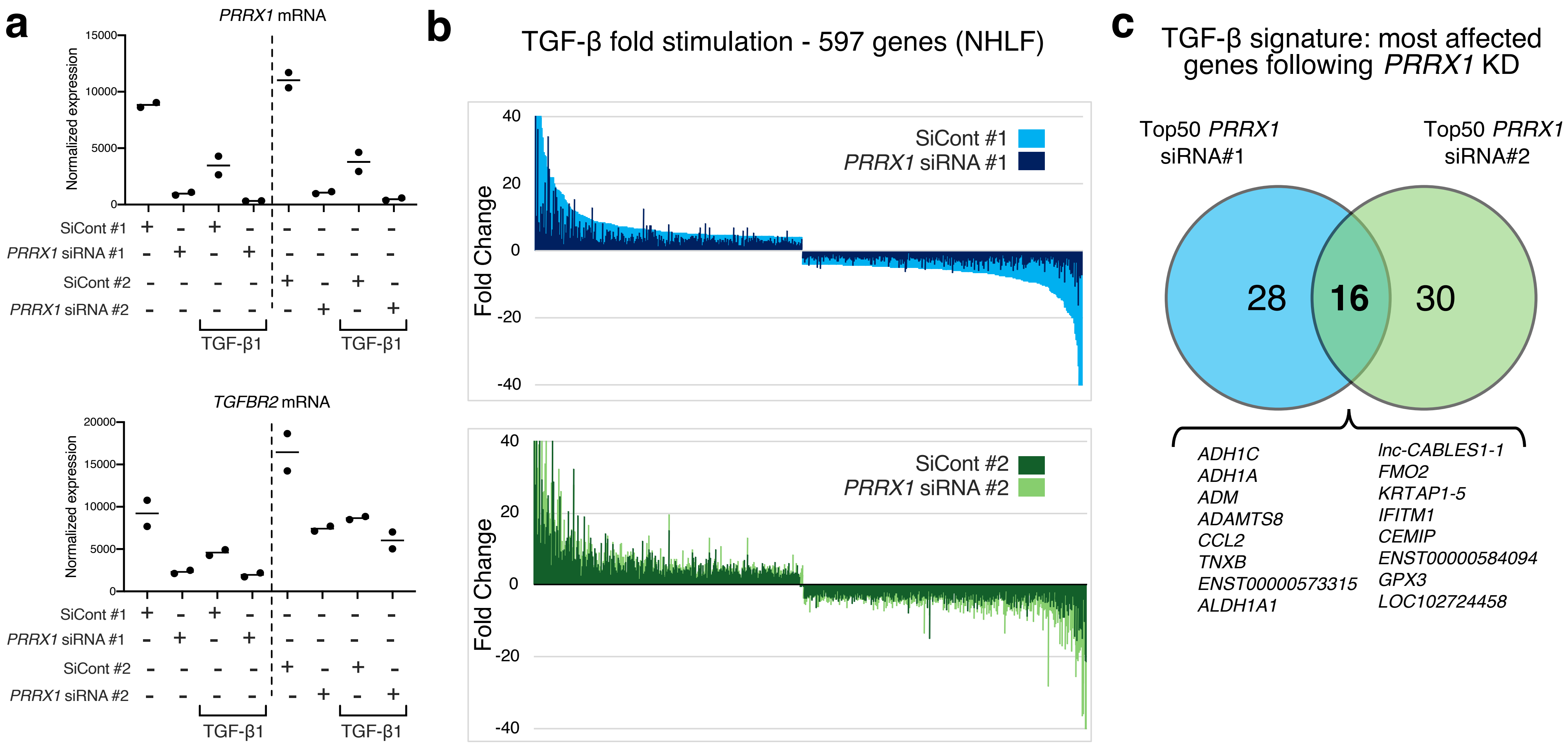


Supplemental Figure 7

**Supplemental Figure S7: *PRRX1* knock-down globally impacted TGF-β pathway response in NHLF.**

NHLF lung fibroblasts were first treated with control siRNA (#1 or #2) or with *PRRX1* siRNA (#1 or #2) for 48h in the absence or presence of TGF-β for an additional 48h. RNA was extracted and analyzed at 96h using whole genome microarrays (n=2). **(a)** Dot blots showing normalized mean expression of *PRRX1* and *TGFBR2* in the different conditions (n=2). **(b)** Box plots showing the attenuation of the TGF-β response by *PRRX1* siRNAs on the 597 most modulated genes by TGF-β in control conditions (threshold: log2 fold change >2, adj. P Value<0.05, see supplemental table 2). Average attenuation of the TGF-β response is 44.4 and 20% with siRNA#1 and #2, respectively. **(c)** Venn diagram showing the most affected genes following *PRRX1* KD within the TGF-β signature in NHLF.

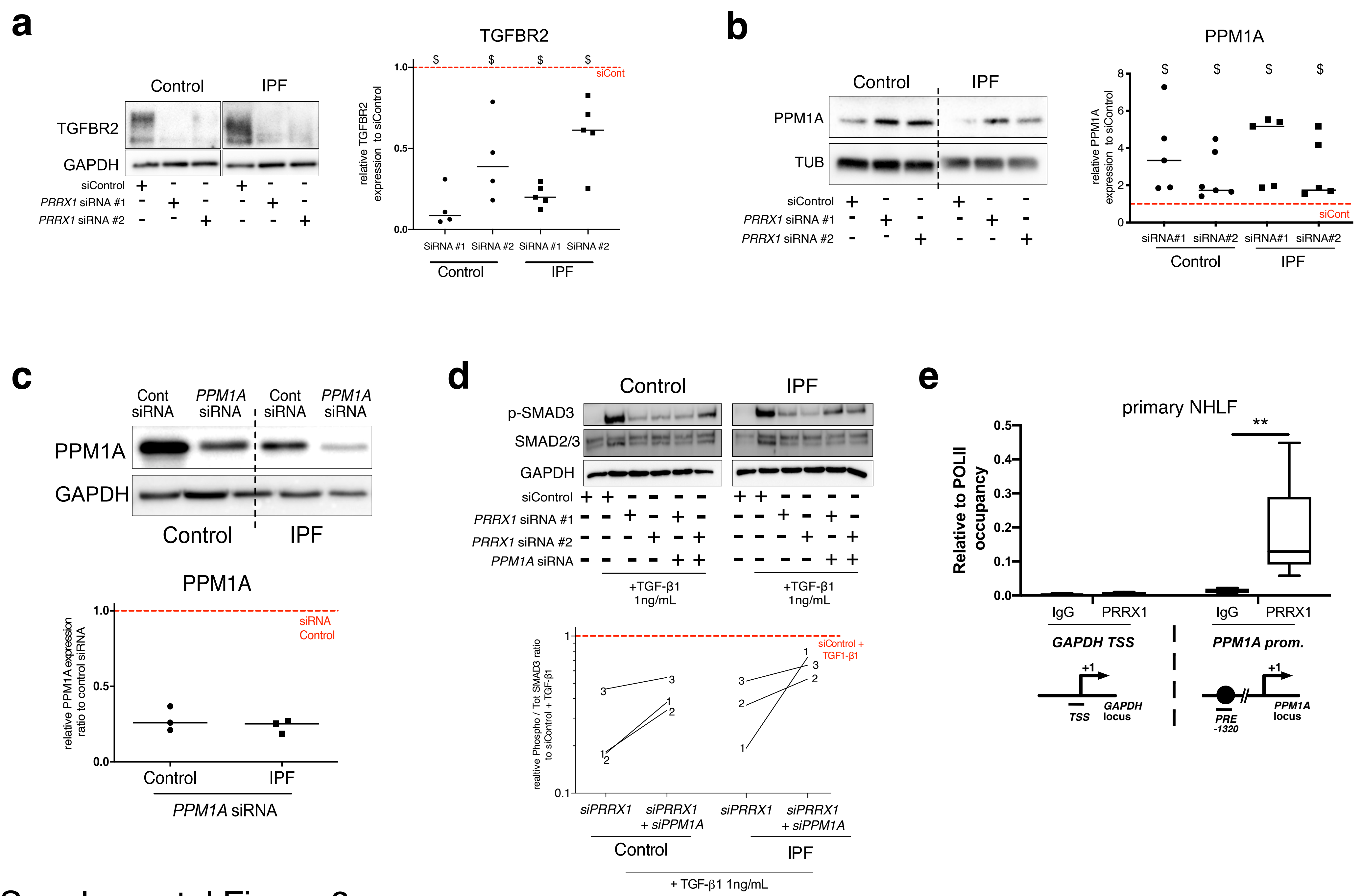


Supplemental Figure 8

**<u>Supplemental Figure S8</u>: PRRX1 TFs modulate TGF-β / SMAD signalling cascade by regulating TGFBR2 and PPM1A expressions**

(**a-b**) Representative immunoblot showing TGFBR2 (a) or PPM1A (b) expression in control and IPF fibroblasts treated for 48h with *PRRX1* siRNA. The quantification of TGFBR2 (a) or PPM1A (b) expression relative to loading controls in control and IPF fibroblasts treated for 48h with *PRRX1* siRNA is displayed as dot plot with median. (**c**) Immunoblot showing PPM1A expression in control and IPF primary Human lung fibroblasts treated 48h with *PPM1A* siRNA or siControl. GAPDH was used as loading control. Quantification of PPM1A (n=3) expression relative to GAPDH in control (circle) and IPF (square) lung fibroblasts treated 48h with *PPMA* siRNA compared to the siControl condition (red dashed line). (**d**) Representative immunoblot (n=3) showing phospho-SMAD2/3 and SMAD2/3 expression in control and IPF lung fibroblasts treated 30 minutes with TGF-β1 after 48h transfection with *PRRX1* siRNA (#1 or #2) and *PPM1A* siRNA. The quantification of phospho-SMAD2/3 and SMAD2/3 expression relative to GAPDH in control (circle) and IPF (square) lung fibroblasts treated for 30 minutes with TGF-β1 after 48h transfection with *PRRX1* siRNA (#1 or #2) and *PPM1A* siRNA relative to the siControl + TGF-β1 condition (red dashed line), is displayed as connected line in the right part of the panel (n=3). (**e**) ChIP analysis for recruitment of PRRX1 at the promoter of *GAPDH* and *PPM1A* genes in NHLF (n=5). An unrelated control IgG was used as negative control. The results are expressed relative to RNA POL-II occupancy of a given locus and displayed as boxes with median and min to max. The schematic diagrams of the different loci are showing the PRE or SRE element positions relative to the TSS of the corresponding gene. The PCR amplified regions are underscored. (*Abbreviations: control siRNA sequence (siControl), TSS (transcription starting site); IgG (Immunoglobulin); PRE (PRRX1 responses element); SRE (SRF response element), control siRNA sequence (siControl)).* Wilcoxon signed-rank test, $ $p \le 0.05$, Kruskal-Wallis test with Dunns post-test, ** $p \le 0.01$.

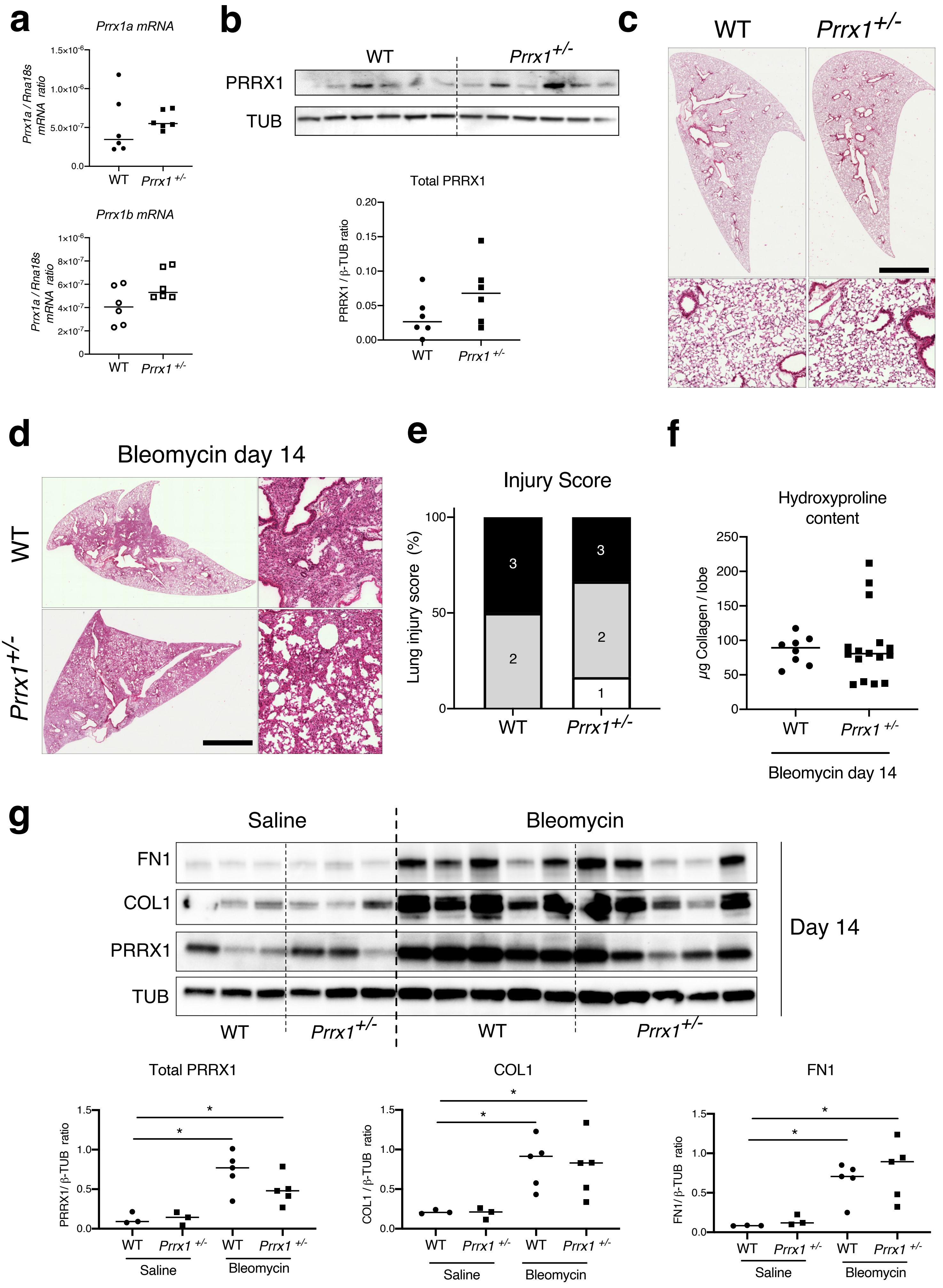


Supplemental Figure 9

**Supplemental Figure S9: lack of haploinsufficiency and lung fibrosis reduction in *Prrx1*$^{+/-}$ heterozygous mice.**

(**a**) Dot plots with median showing the mRNA expression of *Prrx1a* (black) and *Prrx1b* (white) isoforms in naive wild type (circle) and *Prrx1*$^{+/-}$ (square) mice. (**b**) Immunoblot showing PRRX1 expression in naive wild type mice and *Prrx1*$^{+/-}$ mice. TUB was used as loading control. The quantification of PRRX1 expression relative to TUB at day 14 in naive wild type (circle) and *Prrx1*$^{+/-}$ (square) mice is displayed as dot plot with median on the lower panel. Note the lack of *Prrx1* haploinsufficiency at both mRNA and protein levels in *Prrx1* heterozygous mice. No difference in COL1, FN1 and ACTA2 levels were also observed in *Prrx1*$^{+/-}$ mice compared to wild type ones (data not shown). (**c**) Representative hematoxylin – eosin staining images (n=6 per group) showing no histological differences between naive wild type and *Prrx1*$^{+/-}$ lungs. (**d**) Representative hematoxylin – eosin staining images (n=7 at least per group) showing no histological differences between wild type and *Prrx1*$^{+/-}$ lungs at day 14 after bleomycin insult. (**e**) Injury score of wild type and *Prrx1*$^{+/-}$ mice treated bleomycin at day 14 (at least n=7 per group). (**f**) Dot Plot with median showing the Collagen lung content (µg per lobe) as measured by hydroxyproline in wild type and *Prrx1*$^{+/-}$ mice treated with bleomycin at day 14. (**g**) Representative immunoblot showing FN1, COL1 and PRRX1 expression in wild type and *Prrx1*$^{+/-}$ mice treated with saline or bleomycin at day 14. TUB was used as loading control. Lower panels: quantification of PRRX1, COL1 and FN1 relative expression to TUB in wild type and *Prrx1*$^{+/-}$ mice treated with saline or bleomycin at day 14. Note that lung fibrosis is not reduced in *Prrx1* heterozygous mice compared to wild type animals after bleomycin treatment at day 14 (D-G). (Scale bar in D: 80µm in low magnification images and 40µm in high magnification ones); *(Abbreviations: Wild type (WT), $\beta$-TUB (beta-Tubulin)).* Kruskal-Wallis test with Dunns post-test, *p≤0.05.

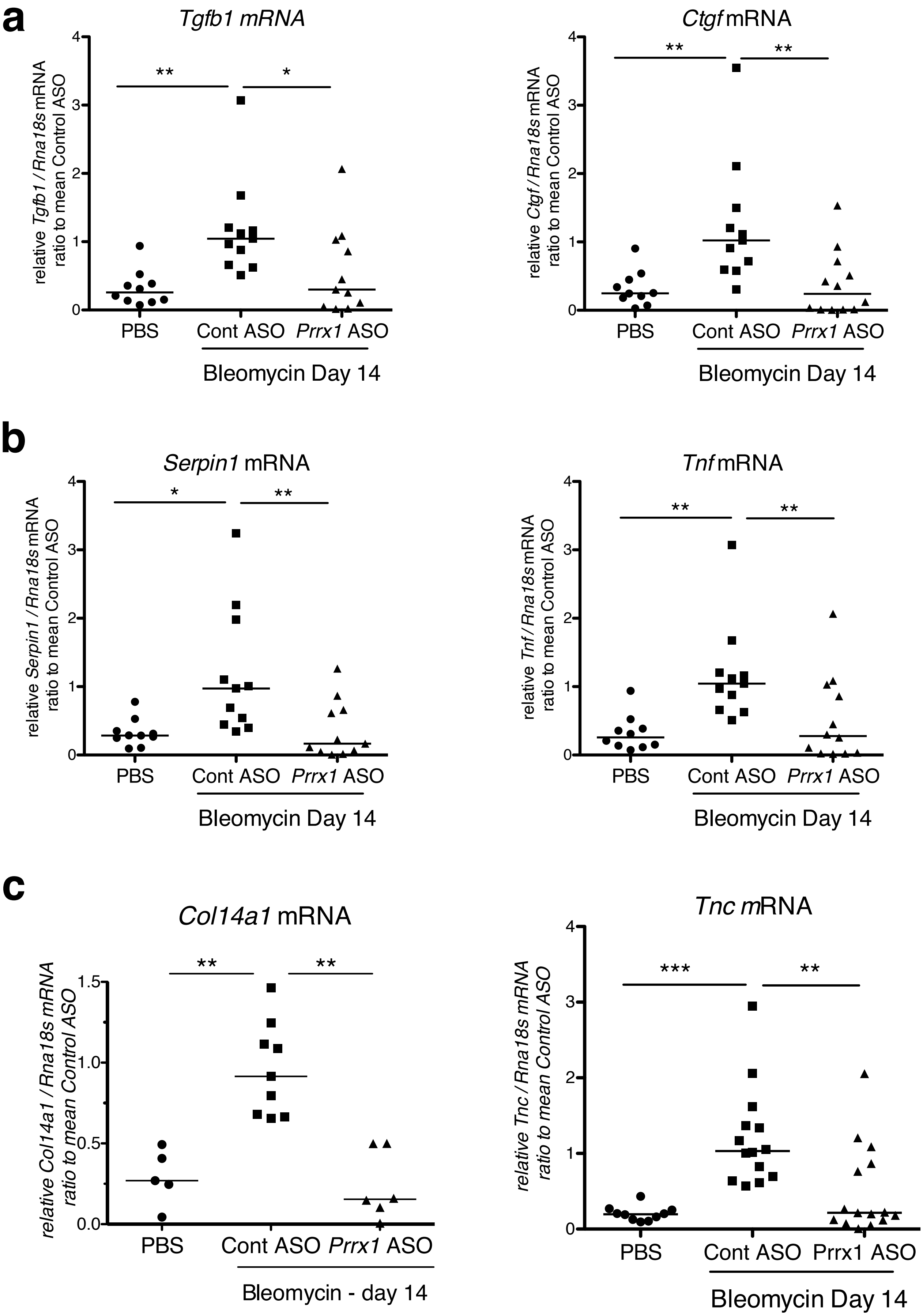


Supplemental Figure 10

**<u>Supplemental Figure S10:</u> PRRX1 inhibition decreases fibrosis and inflammatory markers in bleomycin mice.**

(**a**) Dot plots with median showing the mRNA expression of fibrotic markers such as *Tgfb1* and *Ctgf* at day 14 in PBS mice (circle) and bleomycin mice treated with Control ASO (square) or *Prrx1* ASO (triangle). (**b**) Dot plots with median showing the mRNA expression of inflammatory markers such as *Serpin1* and *Tnf* at day 14 in PBS mice (circle) and bleomycin mice treated with Control ASO (square) or *Prrx1* ASO (triangle). (**c**) Dot plots with median showing the mRNA expression of ECM components such as *Col14a1* and *Tnc* at day 14 in PBS mice (circle) and bleomycin mice treated with Control ASO (square) or *Prrx1* ASO (triangle). *(Abbreviations: Control (Cont), Antisense oligonucleotide (ASO))*. Kruskal-Wallis test with Dunns post-test, *p≤0.05, **p≤0.01, ***p≤0.001

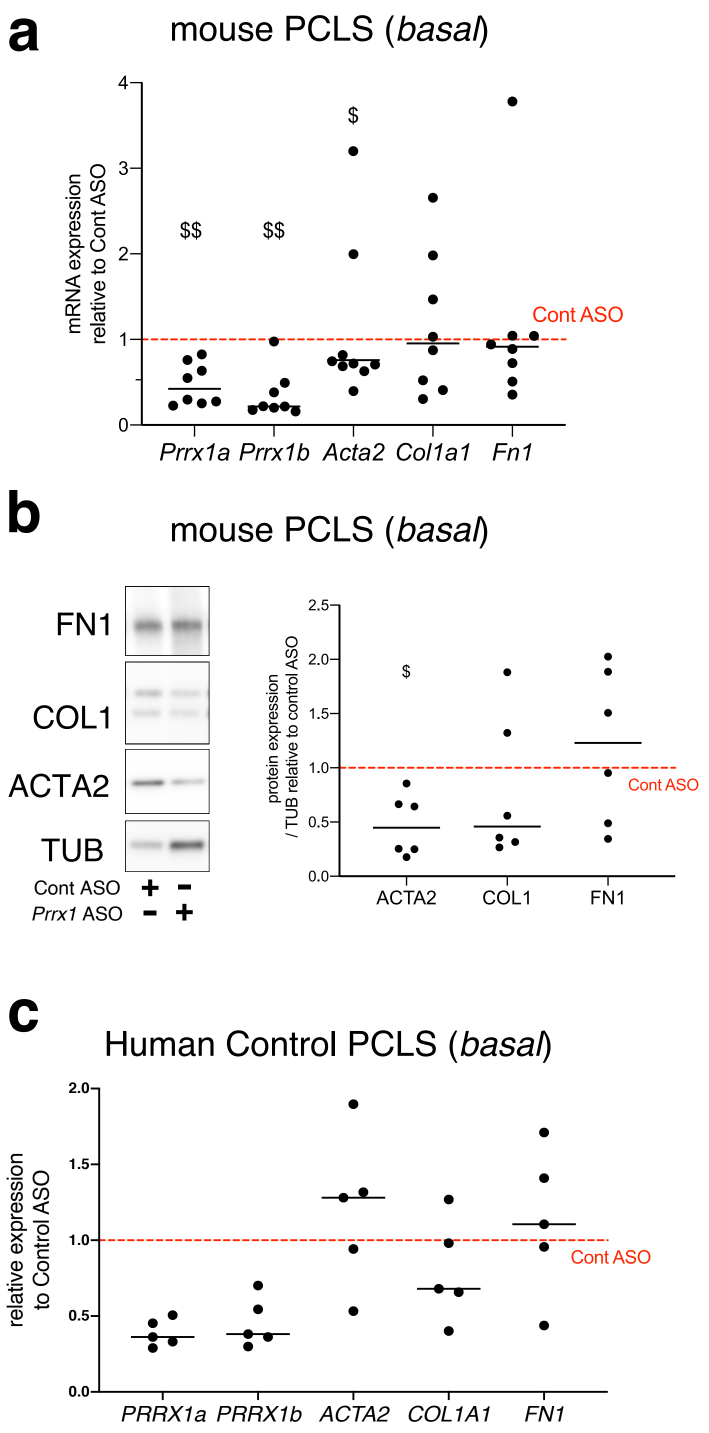


### Supplemental Figure 11

**<u>Supplemental Figure S11</u>: effect of *PRRX1* ASO on the expression of myofibroblastic differentiation markers in control mouse and Human PCLS at basal condition.**

(**a**) Dot plots with median showing the relative mRNA expression of *Prrx1a, Prrx1b, Acta2, Col1a1* and *Fn1* in mouse PLCS treated with *Prrx1* ASO alone relative to control ASO (red dashed line). Note that *Prrx1a*, *Prrx1b and Acta2* mRNA levels were decreased in presence of *Prrx1* ASO compared to control ASO. (**b**) Immunoblot showing FN1, COL1 and ACTA2 expression in mouse PCLS treated with control or *Prrx1* ASO. TUB was used as loading control. Right part: quantification of COL1, FN1 and ACTA2 relative expression to TUB in mouse PCLS treated with *Prrx1* ASO (n=6) relative to control ASO (red dashed line). (**c**) Dot plots with median showing the relative mRNA expression of *PRRX1A, PRRX1B, ACTA2, COL1A1* and *FN1* in control Human PLCS (n=5) treated with *PRRX1* ASO alone relative to control ASO (red dashed line). Note that *PRRX1a*, *PRRX1b* mRNA levels were effectively downregulated in presence of *PRRX1* ASO compared to control ASO. *Abbreviations: Precision-Cut Lung slices (PCLS), Control (Cont), Antisense oligonucleotide (ASO)*. Wilcoxon signed-rank test, \$ $p \leq 0.05$ \$\$ $p \leq 0.01$

#### Supplemental Material and Methods

##### Analysis of publicly available control and IPF lung RNA microarray and single cell datasets

Analysis of curated NCBI GEO dataSet GDS1252 (GSE2052 [1]), GDS4279 (GSE24206 [2]) and GDS3951 (GSE21411 [3]) from control and IPF whole lung transcriptomic data was performed using NCBI "Compare 2 sets of samples" Geo Dataset analysis tool (https://www.ncbi.nlm.nih.gov/geo/info/datasets.html). The list of upregulated genes in IPF samples compared to control ones in the three transcriptomic datasets was established by generating a Venn's diagram (Venny 2.1 tool; http://bioinfogp.cnb.csic.es/tools/venny/). Gene annotation analysis (Protein Class) was then performed with PANTHER [4]. Single-cell transcriptomic analysis of lungs from transplant donors or recipients with pulmonary fibrosis lung was performed using the dataset available at www.nupulmonary.org/resources/ [5] and www.ipfcellatlas.com [6].

##### Gene expression profiling

Integrity of total RNAs was assessed using an Agilent BioAnalyser 2100 (Agilent Technologies) (RIN>9). 150-200 ng of RNA samples (with RIN > 9) were labeled with Cy3 dye using the low RNA input QuickAmp kit (ref 5190-2305, Agilent) as recommended by the supplier. After fragmentation step, 600 ng of Cy3-labeled cRNA probes were hybridized on 8x60K v3 high density SurePrint G3 gene expression human Agilent microarrays (G4851C, Agilent) with respect to the manufacter protocol. Statistical analysis and Biological Theme Analysis. Microarray data analyses were performed using the Bioconductor package limma. Briefly, data were normalized using the quantile method. No background subtraction was performed. Replicated probes were averaged after normalization and control probes removed. Statistical significance was assessed using the limma moderated t-statistic. All P-values were adjusted for multiple testing using the Benjamini-Hochberg procedure. Differentially expressed genes were selected based on an adjusted p-value below 0.05. Enrichment in biological themes (Molecular function, Upstream regulators and canonical pathways) were performed using Ingenuity Pathway Analysis software (http://www.ingenuity.com/). Expression dataset supporting the Figure 5F and the supplemental Figure 12 of this study has been deposited in the Gene Expression Omnibus GSE161364: Impact of PRRX1 knockdown on the transcriptome of normal human lung fibroblasts (NHLF) in the presence or the absence of TGF-β (8 experimental conditions performed in duplicates).

##### Cell culture experiments.

Primary Human lung fibroblasts from control patients were derived from Human lung explants as previously described [7]. Primary HLFs were cultured with Dulbecco's modified Eagle's medium (Thermo-Fisher Scientific, Waltham, USA) supplemented with 10% fetal calf serum (FCS) and antibiotics and used at passage 4 to 6 as previously described [7]. Primary lung fibroblasts were treated for 48 h with TGF-β1 (1ng/ml) (Peprotech, Neuilly sur seine, France), PGE2 (100nM, Sigma, Saint-Louis, USA), NS398 (10μM, Sigma Aldrich, Saint-Louis, USA) and Fasudil (35μM, Sigma Aldrich, Saint-Louis, USA).

*PRRX1* siRNA mediated inhibition: primary HLFs were transfected at 50 to 60% confluency in 6- or 12-well plates. All transfection reactions were performed with, lipofectamine RNAimax (ThermoFisher Scientific, Waltham, USA) reagent transfection, in accordance with the manufacturer's instruction as described previously [7]. To suppress *PRRX1* endogenous expression, lung fibroblasts were transfected with two independent sequences targeting both *PRRX1a* and *PRRX1b* mRNA: siRNA#1, 5'-AGAAAGCAGCGAAGGAAUA-3' (GE Health Care-Dharmacon, Lafayette, USA,) & siRNA#2, 5'-GCGAAGGAAUAGGACAACCUUCAAU-3' (Fisher Scientific, Waltham, USA). siRNA Negative Control Low GC (12935-200, Invitrogen, Carlsbad, USA) was used as a negative control (siControl). A final concentration of 30 nM for each siRNA was used. A *PPM1A* siRNA smartpool was used to knock down *PPM1A* in lung fibroblasts (D-009574, GE Health Care-Dharmacon, Lafayette, USA).

Fibroblast myofibroblastic differentiation: serum-starved fibroblasts for 16h were cultured for 48 hours with *PRRX1* siRNA sequences or negative control, then treated during 48 hours with TGF-β1 (1ng/ml). Cells were harvested and assayed for further mRNA and Western Blot analysis (as described below).

Fibroblast proliferation: was assayed 72h after *PRRX1* siRNA transfection. The proliferation rate was measured by CyQUANT NF Cell Proliferation Assays (Thermofisher, Waltham, USA) which quantify cellular DNA content using a microplate reader (Infinite® 200 PRO,TECAN, Männedorf, Suisse). Cytotoxicity was assayed by trypan blue exclusion assay.

Stiffness experiments: Cells were plated during 48h on substrate with defined elastic modulo (EM) which correspond to the matrix rigidity: 1,5kPa (soft matrix, 81291), 28 kPa (stiff matrix, 81191) or in the GPa range (glass, 81158) (ibidi, GmbH, Planegg, Germany). Coating with Fibronectin (Human plasma, Sigma Aldrich) was performed according to plate's manufacturer instructions (ibidi, GmbH, Planegg, Germany)

Fibroblast-derived matrix experiments: thick matrices were produced from primary Human control and IPF lung fibroblast using an adapted protocol from R. Castello-Crós & colleagues [8]. Control primary lung fibroblasts were next seeded on this 3D matrices and treated with Imatinib (10μg/ml), Nintedanib (10nM) or PDGFR V inhibitor (10nM) (all from Selleckchem, Boston USA) in DMEM medium supplemented with 2% of FCS. 48h after treatment, cells were harvested for mRNA analysis (as described below).

mRNA analysis: Total mRNA was extracted from Human lung samples and from cultured fibroblasts, and cDNA was obtained by standard techniques as described [7]. *Ubiquitin C* (*UBC*) mRNA was used for normalization. Primers are reported in Table S3 in supplemental material

Western blot analysis: Proteins were extracted from Human lung samples and from cultured fibroblast by standard techniques as described [7]. Western blotting was performed under denaturing and reducing conditions. Antibodies are reported in Table S4 in supplemental material.

Immunofluorescence, cells were cultured in serum-free medium on Permanox Lab-Tek Chamber Slides (Nunc, ThermoFisher Scientific, Waltham, USA) as described [9]. The primary antibody was PRRX1 (Rabbit polyclonal antibody HPA051084 from Sigma Aldrich). Anti-Rabbit 568 Alexa Fuor coupled antibodies were used as secondary antibody. Alexa-fluor 488 coupled Phalloidin was used to stain actin fibers and DNA was stained with DAPI (Invitrogen, Carlsbad, USA)

Chromatin immunoprecipitation, ChIP analyses were performed with Abcam's ChIP Kit –" One Step" (ab117138) following manufacturer instructions. Briefly, formaldehyde cross-linked chromatin from NHLF (primary control lung Human fibroblasts purchased from Lonza, Basel, Switzerland) was prepared as previously described [7]. Chromatin fragments (from about $1x10^6$ cells) were immunoprecipitated with antibodies against PRRX1 (Sigma Aldrich HPA051084), RNA POL-II and a control IgG (both from Abcam's "ChIP Kit - One Step"). The precipitated DNA was amplified by real-time PCR with primer sets designed to amplify PRRX1 response elements (PRE) in *CCNA2*, *CCNE2*, *MKI67* and *PPM1A* loci and *GAPDH* TSS (supplemental table 1). The results are expressed relative to RNA POL-II occupancy of a given locus. Putative PRE sequences in those different loci were first identified within the regulatory build / core promoter sequences (NHLF cells) available from Ensembl database [10] using FIMO (MEME Suite Programs [11] ) and LASAGNA-Search [12] tools with PRE consensus sequence (Jaspar database [13]).

**Culture of Mouse and Human Precision-Cut Lung Slices (PCLS) with Fibrosing Cocktail (FC)**

Mouse and Human PCLS were obtained from 6mm biopsy punch performed on agarose-filled lung samples. Mouse PLCS were obtained from 8-week-old C57BL/6 mice, while Human PCLS were prepared form lung samples obtained after cancer surgery, away from the tumor. Lung cylinders were next sliced (300µm) using a McIlwain Tissue Chopper and processed as previously described [14,15]. Briefly, Human PCLS were treated with FC for 48h in presence of *PRRX1* ASO (10µM) or control ASO (10µM) without transfection reagent. Mouse PCLS were first stimulated with FC for 48h then treated with either control or *PRRX1* ASO for additional 48h without transfection reagent. FC was prepared as previously described [15] and all the cytokines were purchased from PeproTech (Neuilly sur Seine, France). In basal condition, mouse PCLS or control Human PCLS were treated only with control ASO (10µM) for 48h. PCLS were harvested for mRNA, protein and histological studies using previously described methods [16]. Picrosirius staining and quantification were performed as described previously [17]

#### Flow Cytometry

FACS analysis was assayed after 72h siRNA PRRX1 transfection. Cells were harvested and prepared as previously described [18]. Cells were stained for propidium iodide (0.5 mg/ml) (Sigma, Saint-Louis, USA) and anti Ki-67 Alexa Fluor 488 (1/500) or isotype control IgG1 Alexa Fluor 488 (1/500) (Becton, Dickinson and Company (BD), Franklin Lakes, US). Samples were analyzed by flow cytometry with a FACSCanto™ II flow cytometer using the FACSDIVA software (Becton Dickinson, New Jersey, USA) for data acquisition (15,000–30,000 events) and the FlowJo, 9.9.6 software (FlowJo LLC, Ashland, Oregon, USA) for analysis.

#### *Immunohistochemistry and hydroxyproline assay.*

Immunohistochemistry, paraffin-embedded sections (*n* = 5 per group) were treated as described [9]. Antibodies are reported in Table S3 in supplemental material. To validate the specificity of immunostaining, antibodies were replaced by a matched control Isotype. All digital images of light microscopy were acquired with a DM400B microscope (Leica) equipped with a Leica DFC420 CDD camera and analyzed with Calopix software (TRIBVN).

Double immunohistochemistry for PRRX1 with Vimentin or ACTA2 were performed as previously described [19] using the Citrate "Heat Induced Epitope Retrieving"(HIER) microwave method between the consecutive stainings. Briefly, ACTA2 or Vimentine stainings were first revealed with "Dako Permanent Red Substrate-Chromogen" (Agilent) after using respectively Rabbit or Mouse "Histofine Simple Stain AP" (Nicheiri Biosciences) as secondary antibodies. PRRX1 staining was next revealed using "Dako Liquid DAB+ Substrate Chromogen System" (Agilent) after using Rabbit "Histofine Simple Stain MAX PO" as secondary antibody (Nicheiri Biosciences)

Hydroxyproline assay, total collagen and protein quantifications were performed using respectively the Quickzyme Biosciences (Leiden, The Netherlands) hydroxyproline assay and total protein assay kits from paraffine lung sections (15 sections of 10µm per samples), following manufacturer instructions.

#### SUPPLEMENTAL TABLES

**Supplemental Table S1: list of common up-regulated genes in all three IPF transcriptome database analyzed.**

| | | | |
|---|---|---|---|
| **UPREGULATED GENES ANNOTATED AS TRANSCRIPTION FACTORS (PANTHER GO)** | | | |
| Gene symbol | Name | Gene symbol | Name |
| FHL2 | four and a half LIM domains 2 | SMAD1 | SMAD family member 1 |
| NFATC4 | nuclear factor of activated T-cells 4 | SOX2 | SRY-box 2 |
| NR2F2 | nuclear receptor subfamily 2 group F member 2 | TP63 | tumor protein p63 |
| **PRRX1** | **paired related homeobox 1** | TRIM29 | tripartite motif containing 29 |
| RHBDD3 | rhomboid domain containing 3 | ZNF671 | zinc finger protein 671 |
| RUNX1 | runt related transcription factor 1 | ZNF862 | zinc finger protein 862 |
| **OTHER UPREGULATED GENES (alphabetical order in column)** | | | |
| Gene symbol | Name | Gene symbol | Name |
| AAK1 | AP2 associated kinase 1 | LPP | LIM domain containing preferred translocation partner in lipoma |
| ABCC5 | ATP binding cassette subfamily C member 5 | LRRC17 | leucine rich repeat containing 17 |
| ACTN1 | actinin alpha 1 | LTBP1 | latent transforming growth factor beta binding protein 1 |
| ADCY3 | adenylate cyclase 3 | MAGED4B | MAGE family member D4B |
| ALDH3A1 | aldehyde dehydrogenase 3 family member A1 | MAP1A | microtubule associated protein 1A |
| ANGPTL2 | angiopoietin like 2 | MAP4K4 | mitogen-activated protein kinase kinase kinase kinase 4 |
| ANKH | ANKH inorganic pyrophosphate transport regulator | MFAP2 | microfibrillar associated protein 2 |
| ANTXR1 | anthrax toxin receptor 1 | MIR4697HG | MIR4697 host gene |
| AP2A1 | adaptor related protein complex 2 alpha 1 subunit | MMP7 | matrix metallopeptidase 7 |
| ASB2 | ankyrin repeat and SOCS box containing 2 | MOXD1 | monooxygenase DBH like 1 |
| ASPN | asporin | MRVI1 | murine retrovirus integration site 1 homolog |
| BACE2 | beta-site APP-cleaving enzyme 2 | MXRA5 | matrix remodeling associated 5 |
| BAG4 | BCL2 associated athanogene 4 | MYOF | Myoferlin |
| BCAS4 | breast carcinoma amplified sequence 4 | NAV2 | neuron navigator 2 |
| BPIFB1 | BPI fold containing family B member 1 | NLGN2 | neuroligin 2 |
| BRD8 | bromodomain containing 8 | NSG1 | neuron specific gene family member 1 |
| C8orf44 | chromosome 8 open reading frame 44 | OGN | Osteoglycin |
| CADPS | calcium dependent secretion activator | OLFM1 | olfactomedin 1 |
| CAMK1D | calcium/calmodulin dependent protein kinase ID | ORAI2 | ORAI calcium release-activated calcium modulator 2 |
| CAPN5 | calpain 5 | OSBPL6 | oxysterol binding protein like 6 |
| CARM1 | coactivator associated arginine methyltransferase 1 | P2RX5 | purinergic receptor P2X 5 |
| CC2D2A | coiled-coil and C2 domain containing 2A | P3H3 | prolyl 3-hydroxylase 3 |
| CCDC142 | coiled-coil domain containing 142 | P3H4 | prolyl 3-hydroxylase family member 4 |
| CCDC88C | coiled-coil domain containing 88C | PCDH7 | protocadherin 7 |
| CCL13 | C-C motif chemokine ligand 13 | PCNX2 | pecanex homolog 2 |
| CCL18 | C-C motif chemokine ligand 18 | PDLIM4 | PDZ and LIM domain 4 |

| | | | |
|---|---|---|---|
| CCND2 | cyclin D2 | PDLIM7 | PDZ and LIM domain 7 |
| CD248 | CD248 molecule | PGM5 | phosphoglucomutase 5 |
| CDH3 | cadherin 3 | PIK3R2 | phosphoinositide-3-kinase regulatory subunit 2 |
| CDKN2C | cyclin dependent kinase inhibitor 2C | PKIB | protein kinase (cAMP-dependent, catalytic) inhibitor beta |
| CEP250 | centrosomal protein 250 | PLA2G15 | phospholipase A2 group XV |
| CFH | complement factor H | PLEKHA4 | pleckstrin homology domain containing A4 |
| CHST3 | carbohydrate sulfotransferase 3 | POSTN | Periostin |
| CLDN15 | claudin 15 | PPP1R12B | protein phosphatase 1 regulatory subunit 12B |
| CLIP2 | CAP-Gly domain containing linker protein 2 | PRAF2 | PRA1 domain family member 2 |
| CLMN | calmin | PRUNE2 | prune homolog 2 |
| COL14A1 | collagen type XIV alpha 1 chain | PSD3 | pleckstrin and Sec7 domain containing 3 |
| COL15A1 | collagen type XV alpha 1 chain | PTGFRN | prostaglandin F2 receptor inhibitor |
| COL16A1 | collagen type XVI alpha 1 chain | R3HDM1 | R3H domain containing 1 |
| COL18A1 | collagen type XVIII alpha 1 chain | RAB3GAP1 | RAB3 GTPase activating protein catalytic subunit 1 |
| COL5A2 | collagen type V alpha 2 chain | RAMP1 | receptor activity modifying protein 1 |
| COL6A1 | collagen type VI alpha 1 chain | RBBP4 | RB binding protein 4, chromatin remodeling factor |
| COL6A2 | collagen type VI alpha 2 chain | RHOD | ras homolog family member D |
| COL6A3 | collagen type VI alpha 3 chain | RNFT2 | ring finger protein, transmembrane 2 |
| COL7A1 | collagen type VII alpha 1 chain | ROR2 | receptor tyrosine kinase like orphan receptor 2 |
| COMP | cartilage oligomeric matrix protein | S100A2 | S100 calcium binding protein A2 |
| CP | ceruloplasmin | SCARA3 | scavenger receptor class A member 3 |
| CPXM1 | carboxypeptidase X, M14 family member 1 | SCG5 | secretogranin V |
| CXCL14 | C-X-C motif chemokine ligand 14 | SEMA3C | semaphorin 3C |
| DCLK1 | doublecortin like kinase 1 | SERINC2 | serine incorporator 2 |
| DDO | D-aspartate oxidase | SERPINB5 | serpin family B member 5 |
| DIO2 | deiodinase, iodothyronine type II | SEZ6L2 | seizure related 6 homolog like 2 |
| DOCK3 | dedicator of cytokinesis 3 | SFI1 | SFI1 centrin binding protein |
| DOK5 | docking protein 5 | SFXN4 | sideroflexin 4 |
| ECM1 | extracellular matrix protein 1 | SH3GL1 | SH3 domain containing GRB2 like 1, endophilin A2 |
| EFNB3 | ephrin B3 | SHC3 | SHC adaptor protein 3 |
| EGFL6 | EGF like domain multiple 6 | SLC1A4 | solute carrier family 1 member 4 |
| EML6 | echinoderm microtubule associated protein like 6 | SLC22A23 | solute carrier family 22 member 23 |
| EPHB2 | EPH receptor B2 | SLC29A3 | solute carrier family 29 member 3 |
| EYA2 | EYA transcriptional coactivator and phosphatase 2 | SLC4A4 | solute carrier family 4 member 4 |
| FBLN1 | fibulin 1 | SLC6A8 | solute carrier family 6 member 8 |
| FBLN2 | fibulin 2 | SNCAIP | synuclein alpha interacting protein |
| FBXW9 | F-box and WD repeat domain containing 9 | SPP1 | secreted phosphoprotein 1 |
| FHOD3 | formin homology 2 domain containing 3 | SRGAP3 | SLIT-ROBO Rho GTPase activating protein 3 |

| | | | |
|---|---|---|---|
| FLRT2 | fibronectin leucine rich transmembrane protein 2 | STK36 | serine/threonine kinase 36 |
| FMO1 | flavin containing monooxygenase 1 | STK38 | serine/threonine kinase 38 |
| FUT6 | fucosyltransferase 6 | STMN3 | stathmin 3 |
| GABBR2 | gamma-aminobutyric acid type B receptor subunit 2 | SYNDIG1 | synapse differentiation inducing 1 |
| GALNT5 | polypeptide N-acetylgalactosaminyltransferase 5 | SYNJ2 | synaptojanin 2 |
| GDF11 | growth differentiation factor 11 | SYNPO2 | synaptopodin 2 |
| GLT8D2 | glycosyltransferase 8 domain containing 2 | SYNRG | synergin gamma |
| GPC1 | glypican 1 | SYT8 | synaptotagmin 8 |
| GPR87 | G protein-coupled receptor 87 | SYTL2 | synaptotagmin like 2 |
| GPX7 | glutathione peroxidase 7 | TAB1 | TGF-beta activated kinase 1 binding protein 1 |
| GUCY1A3 | guanylate cyclase 1 soluble subunit alpha | TENM4 | teneurin transmembrane protein 4 |
| H2AFY2 | H2A histone family member Y2 | TFF3 | trefoil factor 3 |
| HEPH | hephaestin | TGFB3 | transforming growth factor beta 3 |
| HOMER3 | homer scaffolding protein 3 | TGFBI | transforming growth factor beta induced |
| | | TGFBR2 | transforming growth factor beta receptor 2 |
| HS3ST1 | heparan sulfate-glucosamine 3-sulfotransferase 1 | THBS2 | thrombospondin 2 |
| HSD3B7 | hydroxy-delta-5-steroid dehydrogenase, 3 beta- and steroid delta-isomerase 7 | TLDC1 | TBC/LysM-associated domain containing 1 |
| HSPA4L | heat shock protein family A member 4 like | TM7SF3 | transmembrane 7 superfamily member 3 |
| IGF1 | insulin like growth factor 1 | TMED10 | transmembrane p24 trafficking protein 10 |
| IGFBP2 | insulin like growth factor binding protein 2 | TMEM158 | transmembrane protein 158 (gene/pseudogene) |
| IL13RA2 | interleukin 13 receptor subunit alpha 2 | TMEM40 | transmembrane protein 40 |
| ILF3 | interleukin enhancer binding factor 3 | TNPO1 | transportin 1 |
| INPP5D | inositol polyphosphate-5-phosphatase D | TP73 | tumor protein p73 |
| ITGA7 | integrin subunit alpha 7 | TRA2A | transformer 2 alpha homolog |
| ITGB4 | integrin subunit beta 4 | TRO | trophinin |
| KCNMA1 | potassium calcium-activated channel subfamily M alpha 1 | TRPM4 | transient receptor potential cation channel subfamily M member 4 |
| KCNMB1 | potassium calcium-activated channel subfamily M regulatory beta subunit 1 | UGT1A9 | UDP glucuronosyltransferase family 1 member A9 |
| KCNN3 | potassium calcium-activated channel subfamily N member 3 | USP35 | ubiquitin specific peptidase 35 |
| KCNN4 | potassium calcium-activated channel subfamily N member 4 | VILL | villin like |
| KIAA0100 | KIAA0100 | VWA1 | von Willebrand factor A domain containing 1 |
| KIAA0101 | KIAA0101 (PCLAF) | WRAP53 | WD repeat containing antisense to TP53 |
| LDOC1 | leucine zipper down-regulated in cancer 1 | XPR1 | xenotropic and polytropic retrovirus receptor 1 |
| LGALS7 | galectin 7 | ZDHHC13 | zinc finger DHHC-type containing 13 |
| LOC730101 | uncharacterized LOC730101 | ZMIZ1 | zinc finger MIZ-type containing 1 |
| LOXL1 | lysyl oxidase like 1 | ZNF207 | zinc finger protein 207 |
| LOXL2 | lysyl oxidase like 2 | ZNF71 | zinc finger protein 71 |

**Supplemental Table S2: Attenuation of the TGF-β response following PRRX1 knock-down.**

The table contains the best 597 genes significantly modulated by TGF-β in NHLF treated with siControl#1 and #2 (log2 fold change >4 ; adj. P Val<0.05). The table indicates the Fold Change (FC) for each of the genes: i) column 1: TGF-β1" stimulation in presence of siControl#1 ; column 2: TGFβ1 fold stimulation in presence of siRNA#1 ; column 4: TGFβ1 fold stimulation in presence of siControl#2 ; column 5: TGFβ1 fold stimulation in presence of siRNA#2". Columns 5 and 6 give the percentage of residual modulation by TGF-β for each PRRX1 siRNA. Modulations are shown in progressively brighter shades of blue (attenuation) and orange (over-activation). The mean residual fold change following PRRX1 KD is 56.6 % (siRNA#1) and 80.0 % (siRNA#2).

| GeneName | TGF-β1 fold modulation | | Residual FC (%) siRNA#1 | TGF-β1 fold modulation | | Residual FC (%) siRNA#2 |
|---|---|---|---|---|---|---|
| | siControl#1 | PRRX1 siRNA#1 | | siControl#2 | PRRX1 siRNA#2 | |
| COMP | 181.83 | 76.84 | 42.26 | 226.75 | 117.46 | 51.80 |
| NPPB | 92.03 | 10.45 | 11.35 | 54.08 | 27.80 | 51.41 |
| NOX4 | 87.32 | 36.32 | 41.59 | 66.29 | 47.43 | 71.55 |
| ENST00000573315 | 66.91 | 8.42 | 12.58 | 59.96 | 13.83 | 23.07 |
| NPPB | 48.50 | 5.59 | 11.53 | 32.47 | 18.37 | 56.58 |
| IGF1 | 41.21 | 12.97 | 31.47 | 39.15 | 14.68 | 37.50 |
| ELN | 39.80 | 10.85 | 27.26 | 65.16 | 57.14 | 87.69 |
| IGFBP3 | 36.44 | 6.17 | 16.93 | 16.64 | 11.73 | 70.50 |
| ITGA11 | 33.73 | 13.98 | 41.46 | 41.36 | 29.23 | 70.69 |
| TSPAN2 | 29.27 | 6.97 | 23.82 | 25.73 | 21.02 | 81.70 |
| FZD8 | 28.30 | 20.28 | 71.67 | 20.62 | 16.79 | 81.43 |
| SCRG1 | 28.06 | 5.04 | 17.95 | 26.01 | 11.43 | 43.93 |
| GDF6 | 24.92 | 11.94 | 47.91 | 33.72 | 15.22 | 45.12 |
| SLCO2A1 | 24.15 | 34.04 | 140.95 | 40.86 | 16.00 | 39.17 |
| ANKRD1 | 22.65 | 24.48 | 108.08 | 24.10 | 20.21 | 83.84 |
| AMIGO2 | 22.18 | 10.21 | 46.01 | 20.72 | 24.90 | 120.21 |
| SCX | 21.98 | 11.28 | 51.32 | 19.76 | 13.65 | 69.07 |
| MDFI | 21.89 | 4.26 | 19.47 | 18.44 | 14.31 | 77.61 |
| HAPLN1 | 21.83 | 6.13 | 28.08 | 17.97 | 5.47 | 30.44 |
| IL11 | 21.82 | 17.89 | 82.03 | 28.36 | 43.26 | 152.53 |
| SERTAD4 | 19.91 | 10.82 | 54.36 | 36.86 | 17.42 | 47.26 |
| HBEGF | 19.20 | 18.65 | 97.17 | 20.03 | 28.56 | 142.60 |
| PMEPA1 | 18.83 | 15.98 | 84.87 | 18.07 | 14.78 | 81.77 |
| KANK4 | 18.46 | 1.85 | 10.01 | 14.64 | 8.18 | 55.87 |
| CTPS1 | 18.08 | 11.99 | 66.32 | 15.62 | 16.23 | 103.95 |
| ELN | 18.05 | 8.40 | 46.54 | 23.79 | 15.05 | 63.28 |
| PRG4 | 17.89 | 4.65 | 26.02 | 18.10 | 7.58 | 41.90 |
| SERTAD4-AS1 | 17.58 | 13.00 | 73.91 | 32.23 | 24.67 | 76.54 |
| NKX3-1 | 16.94 | 3.47 | 20.51 | 10.40 | 8.59 | 82.63 |
| MEGF6 | 16.78 | 12.62 | 75.24 | 26.04 | 9.22 | 35.40 |
| KANK4 | 16.59 | 1.89 | 11.41 | 12.33 | 6.99 | 56.69 |
| FOXS1 | 16.51 | 7.88 | 47.75 | 7.81 | 9.68 | 124.01 |
| CNN1 | 15.29 | 5.32 | 34.81 | 18.08 | 9.98 | 55.19 |
| LINC01711 | 14.97 | 5.30 | 35.41 | 11.03 | 8.17 | 74.09 |
| LINC02593 | 14.45 | 10.38 | 71.82 | 7.71 | 12.13 | 157.34 |
| CTPS1 | 13.65 | 9.96 | 72.93 | 12.21 | 13.11 | 107.33 |
| EGR2 | 13.25 | 14.07 | 106.15 | 12.00 | 20.39 | 169.94 |
| TNFAIP6 | 13.23 | 3.73 | 28.15 | 8.36 | 11.21 | 134.14 |
| DSP | 12.36 | 6.61 | 53.43 | 13.41 | 9.37 | 69.89 |

| | | | | | | |
|---|---|---|---|---|---|---|
| PMEPA1 | 12.35 | 12.27 | 99.42 | 13.56 | 13.19 | 97.28 |
| TPM1 | 12.12 | 9.58 | 79.00 | 15.34 | 15.06 | 98.18 |
| IGF1 | 11.88 | 4.53 | 38.12 | 10.33 | 6.47 | 62.68 |
| SERPINE1 | 11.84 | 15.32 | 129.40 | 27.83 | 32.38 | 116.34 |
| FSTL3 | 11.73 | 7.31 | 62.28 | 9.21 | 7.15 | 77.55 |
| BMP6 | 11.69 | 7.11 | 60.81 | 14.99 | 6.88 | 45.88 |
| LOC105377123 | 11.50 | 3.62 | 31.46 | 7.09 | 4.72 | 66.62 |
| COL4A1 | 11.48 | 6.98 | 60.80 | 12.26 | 10.42 | 84.95 |
| LYPD1 | 11.03 | 5.40 | 48.98 | 18.58 | 19.39 | 104.35 |
| TPM1 | 10.94 | 8.90 | 81.31 | 12.16 | 8.30 | 68.28 |
| XRCC4 | 10.53 | 2.90 | 27.53 | 6.90 | 7.18 | 104.08 |
| MFAP5 | 10.50 | 4.18 | 39.77 | 12.88 | 2.57 | 19.97 |
| lnc-RP11-597K23.2.1-2 | 10.49 | 3.40 | 32.40 | 8.06 | 5.14 | 63.80 |
| BHLHE40 | 10.38 | 6.97 | 67.22 | 8.17 | 9.36 | 114.62 |
| LOC105369340 | 10.16 | 3.87 | 38.07 | 5.31 | 2.80 | 52.68 |
| SERPINE2 | 10.06 | 8.52 | 84.72 | 12.50 | 19.06 | 152.44 |
| APCDD1L | 9.97 | 6.06 | 60.82 | 11.82 | 7.74 | 65.49 |
| PTGS1 | 9.96 | 6.40 | 64.27 | 8.43 | 5.19 | 61.49 |
| FGD5-AS1 | 9.73 | 4.18 | 42.98 | 9.86 | 6.94 | 70.39 |
| TPM1 | 9.68 | 7.58 | 78.35 | 13.55 | 8.67 | 63.98 |
| LOC105369205 | 9.47 | 7.01 | 74.07 | 16.54 | 8.60 | 51.97 |
| NUAK1 | 9.28 | 5.41 | 58.30 | 11.50 | 7.23 | 62.91 |
| SYT12 | 9.03 | 12.75 | 141.21 | 10.69 | 13.11 | 122.64 |
| COL10A1 | 8.93 | 3.47 | 38.83 | 11.21 | 9.34 | 83.32 |
| NOX4 | 8.76 | 5.78 | 66.00 | 8.13 | 7.67 | 94.27 |
| NPR3 | 8.66 | 3.10 | 35.74 | 5.36 | 4.96 | 92.44 |
| MXRA5 | 8.66 | 7.98 | 92.12 | 5.67 | 10.20 | 180.08 |
| ENC1 | 8.66 | 3.16 | 36.52 | 6.42 | 8.73 | 135.98 |
| IGFBP7 | 8.60 | 2.96 | 34.43 | 9.07 | 4.97 | 54.82 |
| WRB-SH3BGR | 8.48 | 2.45 | 28.90 | 6.78 | 3.19 | 47.02 |
| ADAM12 | 8.42 | 2.55 | 30.24 | 4.39 | 5.83 | 132.76 |
| IGFBP7 | 8.42 | 3.31 | 39.35 | 8.91 | 4.10 | 45.99 |
| LDLRAD4 | 8.40 | 2.62 | 31.24 | 5.18 | 3.48 | 67.10 |
| LOC107985502 | 8.39 | 8.08 | 96.36 | 6.30 | 11.51 | 182.64 |
| NPR3 | 8.36 | 3.51 | 42.02 | 4.34 | 3.95 | 90.97 |
| ADAM12 | 8.34 | 1.95 | 23.42 | 4.52 | 5.96 | 131.70 |
| GXYLT2 | 8.28 | 2.51 | 30.33 | 6.62 | 5.40 | 81.51 |
| PAWR | 8.23 | 4.36 | 53.04 | 9.09 | 7.74 | 85.11 |
| SEMA7A | 8.18 | 6.23 | 76.15 | 12.96 | 14.15 | 109.18 |
| ACTBL2 | 8.14 | 5.47 | 67.25 | 11.75 | 6.54 | 55.68 |
| CHAC1 | 8.14 | 5.78 | 71.00 | 13.74 | 12.93 | 94.11 |
| NALCN | 8.11 | 4.56 | 56.23 | 5.49 | 6.46 | 117.56 |
| ZNF365 | 8.06 | 5.83 | 72.27 | 7.41 | 8.15 | 109.97 |
| DYSF | 8.03 | 3.12 | 38.83 | 4.32 | 2.69 | 62.30 |
| ASNSP1 | 7.92 | 3.44 | 43.39 | 9.46 | 7.35 | 77.76 |
| XYLT1 | 7.84 | 5.52 | 70.37 | 5.89 | 7.03 | 119.35 |
| LYPD1 | 7.80 | 3.92 | 50.24 | 11.91 | 11.42 | 95.83 |
| ASNSP1 | 7.60 | 3.34 | 43.89 | 9.74 | 8.10 | 83.12 |
| PLCB4 | 7.54 | 2.79 | 37.07 | 6.06 | 5.00 | 82.46 |
| MFAP3L | 7.49 | 3.90 | 52.12 | 8.99 | 4.06 | 45.13 |
| HHIP | 7.36 | 4.23 | 57.44 | 8.46 | 3.45 | 40.76 |
| PLXDC2 | 7.34 | 6.83 | 93.06 | 8.37 | 7.46 | 89.12 |
| PI16 | 7.32 | 2.79 | 38.12 | 7.67 | 5.95 | 77.59 |
| DSP | 7.30 | 3.23 | 44.33 | 6.88 | 5.57 | 80.90 |
| MYH11 | 7.23 | 5.28 | 73.02 | 11.30 | 3.86 | 34.20 |
| MYH11 | 7.21 | 4.85 | 67.29 | 11.48 | 3.84 | 33.48 |
| PLPP4 | 7.18 | 4.30 | 59.92 | 7.77 | 4.28 | 55.11 |
| MTHFD2 | 7.16 | 2.51 | 35.14 | 7.77 | 5.38 | 69.21 |

| | | | | | | |
|---|---|---|---|---|---|---|
| A_33_P3368900 | 7.08 | 1.88 | 26.57 | 6.03 | 3.88 | 64.32 |
| PSAT1 | 7.07 | 3.65 | 51.59 | 8.30 | 6.31 | 76.04 |
| AMTN | 7.07 | 1.77 | 25.11 | 6.38 | 3.31 | 51.88 |
| CDH2 | 7.02 | 2.75 | 39.15 | 3.70 | 3.39 | 91.56 |
| ENST00000511103 | 6.94 | 3.88 | 55.82 | 9.41 | 7.92 | 84.15 |
| SPDL1 | 6.94 | 2.86 | 41.27 | 5.23 | 7.67 | 146.86 |
| COL5A1 | 6.91 | 4.13 | 59.85 | 5.59 | 7.97 | 142.45 |
| LOC105369205 | 6.86 | 5.11 | 74.39 | 9.22 | 6.95 | 75.44 |
| HSD17B6 | 6.79 | 3.00 | 44.16 | 8.49 | 7.12 | 83.93 |
| PTHLH | 6.75 | 3.96 | 58.63 | 5.42 | 10.19 | 188.11 |
| ASPN | 6.71 | 2.10 | 31.32 | 8.38 | 4.99 | 59.50 |
| ASNS | 6.70 | 2.92 | 43.68 | 9.01 | 6.75 | 74.98 |
| SYNDIG1 | 6.67 | 3.33 | 50.00 | 4.01 | 4.76 | 118.65 |
| ELFN2 | 6.67 | 4.39 | 65.85 | 7.14 | 3.95 | 55.35 |
| PLOD2 | 6.66 | 3.54 | 53.23 | 4.01 | 7.92 | 197.64 |
| EFR3B | 6.66 | 5.28 | 79.29 | 4.76 | 6.41 | 134.69 |
| lnc-CABLES1-1 | 6.63 | 0.99 | 14.92 | 5.35 | 1.60 | 29.85 |
| NCF2 | 6.58 | 4.05 | 61.56 | 6.01 | 5.05 | 84.05 |
| GFRA1 | 6.53 | 3.41 | 52.27 | 6.86 | 3.90 | 56.88 |
| CEMIP2 | 6.49 | 2.41 | 37.16 | 5.05 | 6.74 | 133.53 |
| GADD45B | 6.49 | 12.37 | 190.64 | 9.88 | 12.23 | 123.75 |
| TES | 6.46 | 3.89 | 60.14 | 6.14 | 3.21 | 52.36 |
| MCAM | 6.45 | 2.78 | 43.10 | 6.85 | 3.74 | 54.57 |
| CRLF1 | 6.40 | 5.43 | 84.78 | 7.24 | 5.26 | 72.65 |
| ACTA2 | 6.38 | 7.65 | 119.88 | 11.69 | 7.65 | 65.43 |
| ENST00000518968 | 6.38 | 3.27 | 51.19 | 8.51 | 7.42 | 87.20 |
| NXPH4 | 6.30 | 6.13 | 97.34 | 6.74 | 6.20 | 92.01 |
| ACTG2 | 6.29 | 7.34 | 116.78 | 13.67 | 9.72 | 71.12 |
| ATP10A | 6.27 | 7.38 | 117.64 | 7.69 | 8.97 | 116.76 |
| A_33_P3255824 | 6.22 | 5.35 | 85.95 | 4.85 | 6.86 | 141.52 |
| CHAC1 | 6.21 | 5.52 | 88.88 | 9.54 | 8.96 | 93.95 |
| COL4A2 | 6.18 | 3.92 | 63.38 | 4.77 | 3.81 | 79.96 |
| ELN-AS1 | 6.11 | 3.56 | 58.16 | 7.39 | 4.67 | 63.25 |
| LIF | 5.93 | 6.14 | 103.54 | 7.15 | 8.07 | 112.93 |
| ADM2 | 5.89 | 4.33 | 73.51 | 8.06 | 5.67 | 70.38 |
| LOX | 5.85 | 3.50 | 59.80 | 6.42 | 5.03 | 78.34 |
| LINC01614 | 5.85 | 1.39 | 23.82 | 2.05 | 4.32 | 210.69 |
| UCK2 | 5.84 | 4.63 | 79.21 | 5.76 | 6.88 | 119.51 |
| ISLR2 | 5.84 | 3.08 | 52.68 | 3.89 | 3.13 | 80.48 |
| TNFSF4 | 5.80 | 4.43 | 76.38 | 10.15 | 3.97 | 39.10 |
| ACKR3 | 5.74 | 2.48 | 43.13 | 3.64 | 2.78 | 76.49 |
| CTH | 5.72 | 2.88 | 50.35 | 5.52 | 4.30 | 77.76 |
| SLC7A1 | 5.71 | 3.54 | 62.05 | 7.88 | 7.75 | 98.30 |
| CDH2 | 5.64 | 2.12 | 37.62 | 2.98 | 2.72 | 91.12 |
| DUSP26 | 5.63 | 1.38 | 24.62 | 5.58 | 2.22 | 39.76 |
| OXTR | 5.62 | 3.02 | 53.67 | 4.13 | 5.66 | 137.21 |
| PLCB4 | 5.61 | 2.56 | 45.52 | 5.17 | 3.60 | 69.69 |
| AMZ1 | 5.56 | 7.71 | 138.73 | 4.63 | 4.30 | 92.68 |
| EDN1 | 5.53 | 6.36 | 115.01 | 19.43 | 15.04 | 77.40 |
| EDIL3 | 5.52 | 2.02 | 36.55 | 4.41 | 3.96 | 89.86 |
| LF212376 | 5.51 | 2.73 | 49.50 | 5.62 | 3.31 | 58.98 |
| KCNN4 | 5.43 | 5.62 | 103.59 | 7.22 | 3.19 | 44.15 |
| HHIP | 5.39 | 2.87 | 53.15 | 5.69 | 2.98 | 52.40 |
| HSPA13 | 5.38 | 2.59 | 48.12 | 4.81 | 4.97 | 103.30 |
| GALNT18 | 5.38 | 3.70 | 68.81 | 7.15 | 4.21 | 58.84 |
| SPDL1 | 5.37 | 2.61 | 48.65 | 5.07 | 6.68 | 131.62 |
| FGF1 | 5.35 | 4.68 | 87.46 | 7.23 | 5.49 | 75.88 |
| LOC101928076 | 5.35 | 3.72 | 69.51 | 4.43 | 3.36 | 75.76 |

| | | | | | | |
|---|---|---|---|---|---|---|
| TNS1 | 5.34 | 4.79 | 89.79 | 4.89 | 3.51 | 71.74 |
| UNC5B | 5.32 | 2.45 | 46.09 | 5.57 | 4.49 | 80.68 |
| ASPN | 5.32 | 1.72 | 32.28 | 5.60 | 4.44 | 79.35 |
| SPOCK1 | 5.30 | 3.95 | 74.52 | 4.92 | 3.92 | 79.65 |
| PRPS1L1 | 5.29 | 3.66 | 69.24 | 4.40 | 4.07 | 92.50 |
| SGCG | 5.27 | 2.71 | 51.39 | 9.71 | 3.18 | 32.79 |
| ADAMTS6 | 5.27 | 3.40 | 64.48 | 4.72 | 4.39 | 93.02 |
| MSMO1 | 5.23 | 2.60 | 49.77 | 4.06 | 4.05 | 99.80 |
| RFLNB | 5.22 | 3.44 | 65.90 | 4.16 | 4.25 | 102.11 |
| COL5A1 | 5.16 | 3.15 | 61.12 | 5.11 | 5.26 | 102.99 |
| PLN | 5.15 | 3.28 | 63.64 | 5.62 | 4.18 | 74.28 |
| LIMS2 | 5.14 | 3.37 | 65.71 | 7.50 | 4.15 | 55.34 |
| LOC105379057 | 5.08 | 3.36 | 66.03 | 5.98 | 4.11 | 68.80 |
| PCED1B | 5.07 | 4.06 | 80.09 | 5.43 | 4.90 | 90.27 |
| LOC105369340 | 5.06 | 3.77 | 74.49 | 4.95 | 2.41 | 48.63 |
| CCN2 | 5.06 | 4.52 | 89.37 | 9.73 | 8.96 | 92.05 |
| SIK1 | 5.04 | 4.34 | 86.06 | 4.06 | 7.46 | 183.85 |
| KLHDC7B | 5.02 | 3.79 | 75.43 | 4.44 | 4.46 | 100.34 |
| OSBPL10 | 5.00 | 3.82 | 76.43 | 6.27 | 4.93 | 78.57 |
| LINC01605 | 4.99 | 4.28 | 85.81 | 7.13 | 7.83 | 109.85 |
| ANXA8L1 | 4.97 | 2.73 | 55.03 | 3.09 | 1.82 | 59.01 |
| LOC105379010 | 4.97 | 2.01 | 40.42 | 3.62 | 2.29 | 63.39 |
| ALPK2 | 4.96 | 3.35 | 67.52 | 5.19 | 5.51 | 106.09 |
| MRAS | 4.95 | 5.16 | 104.25 | 4.63 | 4.66 | 100.69 |
| MYOSLID | 4.94 | 6.93 | 140.21 | 5.67 | 6.12 | 107.88 |
| KRT80 | 4.94 | 3.80 | 76.91 | 4.36 | 5.55 | 127.43 |
| ATP1B1 | 4.94 | 1.59 | 32.16 | 3.14 | 2.53 | 80.63 |
| SKIL | 4.92 | 2.57 | 52.14 | 3.51 | 5.40 | 153.77 |
| GALNT10 | 4.91 | 3.37 | 68.73 | 3.57 | 2.92 | 81.97 |
| LINC01013 | 4.89 | 3.34 | 68.22 | 5.64 | 3.21 | 56.93 |
| PROC | 4.89 | 8.23 | 168.46 | 2.95 | 7.15 | 242.19 |
| LOC105370706 | 4.87 | 2.67 | 54.87 | 3.56 | 1.93 | 54.27 |
| TUFT1 | 4.84 | 3.68 | 76.01 | 5.98 | 7.04 | 117.62 |
| PDLIM5 | 4.83 | 3.04 | 62.82 | 4.20 | 4.04 | 96.33 |
| TAGLN | 4.83 | 6.87 | 142.16 | 13.10 | 8.96 | 68.41 |
| ACTG2 | 4.83 | 3.20 | 66.32 | 4.28 | 7.49 | 175.08 |
| LINC01638 | 4.80 | 3.34 | 69.54 | 3.12 | 2.71 | 86.89 |
| PRPS1 | 4.80 | 3.40 | 70.94 | 4.38 | 3.76 | 85.65 |
| SMCO4 | 4.78 | 3.68 | 76.93 | 6.02 | 5.35 | 88.84 |
| KCNG1 | 4.77 | 4.29 | 89.87 | 7.12 | 6.62 | 93.05 |
| RUNX1 | 4.77 | 3.08 | 64.42 | 4.24 | 5.07 | 119.52 |
| POSTN | 4.77 | 2.39 | 50.13 | 5.68 | 9.83 | 172.89 |
| FLJ43315 | 4.77 | 2.48 | 51.94 | 5.31 | 4.51 | 84.95 |
| LINC00842 | 4.76 | 3.28 | 68.78 | 6.24 | 3.71 | 59.40 |
| ALDH1L2 | 4.75 | 1.89 | 39.75 | 4.37 | 3.74 | 85.52 |
| CALD1 | 4.75 | 3.32 | 69.85 | 4.12 | 4.34 | 105.33 |
| SUSD2 | 4.75 | 3.02 | 63.61 | 10.76 | 4.51 | 41.95 |
| DYNLT3 | 4.72 | 2.56 | 54.32 | 3.12 | 3.96 | 127.17 |
| KRT18 | 4.72 | 3.05 | 64.65 | 5.12 | 4.33 | 84.62 |
| UAP1 | 4.71 | 3.21 | 68.12 | 5.44 | 5.45 | 100.19 |
| DMD | 4.67 | 2.57 | 54.94 | 4.71 | 5.21 | 110.62 |
| ALDH1B1 | 4.67 | 3.61 | 77.28 | 4.72 | 3.66 | 77.53 |
| PRPS1 | 4.66 | 2.39 | 51.33 | 4.56 | 4.23 | 92.78 |
| SLC7A5 | 4.63 | 3.53 | 76.08 | 3.57 | 5.64 | 158.08 |
| ITGBL1 | 4.63 | 3.50 | 75.70 | 4.59 | 3.91 | 85.25 |
| CILP | 4.61 | 1.69 | 36.69 | 5.25 | 2.92 | 55.66 |
| ADAM19 | 4.60 | 3.07 | 66.80 | 5.21 | 5.64 | 108.37 |
| EIF4EBP1 | 4.60 | 3.47 | 75.45 | 7.33 | 4.66 | 63.58 |

| | | | | | | |
|---|---|---|---|---|---|---|
| ADAMTS6 | 4.60 | 2.81 | 61.15 | 4.46 | 3.58 | 80.25 |
| IGFBP7 | 4.57 | 2.30 | 50.25 | 4.57 | 3.14 | 68.63 |
| MYOSLID | 4.57 | 6.78 | 148.27 | 5.60 | 6.00 | 107.23 |
| LINC01969 | 4.57 | 3.72 | 81.50 | 3.27 | 3.61 | 110.26 |
| SLC17A9 | 4.56 | 4.62 | 101.40 | 3.95 | 4.55 | 115.25 |
| TRIB3 | 4.55 | 2.41 | 52.92 | 5.68 | 5.35 | 94.12 |
| SEC11C | 4.53 | 2.64 | 58.35 | 3.76 | 3.08 | 81.85 |
| SRRM3 | 4.53 | 4.75 | 104.86 | 6.51 | 3.72 | 57.06 |
| DIAPH3 | 4.53 | 1.70 | 37.63 | 4.05 | 2.98 | 73.60 |
| QPCT | 4.51 | 1.54 | 34.20 | 3.19 | 2.90 | 90.90 |
| GLS | 4.50 | 1.56 | 34.64 | 2.22 | 2.04 | 92.15 |
| WNT5B | 4.49 | 3.84 | 85.53 | 4.78 | 4.58 | 95.72 |
| MATN3 | 4.48 | 2.26 | 50.46 | 4.79 | 4.11 | 85.80 |
| LINC02407 | 4.47 | 5.85 | 130.89 | 6.60 | 7.00 | 106.09 |
| C1QTNF3 | 4.47 | 1.28 | 28.53 | 2.06 | 1.82 | 88.36 |
| SNAR-F | 4.46 | 3.18 | 71.33 | 4.82 | 3.68 | 76.34 |
| CHST6 | 4.46 | 3.47 | 77.79 | 3.27 | 3.94 | 120.30 |
| TAGLN | 4.46 | 6.83 | 153.28 | 13.19 | 8.36 | 63.41 |
| ENST00000439132 | 4.45 | 2.49 | 55.99 | 5.46 | 5.52 | 101.07 |
| TXNDC5 | 4.42 | 1.80 | 40.80 | 3.65 | 4.72 | 129.09 |
| ITGBL1 | 4.42 | 3.69 | 83.68 | 5.17 | 4.12 | 79.70 |
| MMP24 | 4.40 | 2.72 | 61.86 | 4.81 | 2.71 | 56.29 |
| THUMPD2 | 4.39 | 3.72 | 84.71 | 3.13 | 4.06 | 129.84 |
| CAMK1D | 4.39 | 3.25 | 74.12 | 5.75 | 4.37 | 75.91 |
| HES6 | 4.38 | 5.32 | 121.36 | 6.44 | 4.53 | 70.38 |
| MLLT11 | 4.38 | 3.73 | 85.14 | 5.18 | 3.30 | 63.70 |
| THBS2 | 4.38 | 4.40 | 100.33 | 4.48 | 4.17 | 93.03 |
| ADAM12 | 4.38 | 2.53 | 57.71 | 3.05 | 4.98 | 163.09 |
| GLS | 4.37 | 2.14 | 48.94 | 2.61 | 2.00 | 76.75 |
| THC2642375 | 4.37 | 3.73 | 85.52 | 3.34 | 3.05 | 91.28 |
| SH3PXD2A | 4.37 | 3.58 | 81.97 | 3.94 | 4.04 | 102.74 |
| SH3PXD2A | 4.36 | 2.75 | 63.20 | 3.54 | 3.26 | 92.08 |
| TMC7 | 4.35 | 3.01 | 69.23 | 2.89 | 3.02 | 104.54 |
| RAI14 | 4.33 | 3.20 | 73.92 | 4.28 | 6.15 | 143.74 |
| GLS | 4.31 | 2.23 | 51.89 | 2.99 | 5.21 | 174.06 |
| KRT18 | 4.30 | 2.96 | 68.88 | 5.67 | 4.47 | 78.83 |
| DACT1 | 4.29 | 2.29 | 53.52 | 3.28 | 4.19 | 127.82 |
| SNAR-G2 | 4.28 | 2.77 | 64.69 | 4.25 | 3.30 | 77.70 |
| CLIC4 | 4.28 | 2.87 | 67.00 | 4.53 | 3.63 | 80.06 |
| KRT17 | 4.28 | 2.85 | 66.59 | 5.15 | 2.96 | 57.35 |
| INHBA | 4.27 | 3.16 | 74.01 | 3.27 | 4.63 | 141.40 |
| RTKN2 | 4.27 | 2.69 | 63.03 | 5.48 | 5.08 | 92.79 |
| SAMD11 | 4.26 | 3.80 | 89.21 | 3.42 | 4.12 | 120.31 |
| ABCA3 | 4.25 | 2.53 | 59.60 | 4.48 | 3.52 | 78.49 |
| GLIPR2 | 4.23 | 2.74 | 64.65 | 4.78 | 3.10 | 64.91 |
| PTPRN | 4.23 | 2.28 | 53.78 | 5.33 | 2.32 | 43.48 |
| PDLIM3 | 4.22 | 2.29 | 54.28 | 5.09 | 4.58 | 90.14 |
| CDH6 | 4.22 | 3.46 | 81.96 | 3.24 | 6.39 | 197.23 |
| TSPAN13 | 4.21 | 1.70 | 40.41 | 4.72 | 3.16 | 66.88 |
| DYNLT3 | 4.20 | 2.21 | 52.62 | 2.81 | 3.48 | 123.70 |
| PXDC1 | 4.20 | 4.48 | 106.58 | 4.75 | 4.90 | 103.06 |
| RBP1 | 4.19 | 2.54 | 60.65 | 5.18 | 2.25 | 43.41 |
| SNAR-D | 4.17 | 2.33 | 55.73 | 3.84 | 2.87 | 74.65 |
| LINC01583 | 4.16 | 2.29 | 55.10 | 3.36 | 1.92 | 57.25 |
| SNAR-H | 4.15 | 2.53 | 60.95 | 3.89 | 3.17 | 81.38 |
| lnc-CALHM3-2 | 4.15 | 4.01 | 96.51 | 2.98 | 2.65 | 89.06 |
| LINC00592 | 4.15 | 3.26 | 78.50 | 4.25 | 4.28 | 100.64 |
| PCK2 | 4.14 | 2.84 | 68.72 | 5.77 | 3.00 | 51.98 |

| | | | | | | |
|---|---|---|---|---|---|---|
| SORCS2 | 4.14 | 3.81 | 91.98 | 3.76 | 4.55 | 120.94 |
| LINC01638 | 4.13 | 2.76 | 66.88 | 2.79 | 2.92 | 104.71 |
| PPP1R13L | 4.12 | 6.01 | 145.86 | 5.00 | 4.08 | 81.56 |
| TNFRSF12A | 4.12 | 3.63 | 88.13 | 4.88 | 6.17 | 126.61 |
| AFP | 4.12 | 2.52 | 61.14 | 2.60 | 2.79 | 107.45 |
| CENPN | 4.11 | 2.85 | 69.28 | 3.27 | 2.44 | 74.72 |
| SNAR-B2 | 4.11 | 2.21 | 53.73 | 3.97 | 2.98 | 75.03 |
| NFKBIZ | 4.10 | 2.22 | 54.19 | 3.23 | 3.35 | 103.60 |
| LAMP3 | 4.07 | 3.45 | 84.90 | 8.29 | 3.54 | 42.65 |
| ADAM12 | 4.06 | 2.03 | 50.07 | 3.10 | 3.13 | 100.79 |
| EPRS | 4.06 | 1.78 | 43.79 | 2.90 | 3.41 | 117.72 |
| SLC39A14 | 4.05 | 2.67 | 65.93 | 2.57 | 3.39 | 131.78 |
| LMO7 | 4.05 | 2.49 | 61.46 | 5.88 | 5.58 | 94.79 |
| CLEC18B | 4.05 | 1.42 | 34.98 | 3.78 | 1.61 | 42.61 |
| AK4 | 4.03 | 1.96 | 48.50 | 2.37 | 3.07 | 129.66 |
| CLTCL1 | 4.03 | 2.61 | 64.84 | 2.84 | 2.21 | 77.81 |
| QPCT | 4.02 | 1.51 | 37.59 | 3.51 | -0.33 | 0.00 |
| HHAT | 4.02 | 2.29 | 56.95 | 4.63 | -0.43 | 0.00 |
| CMKLR1 | 4.01 | 3.51 | 87.55 | 5.51 | -0.19 | 0.00 |
| GLI1 | 4.01 | 1.93 | 48.22 | 2.66 | -0.40 | 0.00 |
| DHX58 | -3.99 | -2.30 | 57.72 | -3.54 | -3.07 | 0.00 |
| ALDH3A2 | -3.99 | -2.12 | 53.18 | -5.24 | -3.76 | 71.73 |
| HLA-DMB | -4.00 | -3.40 | 84.97 | -5.31 | -3.73 | 70.24 |
| TYMSOS | -4.01 | -2.46 | 61.47 | -4.43 | -3.72 | 83.99 |
| PCDH18 | -4.02 | -2.08 | 51.81 | -3.33 | -3.21 | 96.31 |
| ZNRF1 | -4.02 | -1.83 | 45.58 | -3.59 | -3.30 | 91.92 |
| CD36 | -4.02 | -4.27 | 106.09 | -7.47 | -2.72 | 36.43 |
| C1S | -4.03 | -1.10 | 27.25 | -2.53 | -2.75 | 108.91 |
| PGM5P4-AS1 | -4.03 | -1.65 | 41.05 | -2.63 | -3.94 | 149.98 |
| LTBP4 | -4.03 | -1.53 | 37.87 | -3.06 | -4.40 | 143.44 |
| HDAC5 | -4.04 | -2.26 | 55.96 | -3.21 | -3.41 | 106.36 |
| TLE1 | -4.04 | -1.88 | 46.40 | -3.28 | -2.96 | 90.32 |
| VAT1L | -4.05 | -2.96 | 73.09 | -3.05 | -2.88 | 94.56 |
| AHNAK2 | -4.05 | -2.23 | 54.96 | -3.88 | -6.34 | 163.34 |
| P2RX7 | -4.05 | -1.99 | 49.11 | -2.47 | -2.43 | 98.39 |
| LINC01315 | -4.06 | -2.44 | 60.06 | -6.08 | -2.92 | 48.05 |
| PPARGC1A | -4.06 | -4.99 | 123.05 | -6.07 | -4.47 | 73.62 |
| SERPING1 | -4.08 | -1.43 | 34.98 | -2.47 | -2.70 | 109.21 |
| CYP3A7 | -4.08 | -3.01 | 73.77 | -4.48 | -2.55 | 57.04 |
| VEGFD | -4.08 | -5.31 | 130.02 | -4.26 | -3.09 | 72.51 |
| SGSH | -4.09 | -1.83 | 44.80 | -3.89 | -3.39 | 87.09 |
| ACKR4 | -4.09 | -2.02 | 49.34 | -4.22 | -1.48 | 35.08 |
| ADRB2 | -4.11 | -2.02 | 49.05 | -3.44 | -3.11 | 90.32 |
| RGS2 | -4.12 | -3.18 | 77.13 | -3.75 | -1.82 | 48.57 |
| SERPING1 | -4.13 | -1.14 | 27.50 | -2.24 | -2.12 | 94.85 |
| TNXB | -4.13 | -1.77 | 42.85 | -4.52 | -3.19 | 70.56 |
| FBLN1 | -4.13 | -1.65 | 39.99 | -4.82 | -3.40 | 70.53 |
| COL4A6 | -4.13 | -1.89 | 45.63 | -4.46 | -3.24 | 72.74 |
| MAST4 | -4.14 | -2.17 | 52.35 | -3.55 | -3.31 | 93.29 |
| AQP3 | -4.14 | -2.82 | 68.17 | -5.39 | -3.68 | 68.22 |
| FAM43A | -4.15 | -2.11 | 50.79 | -2.55 | -3.40 | 133.59 |
| COL13A1 | -4.15 | -2.24 | 53.85 | -3.95 | -4.13 | 104.61 |
| AKR1C3 | -4.16 | -2.19 | 52.67 | -4.54 | -3.31 | 72.75 |
| MX1 | -4.16 | -1.03 | 24.88 | -2.65 | -1.74 | 65.73 |
| TFPI | -4.16 | -2.25 | 53.98 | -3.97 | -2.59 | 65.18 |
| RASSF5 | -4.17 | -2.45 | 58.71 | -3.89 | -3.96 | 101.84 |
| CEMIP | -4.18 | -1.50 | 35.95 | -3.56 | -1.61 | 45.27 |
| STAR | -4.18 | -3.10 | 74.32 | -4.38 | -4.56 | 104.00 |

| | | | | | | |
|---|---|---|---|---|---|---|
| CIT | -4.18 | -3.08 | 73.69 | -3.90 | -3.78 | 96.95 |
| CD302 | -4.18 | -1.92 | 45.79 | -5.47 | -2.88 | 52.68 |
| HLA-DMA | -4.19 | -1.18 | 28.30 | -3.73 | -4.29 | 115.02 |
| IFIT3 | -4.19 | -2.04 | 48.77 | -3.50 | -2.44 | 69.75 |
| BTN3A2 | -4.19 | -1.98 | 47.25 | -4.10 | -3.61 | 88.15 |
| HMOX1 | -4.20 | -1.41 | 33.63 | -4.12 | -2.56 | 62.20 |
| ACP5 | -4.21 | -2.28 | 54.13 | -4.66 | -2.81 | 60.27 |
| LYNX1 | -4.21 | -2.29 | 54.40 | -3.81 | -3.80 | 99.64 |
| ADSSL1 | -4.23 | -2.30 | 54.37 | -4.15 | -3.15 | 75.83 |
| ABCC6 | -4.24 | -3.54 | 83.65 | -6.40 | -4.19 | 65.44 |
| HACD4 | -4.24 | -2.85 | 67.37 | -4.06 | -3.73 | 91.89 |
| CDHR3 | -4.24 | -2.43 | 57.23 | -3.92 | -2.88 | 73.27 |
| TNS2 | -4.25 | -2.10 | 49.41 | -3.40 | -2.89 | 85.09 |
| IGFBP5 | -4.25 | -1.90 | 44.72 | -2.36 | -2.96 | 125.40 |
| PLA2G4C | -4.26 | -2.52 | 59.16 | -4.52 | -5.20 | 115.15 |
| SLC7A8 | -4.31 | -2.65 | 61.62 | -2.56 | -3.31 | 128.90 |
| KCTD12 | -4.32 | -4.49 | 104.04 | -3.82 | -4.49 | 117.32 |
| GCHFR | -4.32 | -2.61 | 60.47 | -2.76 | -2.57 | 93.27 |
| DHRS13 | -4.32 | -3.23 | 74.74 | -3.10 | -3.86 | 124.63 |
| SOD3 | -4.34 | -1.88 | 43.40 | -4.02 | -2.90 | 72.00 |
| SBSN | -4.34 | -2.62 | 60.27 | -4.41 | -2.32 | 52.65 |
| CU678501 | -4.35 | -1.92 | 44.11 | -4.02 | -3.54 | 88.20 |
| IMPA2 | -4.36 | -1.84 | 42.30 | -2.45 | -2.78 | 113.75 |
| AK5 | -4.36 | -2.25 | 51.65 | -4.00 | -3.77 | 94.13 |
| ANO3 | -4.40 | -2.17 | 49.22 | -4.07 | -2.59 | 63.78 |
| CDC25B | -4.40 | -2.15 | 48.92 | -4.10 | -3.91 | 95.52 |
| LOC101927809 | -4.41 | -2.50 | 56.75 | -3.29 | -3.02 | 91.78 |
| MT1X | -4.41 | -2.49 | 56.53 | -5.03 | -4.03 | 80.20 |
| FER1L4 | -4.44 | -2.54 | 57.33 | -3.80 | -4.19 | 110.14 |
| HLA-DMA | -4.44 | -2.02 | 45.42 | -4.87 | -4.39 | 90.01 |
| PCDH18 | -4.46 | -2.36 | 52.95 | -3.11 | -2.72 | 87.28 |
| RAC2 | -4.46 | -2.03 | 45.45 | -3.28 | -2.94 | 89.48 |
| LPAR1 | -4.46 | -2.11 | 47.25 | -2.98 | -3.01 | 100.83 |
| SERPINF1 | -4.47 | -1.54 | 34.35 | -3.93 | -2.67 | 67.93 |
| SLC27A3 | -4.49 | -2.76 | 61.53 | -3.47 | -5.42 | 156.41 |
| ITGB8 | -4.49 | -2.15 | 47.85 | -4.99 | -2.69 | 53.81 |
| THBD | -4.51 | -4.03 | 89.35 | -3.78 | -3.84 | 101.82 |
| CATSPERZ | -4.51 | -4.00 | 88.70 | -6.82 | -4.69 | 68.75 |
| LTBP4 | -4.57 | -1.86 | 40.66 | -3.67 | -3.42 | 93.26 |
| C16orf89 | -4.57 | -2.66 | 58.07 | -5.42 | -4.49 | 82.87 |
| GAPLINC | -4.58 | -3.67 | 80.30 | -5.09 | -3.68 | 72.29 |
| CCDC102B | -4.59 | -3.99 | 86.90 | -5.78 | -3.11 | 53.75 |
| TNXB | -4.63 | -2.20 | 47.41 | -6.57 | -3.94 | 59.90 |
| S1PR2 | -4.64 | -2.18 | 47.04 | -4.88 | -3.60 | 73.67 |
| ITPKB | -4.65 | -2.50 | 53.70 | -3.42 | -3.40 | 99.26 |
| UBA7 | -4.65 | -1.69 | 36.40 | -3.46 | -3.09 | 89.11 |
| H19 | -4.67 | -1.41 | 30.19 | -4.00 | -3.16 | 79.12 |
| CCR10 | -4.67 | -3.19 | 68.35 | -4.24 | -4.28 | 101.13 |
| BDKRB1 | -4.67 | -2.68 | 57.35 | -3.19 | -2.01 | 63.03 |
| IL6R | -4.68 | -2.41 | 51.43 | -7.17 | -4.62 | 64.44 |
| LSAMP | -4.68 | -1.89 | 40.40 | -3.58 | -2.12 | 59.37 |
| COL14A1 | -4.68 | -3.20 | 68.31 | -6.26 | -3.07 | 49.12 |
| PTN | -4.69 | -1.68 | 35.77 | -5.81 | -1.85 | 31.87 |
| FXYD1 | -4.70 | -1.76 | 37.41 | -3.44 | -3.16 | 91.81 |
| SHC3 | -4.70 | -1.92 | 40.73 | -4.88 | -2.18 | 44.77 |
| KLHL41 | -4.71 | -4.71 | 99.90 | -4.15 | -3.33 | 80.29 |
| lnc-GMDS-3 | -4.81 | -3.47 | 72.06 | -5.06 | -4.04 | 79.90 |
| TM4SF1 | -4.82 | -2.56 | 53.05 | -4.64 | -3.47 | 74.93 |

| | | | | | | |
|---|---|---|---|---|---|---|
| NR1H3 | -4.84 | -2.73 | 56.27 | -6.20 | -4.58 | 73.83 |
| PLEKHO2 | -4.87 | -1.91 | 39.29 | -3.86 | -4.01 | 103.68 |
| MBP | -4.89 | -1.88 | 38.49 | -4.71 | -2.93 | 62.09 |
| SIPA1L2 | -4.90 | -2.15 | 43.97 | -4.61 | -2.20 | 47.81 |
| CD302 | -4.90 | -2.29 | 46.81 | -5.88 | -2.32 | 39.45 |
| TBX2 | -4.91 | -2.34 | 47.73 | -4.17 | -3.31 | 79.29 |
| RAB7B | -4.91 | -2.92 | 59.47 | -4.82 | -4.54 | 94.16 |
| ZNF436-AS1 | -4.91 | -2.94 | 59.86 | -4.99 | -3.99 | 79.80 |
| lnc-MRGPRF-1 | -4.91 | -2.89 | 58.74 | -5.18 | -4.32 | 83.31 |
| GUCY1B1 | -4.92 | -5.10 | 103.63 | -4.54 | -2.62 | 57.75 |
| BAALC | -4.93 | -2.50 | 50.84 | -4.31 | -2.94 | 68.11 |
| SCN4B | -4.94 | -3.98 | 80.74 | -3.77 | -4.07 | 107.96 |
| CYP27A1 | -4.95 | -2.55 | 51.44 | -5.47 | -3.36 | 61.48 |
| C16orf74 | -4.97 | -3.67 | 73.89 | -4.42 | -3.84 | 87.05 |
| TNXB | -4.98 | -1.96 | 39.27 | -5.27 | -4.09 | 77.55 |
| LAMA4 | -4.99 | -3.00 | 60.24 | -3.88 | -3.70 | 95.27 |
| PDK4 | -4.99 | -6.37 | 127.74 | -5.35 | -2.62 | 48.89 |
| SLITRK6 | -5.00 | -5.26 | 105.24 | -6.17 | -7.93 | 128.44 |
| ANGPTL2 | -5.01 | -2.60 | 51.76 | -6.35 | -3.46 | 54.45 |
| TMTC1 | -5.02 | -0.71 | 14.12 | -3.35 | -1.97 | 58.81 |
| LAMA4 | -5.02 | -2.82 | 56.17 | -5.04 | -4.10 | 81.37 |
| SNED1 | -5.03 | -1.95 | 38.86 | -4.81 | -2.82 | 58.62 |
| BMP2 | -5.03 | -2.08 | 41.29 | -4.48 | -2.65 | 59.15 |
| PLEKHG4 | -5.03 | -1.90 | 37.78 | -3.45 | -3.19 | 92.33 |
| PPARG | -5.04 | -2.13 | 42.18 | -4.57 | -3.78 | 82.71 |
| IGFBP4 | -5.05 | -2.12 | 42.01 | -2.99 | -2.31 | 77.30 |
| PKDCC | -5.08 | -2.47 | 48.72 | -5.63 | -5.09 | 90.47 |
| KCNK2 | -5.09 | -2.35 | 46.17 | -4.76 | -2.40 | 50.57 |
| DPP4 | -5.12 | -2.12 | 41.33 | -6.18 | -1.66 | 26.92 |
| PKDCC | -5.14 | -2.59 | 50.45 | -4.85 | -4.33 | 89.17 |
| HS6ST1 | -5.14 | -3.21 | 62.36 | -5.85 | -4.25 | 72.68 |
| PYCARD | -5.16 | -2.97 | 57.54 | -3.70 | -3.53 | 95.38 |
| CASP1 | -5.16 | -2.89 | 55.92 | -5.53 | -2.64 | 47.70 |
| FENDRR | -5.19 | -2.11 | 40.78 | -4.41 | -3.17 | 71.86 |
| LCE2C | -5.19 | -4.87 | 93.99 | -13.39 | -3.07 | 22.93 |
| ZNF395 | -5.19 | -2.16 | 41.62 | -5.09 | -3.81 | 74.86 |
| CAMK2N1 | -5.19 | -2.39 | 46.09 | -3.99 | -3.65 | 91.46 |
| PRRT2 | -5.20 | -5.60 | 107.75 | -6.13 | -6.21 | 101.33 |
| HGF | -5.20 | -1.81 | 34.71 | -3.86 | -1.72 | 44.55 |
| QSOX1 | -5.21 | -1.30 | 25.01 | -3.88 | -2.99 | 77.07 |
| RNA5-8SN5 | -5.21 | -3.08 | 59.14 | -8.32 | -14.95 | 179.63 |
| FMO3 | -5.21 | -3.44 | 66.07 | -4.00 | -3.03 | 75.72 |
| C4B | -5.22 | -2.03 | 38.94 | -3.10 | -2.31 | 74.37 |
| NR2F1 | -5.24 | -3.33 | 63.58 | -5.57 | -4.54 | 81.59 |
| BMPER | -5.24 | -2.92 | 55.63 | -3.52 | -3.12 | 88.50 |
| FENDRR | -5.25 | -1.92 | 36.64 | -4.48 | -2.80 | 62.46 |
| CA11 | -5.26 | -2.39 | 45.42 | -3.55 | -3.30 | 93.05 |
| CHI3L2 | -5.27 | -2.94 | 55.72 | -4.97 | -3.26 | 65.51 |
| PDE7B | -5.29 | -4.02 | 75.98 | -4.67 | -3.11 | 66.56 |
| SPIRE2 | -5.30 | -2.65 | 50.01 | -4.93 | -4.21 | 85.49 |
| SLC15A3 | -5.31 | -1.95 | 36.66 | -3.70 | -2.92 | 78.85 |
| S100A4 | -5.32 | -2.75 | 51.75 | -3.89 | -3.33 | 85.55 |
| ISYNA1 | -5.35 | -2.15 | 40.13 | -3.27 | -2.46 | 75.19 |
| PSMB9 | -5.36 | -2.73 | 50.96 | -4.05 | -3.48 | 85.71 |
| IL33 | -5.41 | -3.16 | 58.37 | -6.88 | -1.88 | 27.35 |
| PLEKHG4 | -5.44 | -2.43 | 44.69 | -3.92 | -3.38 | 86.24 |
| CLDN23 | -5.44 | -3.47 | 63.79 | -6.36 | -5.02 | 79.00 |
| QSOX1 | -5.52 | -1.39 | 25.24 | -3.70 | -2.80 | 75.84 |

| | | | | | | |
|---|---|---|---|---|---|---|
| OSR2 | -5.55 | -4.71 | 84.91 | -4.86 | -7.07 | 145.49 |
| BATF2 | -5.55 | -2.47 | 44.42 | -5.58 | -3.65 | 65.33 |
| ANKRD33B | -5.63 | -3.27 | 58.14 | -7.53 | -5.29 | 70.26 |
| H19 | -5.63 | -1.43 | 25.30 | -4.86 | -3.57 | 73.33 |
| GAPLINC | -5.66 | -4.07 | 71.96 | -5.52 | -4.31 | 77.98 |
| BDKRB2 | -5.66 | -4.10 | 72.42 | -5.19 | -4.07 | 78.40 |
| AKAP12 | -5.68 | -2.15 | 37.80 | -3.32 | -2.00 | 60.16 |
| NEURL1B | -5.72 | -4.41 | 77.16 | -4.22 | -4.79 | 113.70 |
| DAPK2 | -5.73 | -1.97 | 34.42 | -2.26 | -2.21 | 97.77 |
| S1PR1 | -5.73 | -6.43 | 112.28 | -6.59 | -4.96 | 75.33 |
| TGFBR3 | -5.73 | -3.93 | 68.64 | -6.60 | -2.67 | 40.43 |
| OLFML2A | -5.79 | -2.62 | 45.23 | -3.90 | -3.57 | 91.42 |
| BFSP1 | -5.79 | -3.38 | 58.36 | -6.44 | -4.07 | 63.11 |
| TNFRSF14 | -5.87 | -3.09 | 52.59 | -5.74 | -4.66 | 81.28 |
| TMEM158 | -5.89 | -2.68 | 45.51 | -4.39 | -1.74 | 39.61 |
| ATOH8 | -5.90 | -3.24 | 54.88 | -7.66 | -3.94 | 51.37 |
| KRT19 | -5.90 | -2.84 | 48.19 | -5.56 | -4.06 | 73.05 |
| PHLDA1 | -5.90 | -3.04 | 51.44 | -3.49 | -3.42 | 98.20 |
| VWCE | -5.97 | -3.40 | 56.94 | -7.67 | -4.76 | 62.14 |
| RASL12 | -5.98 | -4.51 | 75.36 | -5.87 | -3.52 | 60.02 |
| AKAP12 | -6.18 | -1.42 | 22.99 | -3.37 | -1.96 | 58.10 |
| SOCS2 | -6.19 | -2.76 | 44.60 | -5.64 | -4.40 | 78.05 |
| GRB14 | -6.21 | -5.42 | 87.40 | -2.19 | -2.81 | 128.18 |
| SELENBP1 | -6.21 | -4.22 | 67.89 | -6.70 | -7.19 | 107.26 |
| CLEC3B | -6.22 | -3.02 | 48.59 | -4.66 | -3.14 | 67.35 |
| C9orf47 | -6.26 | -4.64 | 74.08 | -7.38 | -3.87 | 52.47 |
| FBLN1 | -6.28 | -2.21 | 35.22 | -6.27 | -2.78 | 44.40 |
| RTP4 | -6.31 | -2.80 | 44.46 | -5.41 | -3.99 | 73.74 |
| HGF | -6.33 | -2.83 | 44.81 | -5.75 | -2.63 | 45.73 |
| IFIT1 | -6.33 | -2.95 | 46.61 | -4.59 | -4.22 | 91.88 |
| DNM3 | -6.34 | -4.62 | 72.79 | -9.25 | -6.25 | 67.59 |
| LRRC20 | -6.35 | -3.30 | 51.96 | -4.89 | -4.85 | 99.06 |
| CARD16 | -6.42 | -2.13 | 33.25 | -4.99 | -2.57 | 51.55 |
| NR0B1 | -6.43 | -5.57 | 86.75 | -7.50 | -6.70 | 89.33 |
| FGF13 | -6.45 | -2.89 | 44.80 | -5.45 | -4.36 | 80.05 |
| PCOTH | -6.45 | -2.93 | 45.36 | -5.42 | -4.21 | 77.65 |
| FBLN1 | -6.46 | -1.60 | 24.81 | -6.00 | -3.28 | 54.67 |
| DCLK1 | -6.48 | -4.17 | 64.30 | -4.27 | -3.11 | 72.83 |
| GUCY1A1 | -6.50 | -6.04 | 92.95 | -7.81 | -5.62 | 71.88 |
| MME | -6.51 | -3.18 | 48.87 | -6.44 | -2.92 | 45.38 |
| P2RY1 | -6.53 | -3.04 | 46.53 | -9.48 | -3.94 | 41.54 |
| PCOTH | -6.55 | -2.86 | 43.66 | -6.46 | -4.49 | 69.53 |
| SAMD5 | -6.57 | -4.21 | 64.17 | -4.23 | -4.19 | 99.06 |
| CCDC102B | -6.62 | -5.38 | 81.25 | -8.12 | -4.64 | 57.07 |
| NEDD4L | -6.63 | -3.17 | 47.87 | -5.48 | -4.03 | 73.43 |
| CARD17 | -6.66 | -1.94 | 29.06 | -4.85 | -2.43 | 50.03 |
| GRK5 | -6.69 | -2.98 | 44.54 | -7.72 | -4.68 | 60.64 |
| CASP17P | -6.70 | -2.87 | 42.85 | -7.03 | -3.27 | 46.48 |
| AQP3 | -6.74 | -6.01 | 89.20 | -8.67 | -5.31 | 61.27 |
| MAOA | -6.78 | -2.07 | 30.55 | -5.42 | -3.85 | 71.03 |
| SPRY1 | -6.91 | -1.58 | 22.82 | -4.30 | -3.21 | 74.72 |
| APOC1 | -6.92 | -2.99 | 43.27 | -6.21 | -4.53 | 73.00 |
| ENST00000528781 | -6.93 | -1.85 | 26.68 | -3.54 | -3.64 | 103.00 |
| PDE5A | -6.93 | -3.66 | 52.89 | -4.62 | -3.10 | 66.99 |
| ZFP36L2 | -6.98 | -3.17 | 45.47 | -7.28 | -3.66 | 50.31 |
| COLEC12 | -6.99 | -3.89 | 55.56 | -8.25 | -2.94 | 35.62 |
| EPAS1 | -7.16 | -2.32 | 32.45 | -5.62 | -4.00 | 71.16 |
| KIT | -7.16 | -1.97 | 27.59 | -2.52 | -1.85 | 73.62 |

| | | | | | | |
|---|---|---|---|---|---|---|
| ACVRL1 | -7.19 | -3.04 | 42.34 | -2.81 | -3.66 | 130.36 |
| ATP1A2 | -7.19 | -3.34 | 46.49 | -4.31 | -3.47 | 80.47 |
| OLFML1 | -7.22 | -2.50 | 34.64 | -4.19 | -4.08 | 97.40 |
| PTGIR | -7.36 | -2.04 | 27.67 | -3.42 | -1.99 | 58.29 |
| RASL11A | -7.38 | -5.73 | 77.66 | -8.14 | -4.37 | 53.63 |
| ALDH1A3 | -7.41 | -4.40 | 59.41 | -13.50 | -3.88 | 28.72 |
| CEMIP | -7.45 | -1.39 | 18.60 | -8.54 | -1.57 | 18.38 |
| CCBE1 | -7.54 | -2.23 | 29.61 | -7.22 | -5.14 | 71.18 |
| APCDD1 | -7.54 | -3.10 | 41.10 | -6.08 | -5.28 | 86.79 |
| FAXDC2 | -7.60 | -4.31 | 56.66 | -5.99 | -5.37 | 89.54 |
| PALMD | -7.66 | -3.42 | 44.59 | -6.85 | -4.87 | 71.03 |
| CCL2 | -7.76 | -0.92 | 11.87 | -5.95 | -2.18 | 36.54 |
| MYOC | -7.76 | -3.52 | 45.42 | -6.19 | -6.67 | 107.71 |
| LAMA4 | -7.79 | -2.82 | 36.16 | -7.22 | -4.82 | 66.76 |
| KITLG | -7.82 | -3.00 | 38.33 | -6.48 | -3.00 | 46.25 |
| ST8SIA1 | -7.90 | -4.59 | 58.15 | -8.26 | -6.09 | 73.65 |
| SVEP1 | -7.94 | -1.43 | 17.96 | -4.47 | -2.46 | 55.03 |
| LINC00484 | -8.08 | -6.04 | 74.73 | -7.21 | -6.41 | 88.90 |
| MME | -8.18 | -4.33 | 52.91 | -8.03 | -3.63 | 45.21 |
| NTN1 | -8.25 | -2.89 | 35.04 | -7.61 | -4.01 | 52.62 |
| RIPOR3 | -8.27 | -3.65 | 44.22 | -4.27 | -4.61 | 108.12 |
| CXCL12 | -8.30 | -1.73 | 20.79 | -9.20 | -4.16 | 45.17 |
| SLC9A9 | -8.36 | -5.47 | 65.38 | -7.32 | -6.36 | 86.86 |
| AMPD3 | -8.40 | -3.35 | 39.85 | -5.54 | -3.45 | 62.31 |
| IMPA2 | -8.42 | -2.26 | 26.85 | -4.51 | -4.28 | 94.94 |
| MOXD1 | -8.44 | -1.95 | 23.10 | -6.71 | -3.27 | 48.77 |
| GDF5 | -8.57 | -1.35 | 15.78 | -3.66 | -2.91 | 79.54 |
| IFITM1 | -8.78 | -1.82 | 20.73 | -5.83 | -1.87 | 32.00 |
| TBX2-AS1 | -8.81 | -3.97 | 45.11 | -6.90 | -4.97 | 72.11 |
| TCF21 | -8.91 | -5.15 | 57.80 | -9.79 | -8.82 | 90.08 |
| QPRT | -8.92 | -4.17 | 46.70 | -6.73 | -5.79 | 86.08 |
| APOL6 | -8.92 | -5.06 | 56.74 | -2.80 | -2.10 | 75.21 |
| TBX2-AS1 | -9.05 | -4.02 | 44.47 | -7.14 | -5.01 | 70.23 |
| DENND2A | -9.13 | -3.82 | 41.86 | -7.17 | -3.93 | 54.72 |
| METTL7A | -9.18 | -3.55 | 38.69 | -6.60 | -4.68 | 70.80 |
| GALNT15 | -9.26 | -3.40 | 36.75 | -4.83 | -3.49 | 72.13 |
| HSD11B1 | -9.30 | -2.82 | 30.29 | -10.23 | -3.99 | 39.01 |
| ACPP | -9.33 | -1.90 | 20.38 | -5.52 | -3.58 | 64.88 |
| TMTC1 | -9.33 | -2.35 | 25.22 | -5.34 | -2.93 | 54.86 |
| HTR2B | -9.40 | -2.01 | 21.39 | -3.84 | -1.83 | 47.53 |
| SFRP1 | -9.43 | -3.25 | 34.43 | -8.06 | -4.34 | 53.91 |
| CASP17P | -9.45 | -2.81 | 29.74 | -8.41 | -3.18 | 37.81 |
| MME | -9.47 | -3.41 | 35.96 | -7.84 | -3.15 | 40.17 |
| SMAD3 | -9.49 | -2.70 | 28.42 | -6.05 | -3.58 | 59.07 |
| CCN3 | -9.54 | -2.62 | 27.43 | -5.19 | -3.99 | 76.86 |
| HSPB3 | -9.64 | -2.38 | 24.71 | -6.40 | -2.73 | 42.67 |
| NR4A3 | -9.72 | -4.45 | 45.80 | -12.57 | -4.47 | 35.60 |
| IFITM1 | -9.92 | -1.83 | 18.46 | -6.86 | -2.26 | 32.95 |
| KRT32 | -9.95 | -2.87 | 28.85 | -6.73 | -4.87 | 72.34 |
| RGCC | -10.21 | -3.69 | 36.11 | -4.65 | -3.51 | 75.54 |
| PLEKHA6 | -10.29 | -3.88 | 37.76 | -9.24 | -5.01 | 54.26 |
| KCNJ2 | -10.32 | -3.87 | 37.56 | -8.59 | -3.64 | 42.41 |
| LCE2A | -10.41 | -6.14 | 59.00 | -28.27 | -4.48 | 15.86 |
| KRTAP1-5 | -10.49 | -1.89 | 18.01 | -6.80 | -1.89 | 27.74 |
| LRRN4CL | -10.59 | -2.68 | 25.34 | -7.05 | -6.06 | 85.94 |
| SPRY1 | -10.70 | -2.18 | 20.36 | -7.26 | -4.41 | 60.70 |
| MAN1C1 | -11.00 | -3.05 | 27.70 | -7.78 | -4.15 | 53.37 |
| S1PR3 | -11.17 | -3.86 | 34.59 | -9.45 | -5.93 | 62.74 |

| | | | | | | |
|---|---|---|---|---|---|---|
| ANGPTL4 | -11.89 | -1.50 | 12.66 | -4.28 | -1.97 | 46.09 |
| GPX3 | -12.16 | -2.61 | 21.49 | -11.57 | -4.50 | 38.94 |
| LOC102724458 | -12.55 | -2.45 | 19.54 | -10.74 | -3.85 | 35.88 |
| PPL | -12.61 | -1.80 | 14.31 | -5.67 | -6.58 | 116.04 |
| ASPA | -12.67 | -4.76 | 37.58 | -10.56 | -5.80 | 54.96 |
| SYNE3 | -12.95 | -4.16 | 32.16 | -8.17 | -7.23 | 88.46 |
| PLIN4 | -13.00 | -3.45 | 26.56 | -16.30 | -9.20 | 56.45 |
| LSP1 | -13.52 | -5.12 | 37.88 | -7.28 | -6.20 | 85.13 |
| GPX3 | -13.70 | -2.68 | 19.53 | -10.27 | -4.92 | 47.91 |
| SHISA3 | -13.89 | -8.04 | 57.92 | -16.90 | -9.13 | 54.00 |
| LINC02154 | -14.07 | -3.45 | 24.50 | -7.49 | -4.86 | 64.90 |
| VSIR | -14.20 | -4.25 | 29.90 | -7.67 | -8.84 | 115.30 |
| FGL2 | -14.56 | -6.37 | 43.74 | -17.82 | -2.93 | 16.46 |
| PLPP3 | -14.66 | -4.75 | 32.41 | -11.48 | -5.58 | 48.62 |
| TMEM119 | -15.06 | -3.45 | 22.94 | -16.32 | -7.36 | 45.11 |
| SLC40A1 | -15.81 | -9.40 | 59.48 | -13.05 | -8.32 | 63.75 |
| RARRES3 | -16.76 | -4.10 | 24.49 | -10.44 | -5.07 | 48.56 |
| ADAMTS8 | -16.97 | -0.97 | 5.70 | -5.00 | -2.00 | 39.90 |
| A2M | -17.72 | -2.23 | 12.59 | -12.52 | -6.56 | 52.42 |
| SECTM1 | -18.30 | -4.51 | 24.67 | -18.30 | -6.98 | 38.11 |
| ALDH1A1 | -18.45 | -2.65 | 14.35 | -11.89 | -3.01 | 25.35 |
| SELENOP | -19.14 | -5.48 | 28.64 | -14.91 | -5.55 | 37.19 |
| TSGA10IP | -19.38 | -6.73 | 34.75 | -14.33 | -14.35 | 100.10 |
| SELENOP | -19.41 | -6.60 | 34.01 | -16.79 | -5.84 | 34.78 |
| ADH1B | -20.05 | -7.89 | 39.34 | -25.55 | -15.60 | 61.06 |
| TNXB | -20.13 | -3.28 | 16.28 | -20.56 | -5.35 | 26.03 |
| TNXB | -21.44 | -2.68 | 12.49 | -21.07 | -6.22 | 29.51 |
| SOCS1 | -24.81 | -10.60 | 42.71 | -18.63 | -12.67 | 68.01 |
| CLDN11 | -26.38 | -3.48 | 13.17 | -18.07 | -11.83 | 65.50 |
| FOXQ1 | -27.96 | -9.44 | 33.78 | -17.14 | -12.11 | 70.65 |
| NPTX1 | -34.26 | -16.29 | 47.56 | -36.43 | -14.79 | 40.61 |
| FMO2 | -42.81 | -7.63 | 17.82 | -27.39 | -3.88 | 14.17 |
| ENST00000584094 | -43.29 | -8.25 | 19.06 | -35.74 | -10.24 | 28.64 |
| ADM | -60.64 | -3.17 | 5.22 | -19.16 | -6.63 | 34.60 |
| ADH1A | -163.06 | -7.29 | 4.47 | -73.21 | -21.03 | 28.73 |
| ADH1C | -246.54 | -7.06 | 2.86 | -96.87 | -21.31 | 22.00 |

**Supplemental Table S3:PCR primer sequences**

| Gene | *Forward* | *Reverse* |
|---|---|---|
| *hUBC* | GTGGTGCGTCCAGAGAGAC | GGCCTTCGCCATATCCTTTTC |
| *hPRRX1a* | AGCGTCTCCGTACAGATCCT | GTAGCCATGGCGCTTTTCAG |
| *hPRRX1b* | TCCGAGACCCACCGATTATCT | AAGTAGCCATGGCGCTGTA |
| *hACTA2* | GAAGAGCATCCCACCCTGC | ATTTTCTCCCGGTTGGCCT |
| *hCOL1A1* | GCCAAGACGAAGACATCCCA | GTTTCCACACGTCTCGGTCA |
| *hFN1* | AGCAAGCCCGGTTGTTATGA | CCCACTCGGTAAGTGTTCCC |
| *hCCNA2* | CATGTCACCGTTCCTCCTTG | CCAATGGTTTTCTGGGTCCA |
| *hCCNE2* | TGGCCACCTGTATTATCTGGG | TCCCCAGCTTAAATCAGGCA |
| *hACTG2* | ATGTACGTCGCCATTCAAGC | TCTCTCTCAGCTGTGGTCAC |
| *hTAGLN* | GTATGACGAGGAGCTGGAGG | TCAGGGTACAGGCTGTTCAC |
| *hPPM1A* | TGCATGTGATGGTATCTGGGA | GCTTCTGGCGATACTTTGGG |
| *hTGFBR2* | ATGCTGCTTCTCCAAAGTGC | GCTGATGCCTGTCACTTGAA |
| *mRna18S* | CTTAGAGGGACAAGTGGCG | ACGCTGAGCCAGTCAGTGTA |
| *mPrrx1a* | CTCTCCGTACAGCGCCAT | GTTGGCCATGTTGATACCCT |
| *mPrrx1b* | CCGTACAGATCTTCGTCCCT | TTCCTCAGTTGACTGTTGGC |
| *mActa2* | AGTCGCTGTCAGGAACCCTGAGA | ATTGTCGCACACCAGGGCTGTG |
| *mCol1a1* | GTGTGTGACAAGGGTGAGACA | GAGAACCAGGAGAACCAGGA |
| *mFn1* | TGGTGGCCACTAAATACGAA | GGAGGGCTAACATTCTCCAG |
| *mCol14a1* | GTTCAACGTGGGCTCAGAAA | ACTCCTCGATCCTGCTTCTG |
| *mMki67* | AACAAGAGTGAGGGAATGCC | GCTGTGAGTGCCAAGAGACT |
| *hCCNA2* promoter | TTTAACACGGAGCTCACATAGT | CAGTAGTTCAAGGTGCCATCTTA |
| *hCCNE2* promoter | CGTGACGCCGGCAAAATAAT | AGCGTTAGAAATGGCAGAAAGT |
| *hMKI67* promoter | CCGCAACATTAGCAAATCGATTT | CGTCACTTTTCCTTGGTGCT |
| *hPPM1A* promoter | TGCGAATGTGGTGTAGGTCA | CGCCGGGGATAGAATGACAA |

#### Supplementary Table S4: Antibody list

| *Antibody* | *Concentration* | *Application* |
|---|---|---|
| GAPDH | Mouse monoclonal, Covalab, Villeurbanne, France (00006513) | Western Blot |
| β-TUBULIN | Rabbit polyclonal, Abcam, Cambridge, USA (ab6046) | Western Blot |
| β-ACTIN | Mouse monoclonal clone AC-74 (A2228), Sigma, Saint-Louis, USA | Western Blot |
| PRRX1 | Mouse monoclonal, clone 1E2, Sigma, Saint-Louis, USA (SAB1412737) | Western Blot |
| | Rabbit polyclonal, Sigma, Saint-Louis, USA **(**HPA051084**)** | Immunochemistry; immunofluorescence and chromatin Immunoprecipitation |
| COL-I | Goat polyclonal ,Southern Biotech, Birmingham, USA ( 1310-01) | Western Blot |
| | Rabbit polyclonal, Abcam, Cambridge, UK (ab34710) | Immunochemistry |
| FN-1 | Rabbit polyclonal, Abcam, Cambridge, UK (ab2413) | Western Blot |
| ACTA2 | Mouse monoclonal, clone 1A4, Sigma, Saint-Louis, USA (A5228) | Western Blot and immunochemistry |
| Phospho(S423+S425) SMAD3 | Rabbit monoclonal, Abcam, Cambridge, USA (ab52903) | Western Blot |
| SMAD3 | Rabbit polyclonal, Abcam, Cambridge, UK (ab28379) | Western Blot |
| Phospho(S255)-SMAD2 | Rabbit monoclonal, Abcam, Cambridge, USA (ab188334) | Western Blot |
| SMAD2 | Rabbit monoclonal, Cell signalling Technology, Danves, USA (5339S) | Western Blot |
| TGFBR2 | Rabbit polyclonal, Cell signalling Technology, Danves, USA (79424) | Western Blot |
| PPM1A | Rabbit polyclonal, Sigma, Saint-Louis, USA (HPA029209) | Western Blot |
| KI67 | Rabbit monoclonal, clone RM227, Sigma, Saint-Louis, USA (SAB5600050) | Immunochemistry |
| Vimentin | Rabbit monoclonal, Abcam, Cambridge, USA (ab92547) | Immunochemistry |
| CD45 | Mouse Monoclonal, Dako Agilent Technologies, Les Ulis France (M0701) | Immunochemistry |
